# A systems pharmacology approach to determine the mechanisms of action of pleiotropic natural products in breast cancer from transcriptome data

**DOI:** 10.1101/2020.04.18.048454

**Authors:** Regan Odongo, Asuman Demiroğlu Zergeroğlu, Tunahan Çakir

## Abstract

Plant-derived compounds as natural products have attracted a lot of attention in the treatment of complex diseases, especially cancers, primarily due to their poly-pharmacologic mechanisms of action. However, methodological limitations have impeded gaining complete knowledge of their molecular targets. While most of the current understanding of these compounds is based on reductive methods, it is increasingly becoming clear that holistic techniques, leveraging current improvements in omic data collection and bioinformatics methods, are better suited for elucidating their systemic effects. Here, to provide an explanation to the mechanisms of action of plant-derived natural products in breast cancer, we applied a data integration approach to comprehensively study oncogenic signaling pathways targeted by withaferin A, actein, compound kushen injection and indole-3-carbinol. Specifically, we mapped the transcriptome-level response of cancer cell lines to these molecules on a human protein-protein interaction network and constructed the underlying active subnetworks. We used these subnetworks to define the perturbed signaling pathways and validated their relevance in carcinogenesis. The similarity of each identified oncogenic signaling pathway in terms of overlapping genes was subsequently used to construct pathway-pathway interaction networks, which were used to reduce pathway redundancy and to identify pathway crosstalk. Filtered pathways were then mapped on three major carcinogenesis processes. The results showed that the pleiotropic effects of plant-derived drugs at the gene expression level can be used to predict targeted pathways. Thus, from such pathways, it is possible to infer a systemic mechanism of action of such natural products.

## Introduction

While reductionist-based approaches generated much of the drugs and drug targets known today, drug-human interactions are rather complex since the mechanism of action of most pharmacologically effective drugs results from the perturbation of multi-dimensional cellular networks^1^. Thus, a phenotypic change following a treatment is the result of regulation cascades covering various biomolecular interactions, which can be traced in omics scale^1,2^. Within this scope, several studies have utilized transcriptomic data to generate novel hypotheses from drug perturbations in various diseases. In order to decipher meaningful information from such high-throughput perturbation data, novel computational approaches in the context of systems biology need to be applied^2^.

Most cancers are driven by multiple genetic mutations and epigenetic dysregulations^3,4^ interconnected by biomolecular players. Breast cancer is the most prevalent form of cancer in women. Distinct subtypes have been defined for this cancer, and inter-group subtle genetic variations are known to exist. Owing to the understanding of the existence of somatic mutations that aggregate in a few signaling and regulatory pathways^5^, a number of small molecule targeted therapies have been developed for breast cancer in the last decade. However, treatment success rates above 40% are yet to be recorded^6^. A plausible explanation is the inherent oncogenic signaling pathway cross-talks and the bypass of targets by alternative activating pathways. This explicitly points to a need for multi-targeted therapeutic approaches.

Experimental evidences from separate molecular biology studies on the use of plant-based drugs in cancer cells have strongly suggested a multi-targeting therapeutic strategy. In fact, ancient civilizations relied on plant-based drugs due to their low systemic toxicities and ability to simultaneously ameliorate multiple disease symptoms^7^. Justifiably, current systems biology analyses through differential gene expression enumerations have confirmed similar observations. Yet, despite their observed anti-cancer effects, no attempt has been made to integrate transcriptome-level response to these drugs with molecular interaction networks to systemically evaluate the mechanism of action of these drugs. Emboldened by the idea that co-regulated and co-expressed biomolecules tend to converge on well-defined biological pathways, we hypothesised that genes targeted by plant-based drugs form unique subnetworks; enriched with oncogenic signaling pathways critical in regulating information flow in response to drug treatment. To test such a hypothesis, we aimed to catalogue all the molecular players in a perturbed subnetwork module and use the resulting observations to devise an approach for elucidating the mechanism of action of plant-based molecular compounds.

Network biology is a holistic approach in systems biology to understand biological systems, where biomolecules and their binary interactions are projected onto a graph to depict molecular relationships^8,9^. Nowadays, concurrent integration of experimentally-derived omics data with a priori interaction data is a common approach in systems biology to obtain context-specific subnetworks^10^. To this end, a number of computational tools have been proposed by different groups to map transcriptome data on protein-protein interaction networks^11^. These tools have been applied to several diseases so far, including breast cancer^12^, hepatocellular carcinoma^13,14^, liver fibrosis^15^ and neurodegenerative diseases^16,17^.

In this study, we mapped transcriptome data of breast cancer cell lines treated with plant-derived drugs onto protein-protein interaction networks and applied a subnetwork discovery algorithm to extract condition-specific subnetworks. Subsequently, we applied bioinformatic pathway enrichment methods to gain an understanding of the underlying biological processes. Specifically, we studied actein^18^, compound kushen injection (CKI)^19^, indole-3-carbinol^20^ and Withaferin A^21^ treated breast cancer cell lines. Next, we investigated the prognostic roles of the topologically significant genes in each subnetwork. We also defined an approach to decipher the mechanism of action of each drug from a network perspective, based on the construction of pathway-pathway interactions and mapping oncogenic signalling pathways to major carcinogenesis processes. Overall, we showed that these pharmacognostic products possess pleiotropic properties and their effects are reverberated through well-established oncogenic signaling pathways. Notably, multiple perturbed oncogenic signaling pathways coordinate to control common carcinogenesis processes. We anticipate that the approach proposed here will be instrumental in accelerating drug evaluation of poly-pharmacologic lead compounds of plant origin for applications in oncology precision medicine and other complex diseases.

### Methodology

The computational analysis steps utilized in this study are summarized in **Figure 1**.

**Figure 1:**
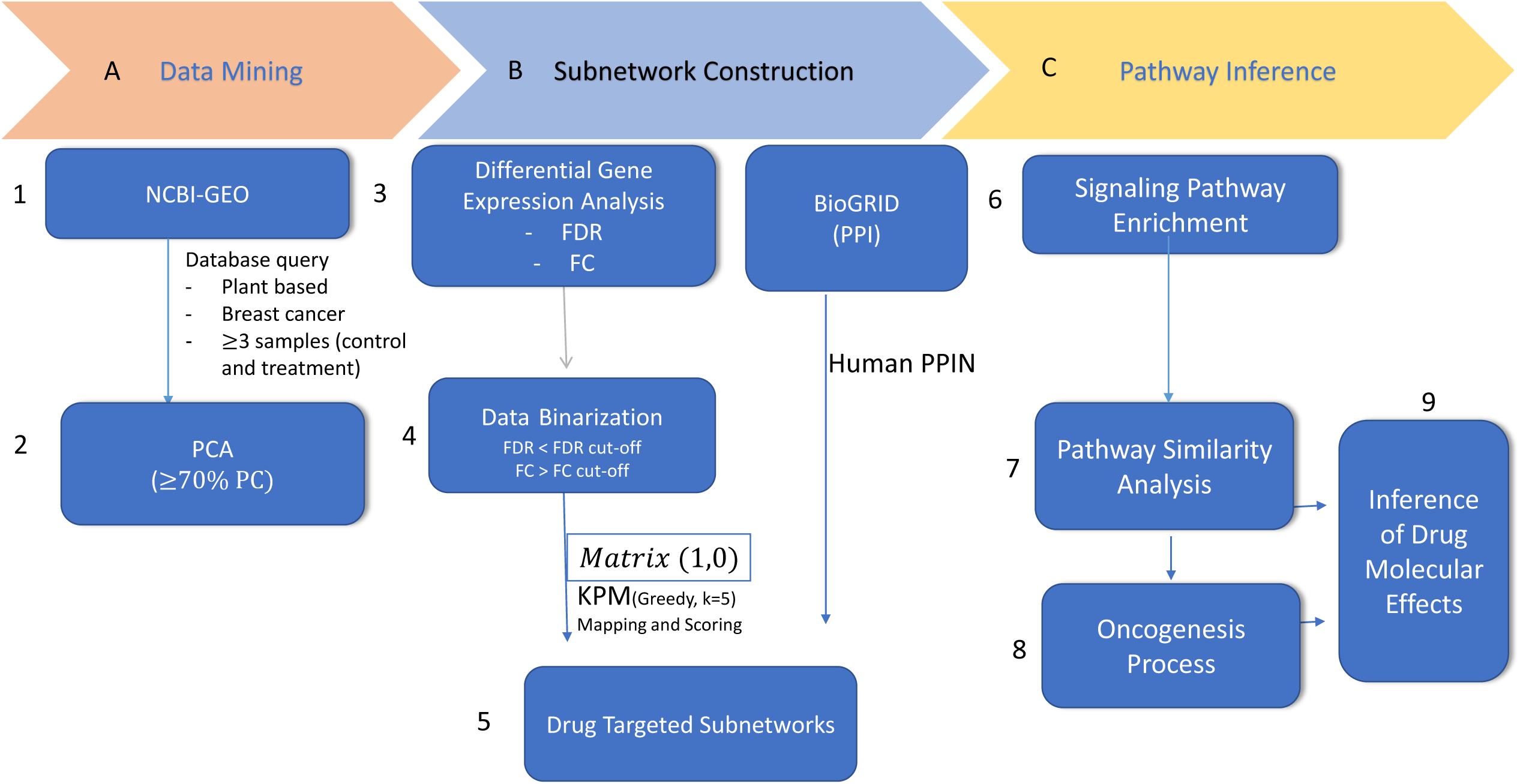
Computational analysis workflow applied in this study. The approach is centred on three main analysis sections: data mining, subnetwork discovery and pathway inference. PCA: Principal component analysis, FDR: False discovery rate, FC: Fold change, KPM: KeyPathwayMiner

### Data acquisition

We used a structured query statement to interrogate and download gene expression datasets for the breast cancer cell lines treated with withaferin A (GSE53049)^21^, actein (GSE7848)^18^, CKI (GSE78512)^19^ and indole-3-carbinol (GSE55897)^20^ from the NCBI GEO depository. We selected these four plant-based drugs among others since the corresponding datasets had at least 3 control and 3 treatment groups, and there was a distinct separation between the control and treatment groups (tested using the unsupervised dimension reduction method, principal component analysis).

### Data processing and differential gene expression analysis

The expression datasets included microarray expression profiles and RNA-seq counts and, therefore, platform specific protocols were followed. For the microarray derived datasets (withaferin A, actein and indole-3-carbinol), probeset mapping was performed by choosing the probe with the maximum average expression value among multiple probesets of a gene. For RNA-seq data (CKI), we selected only those genes with above zero counts in at least two samples in either control or treatment group. Overall, we log2 normalized all the pre-processed datasets. Subsequently, we used LIMMA^22^ package in R to identify differentially expressed genes between the treated versus control (untreated) groups. We used Benjamini-Hochberg p-value correction to control false discovery rates (FDR). Fold change and FDR cut-offs were simultaneously used to select differentially expressed genes.

### Active subnetwork scoring and construction using KeyPathwayMiner

The challenge of discovering most-connected drug specific subnetworks in the human protein-protein interaction network was solved using KeyPathwayMiner (KPM)^23^, one of the tools reported to have a high performance among subnetwork discovery methods^11^. In this approach, given a priori protein-protein interaction network (PPIN), we were interested in a maximally connected clique based on a significance score. Hence, we treat this problem as an optimization problem with two main constraints: (i) the maximum allowable non-differentially expressed genes, and (ii) the significance cut-off. In this work, we used the Cytoscape (v3.7.1) based KPM (v5.0.1) plugin.

In our analysis, we made a few modifications to the input data and constraints as we describe next. We applied a uniform fold-change cut-off of 2 and a varied FDR cut-off of 5X 10^−3^ (for indole-3-carbinol and withaferin A) or 1X 10^™2^ (for actein and CKI) to identify differentially expressed genes. Thus, our approach is a little stricter; with the intent to further limit the rate of false positives. These two cut-offs were used to assign binary values to all the genes in a dataset. Specifically, we used ‘1’ to denote differentially expressed genes based on our criteria, and ‘0’ for other genes. In the subnetwork construction, significantly changed and physically interacting proteins are used. These interconnected proteins essentially denote drug-targeted cellular pathways. We allowed a maximum of 5 non-differentially expressed genes in each subnetwork solution, a parameter available in KPM. For the priori human PPIN, we used BioGRID^24^ (release 3.5.173; 25^th^ March, 2019) containing 22,435 proteins and 478,529 interactions.

### Subnetwork analysis and prospective validation of high centrality genes

Using CytoNCA (v2.1.6)^25^ Cytoscape plugin, we analysed two network topological features to identify the major genes in the subnetworks: degree and betweenness centrality. Next, we used the TCGA breast cancer RNA-Seq data to investigate the prognostic values of the top 5 (based on high degree and betweenness centrality) identified genes. Specifically, we used the online tool KM-Express^26^ to determine the effect of the identified genes on overall survival and their association with samples from normal, primary and metastatic cases. For the overall survival, the tool uses the median gene expression across all samples and a hazard ratio to infer statistical significance based on log-rank p-value. A p-value cut-off of 0.05 was used in this study.

### Pathway enrichment analysis

We used enrichR^27^ package in R to perform pathway enrichment analysis for the respective subnetwork nodes (genes). It takes pathway definitions from Kyoto Encyclopaedia of Genes and Genomes (KEGG), WikiPathway, Reactome and Gene Ontology Biological Process (GO-BP) databases, among others. We limited our results to the enriched pathways with an FDR cut-off of 0.05 and containing the terms: ‘signal’, ‘apoptosis’, and ‘cell cycle’. Also, those pathways with less than 3 associated genes were removed at this step.

### Construction of pathway-pathway interaction network

Oncogenic signaling pathways do not function in isolation but are known to crosstalk with each other while redirecting cellular processes. Construction of pathway interaction networks has been previously applied to visually elaborate the pathway-pathway interrelationships and infer associated biological phenomenon^28,29^. On the other hand, since pathway enrichment via enrichR was based on multiple pathway databases, redundant pathways were inevitable in the enrichment results. Therefore, pathway-pathway similarity can also be used to identify redundant pathways. One approach to computationally enumerate such relationships is to evaluate the degree of pathway-pathway overlap based on gene similarities in any given two pathways. We used the Jaccard index; which is a measure of the similarity between a pair of sets. Here, given two pathways, *P*_*i*_ and *P*_*j*_, with enriched gene sets, *G*_*i*_ and *G*_*j*_, we computed the Jaccard index (*J*) using the formula below:

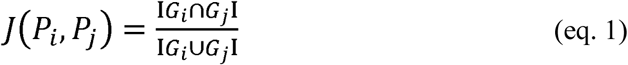

This evaluates to the number of genes common in the two pathways divided by the total number of genes in both pathways without repeats. Hence, Jaccard index takes values between 0 and 1, and, using this metric, the proportional similarity between two pathways can be deduced. Here, we defined two pathways to be either in crosstalk or similar based on their Jaccard scores. We relied on a cut-off of 0.60 and 0.25 to infer pathway redundancy and pathway crosstalk respectively. Since we used multiple pathway databases (KEGG, GO-BP, WikiPathways and Reactome pathway definitions) in our analysis, which increased the possibility of pathway redundancies, this approach allowed us to prioritize a family representative for redundant pathways, effectively eliminating sub-pathways originating from the same pathway database. To graphically illustrate the outcome of the Jaccard analysis and visually inspect the pathways for prioritization, we used the igraph R package^30^ to construct pathway-pathway interaction network as we describe later. The pathway definitions were used as the network nodes while a cut-off of 0.25 was used to insert an edge between any pathways with at least 25% common genes. Furthermore, we used greedy optimization algorithm in igraph to define clusters in a pathway-pathway interaction network.

### Oncogenic signaling pathway inference

Using the pathway-pathway interaction networks, we applied a two-tier approach to infer biological significance. First, we relied on the 10 canonical oncogenic signaling pathways from the comprehensive pathway analysis by the TCGA Pan-Cancer Consortia^31^, which are cell cycle, Hippo, Myc, Notch, NRF2, PI-3-Kinase/Akt, RTK-RAS-MAPK, TGF-beta P53 and β-catenin/Wnt signalling pathways. Among the terms identified in our enrichment analysis, we selected the terms that were semantically related to the aforementioned canonical pathways as drug-targeted signaling pathways. Subsequently, we grouped such terms into three broad clusters depicting the main cancer pathophysiologic processes: (i) cell cycle, proliferation and apoptosis, (ii) cell metastasis and invasion, and (iii) angiogenesis^32^.

## Results

### Construction of drug responsive protein interaction subnetworks from transcriptome data

Breast cancer is molecularly classified into three main subtypes: luminal (A and B), triple negative and human epidermal receptor 2 positive (HER2+); based on hormone receptor and HER2 expression^33^. While the datasets used in this study included representative cell lines from the three subtypes, they differ on the transcriptomic platforms used to collect the data and the drug applied. Nevertheless, we believe that the approach applied here captures the systemic drug effects and is sufficient to study the pleiotropic nature of plant derived drugs. We summarise these datasets in **Supplementary Table 1**. In general, our datasets include luminal A (T47D, MCF-7, ZR751), triple negative (MDA-MB-231, MDA-MB-157 and MDA-MB-436) and human epidermal receptor 2 positive (MDA-MB-468) breast cancer cell lines treated with at least one of indole-3-carbinol, Withaferin A, CKI and Actein. The Principal Component Analysis analysis results showing separate grouping of treatment and control samples is available as **Supplementary Figure 1**. To identify drug affected genes, we performed differential gene expression analysis. We relied on fold change and FDR scores as cut-offs for significance; which were eventually used for data binarization for KPM analysis, as described in the Methods section. Corresponding numbers of differentially expressed genes are given in **Supplementary Table 2**.

Network mapping and subnetwork scoring approaches have been extensively used in integrative biology field to discover active disease- and drug-specific modules in various experiments^11,23,34,35^. To elucidate the molecular effects of plant derived drugs in breast cancer, we constructed the active subnetworks from transcriptome data using KeyPathwayMiner^34^. Concurrently, using the same approach and parameters, we also constructed active subnetworks from the up- and down-regulated genes separately. The number of proteins and their interactions for all the subnetworks solutions are reported in **Table 1**.

**Table 1:** Summary of topological structure of subnetwork solutions indicating the number of proteins and their interactions in each dataset studied. CKI: Compound kushen injection, I3C: Indole-3-carbinol and WA: Withaferin A

| Drugs | Cell Lines | Genes | Interactions |  | Genes | Interactions |
| --- | --- | --- | --- | --- | --- | --- |
| Actein | MDA-MB-453 | 829 | 3858 | Up | 327 | 687 |
|  |  |  |  | Down | 455 | 2166 |
| CKI | MCF-7 | 1332 | 9331 | Up | 933 | 2838 |
|  |  |  |  | Down | 304 | 1676 |
| I3C | MCF-7 | 1974 | 10684 | Up | 453 | 1162 |
|  |  |  |  | Down | 1399 | 6816 |
|  | T47D | 1681 | 7050 | Up | 620 | 1324 |
|  |  |  |  | Down | 959 | 3254 |
|  | ZR751 | 1403 | 5457 | Up | 545 | 1105 |
|  |  |  |  | Down | 961 | 6323 |
|  | MDA-MB-231 | 93 | 126 | Up | 17 | 17 |
|  |  |  |  | Down | 86 | 111 |
|  | MDA-MB-157 | 86 | 110 | Up | 18 | 19 |
|  |  |  |  | Down | 75 | 106 |
|  | MDA-MB-436 | 541 | 1275 | Up | 98 | 120 |
|  |  |  |  | Down | 402 | 932 |
| WA | MCF-7 | 333 | 941 | Up | 117 | 353 |
|  |  |  |  | Down | 202 | 564 |
|  | MDA-MB-231 | 998 | 3277 | Up | 456 | 1011 |
|  |  |  |  | Down | 480 | 1208 |

Overall, we observed a compound- and breast cancer subtype-specific number of proteins and their interactions. Thus, it is deducible that the different drugs studied had substantial differential effects on the activity of the underlying protein interaction networks in the disease conditions. With the differences in the number of targeted proteins, this deduction reinforces the dominant idea that no two drugs have a similar mechanism of action in complex diseases^2,36^. As expected, the role of molecular heterogeneity of the different breast cancer subtypes in drug response can be explicitly delineated from the sizes of the subnetworks. For instance, under indole-3-carbinol, in terms of the number of enriched genes, a relatively higher number was targeted by LA than TN, while the reverse was observed under Withaferin A treatment of LA and TN cell types (**Table 1**). The current drug research regime focusses on targeted therapy (famously defined as ‘magic bullets’)^2,36^. However, with the increasing acceptance of the poly-pharmacologic paradigm as an effective alternative in the treatment of complex diseases, our network analysis results indicate that the analysed plant-derived natural products target multiple proteins simultaneously to exert their effects and is a testament to a multi-targeting mechanism of action by the individual drugs. This observation would be beneficial under disease conditions, particularly if the cohort of targeted proteins can be linked to or are known disease drivers.

### The drug-specific subnetworks capture key breast cancer carcinogenesis-related genes as revealed by prospective prognostic prediction using network topology analysis

An overarching question is whether the genes enriched in the subnetwork solutions have any significance in breast cancer prognosis. In therapeutic terms, effective anti-carcinogenic drug candidates are known to regulate a niche of known proto-oncogenes in a disease network. To address this, network centrality measures can be used to identify topologically important target vertices (genes) in the subnetwork solutions^9,25^. In disease networks under compound perturbations, such genes are significantly enriched as a result of the condition (treatment) change. In this study, with the aim to prospectively validate the constructed subnetworks, we used CytoNCA^25^ to extract the top five genes based on both high betweenness and degree centralities from each subnetwork. The result from this analysis is reported in **Table 2.** Betweenness and degree centrality scores of all genes in the subnetworks are given in **Supplementary Table 3.** Subsequently, we analysed the top-five genes by using the KM-Express^26^ tool for their association with overall survival and for their relationship with pathological stages (median expression in normal, tumor and metastasis states).

**Table 2:**
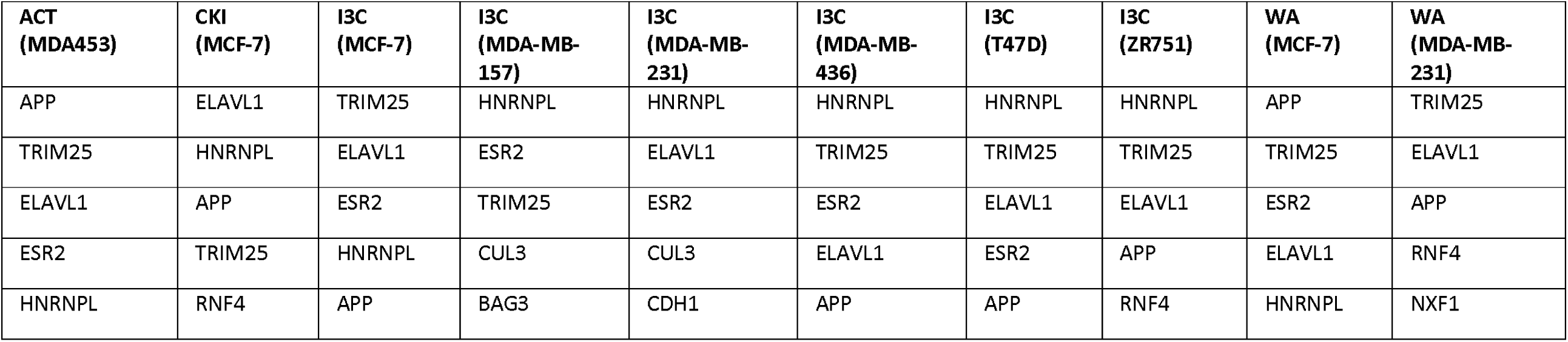
Top 5 genes from the subnetworks for each dataset based on their betweenness and degree centrality scores. The genes are labelled using their respective universal identifiers. ACT: Actein, CKI: Compound kushen injection, I3C: Indole-3-carbinol, and WA: Withaferin A

| ACT<br>(MDA453) | CKI<br>(MCF-7) | I3C<br>(MCF-7) | I3C<br>(MDA-MB-157) | I3C<br>(MDA-MB-231) | I3C<br>(MDA-MB-436) | I3C<br>(T47D) | I3C<br>(ZR751) | WA<br>(MCF-7) | WA<br>(MDA-MB-231) |
| --- | --- | --- | --- | --- | --- | --- | --- | --- | --- |
| APP | ELAVL1 | TRIM25 | HNRNPL | HNRNPL | HNRNPL | HNRNPL | HNRNPL | APP | TRIM25 |
| TRIM25 | HNRNPL | ELAVL1 | ESR2 | ELAVL1 | TRIM25 | TRIM25 | TRIM25 | TRIM25 | ELAVL1 |
| ELAVL1 | APP | ESR2 | TRIM25 | ESR2 | ESR2 | ELAVL1 | ELAVL1 | ESR2 | APP |
| ESR2 | TRIM25 | HNRNPL | CUL3 | CUL3 | ELAVL1 | ESR2 | APP | ELAVL1 | RNF4 |
| HNRNPL | RNF4 | APP | BAG3 | CDH1 | APP | APP | RNF4 | HNRNPL | NXF1 |

In general, we found 11 unique genes from all the subnetworks. Five of these genes (APP, TRIM25, ELAVL1, HNRNPL and ESR2) were found to be the most frequent across all subnetworks (**Table 2**). Since we had allowed the parameter *K* = 5 in KPM-based subnetwork extraction, top five genes mainly consisted of non-significantly expressed but highly connected genes in response to treatment. Coincidentally, they had the highest betweenness centrality scores as well. Survival analysis found APP, TRIM25 and ELAVL1 to have significant associations with overall survival (log-rank p-value< 0.05) in breast cancer. Overexpression of APP and TRIM25 in cancer patients was associated with low overall survival and the reverse was true for ELAVL1 (**Supplementary Figure 2a-c**). In the literature, APP is a well-established cancer biomarker, a target of ADAM10, and has been strongly linked with breast cancer growth, metastasis and migration^37^. A comprehensive study identified TRIM25 as a key gene in regulating TN breast cancer metastasis^38^. ELAVL1 codes for an RNA binding protein controlling multiple facets of carcinogenesis, and literature reports show its over-expression to be associated with adverse-event free tumors^39^. Indeed, our current finding concurs that its low expression in cancer patients correlates with low overall survival and that over-expression may increase the patient overall survival. On the other hand, HNRNPL and ESR2, which have been reported to be associated with breast cancer elsewhere^40,41^, were not significantly associated with patient survival at the median gene expression cut-off. However, further interrogation revealed their significant association with overall survival at 75% vs 25% (high vs low) and 75% gene expression cut-offs respectively (**Supplementary Figure 2d-e**). From **Supplementary Figure 2f-j**, high expression levels of TRIM25 is associated with metastatic tumors while that of ELAVL1 is associated with primary tumors. The expression of APP, on the other hand, decreases in both primary and metastatic tumors., We found TRIM25 to be indirectly targeted by all the compounds, except in MDA-MB-231 under indole-3-carbinol (Figure 2). Also, under indole-3-carbinol treatment, APP was not present amongst the top-five genes in MDA-MB-231 and MDA-MB-157, indicating a transcriptome deviation from the other TNBC-specific cell line, MDA-MB-436.

**Figure 2:**
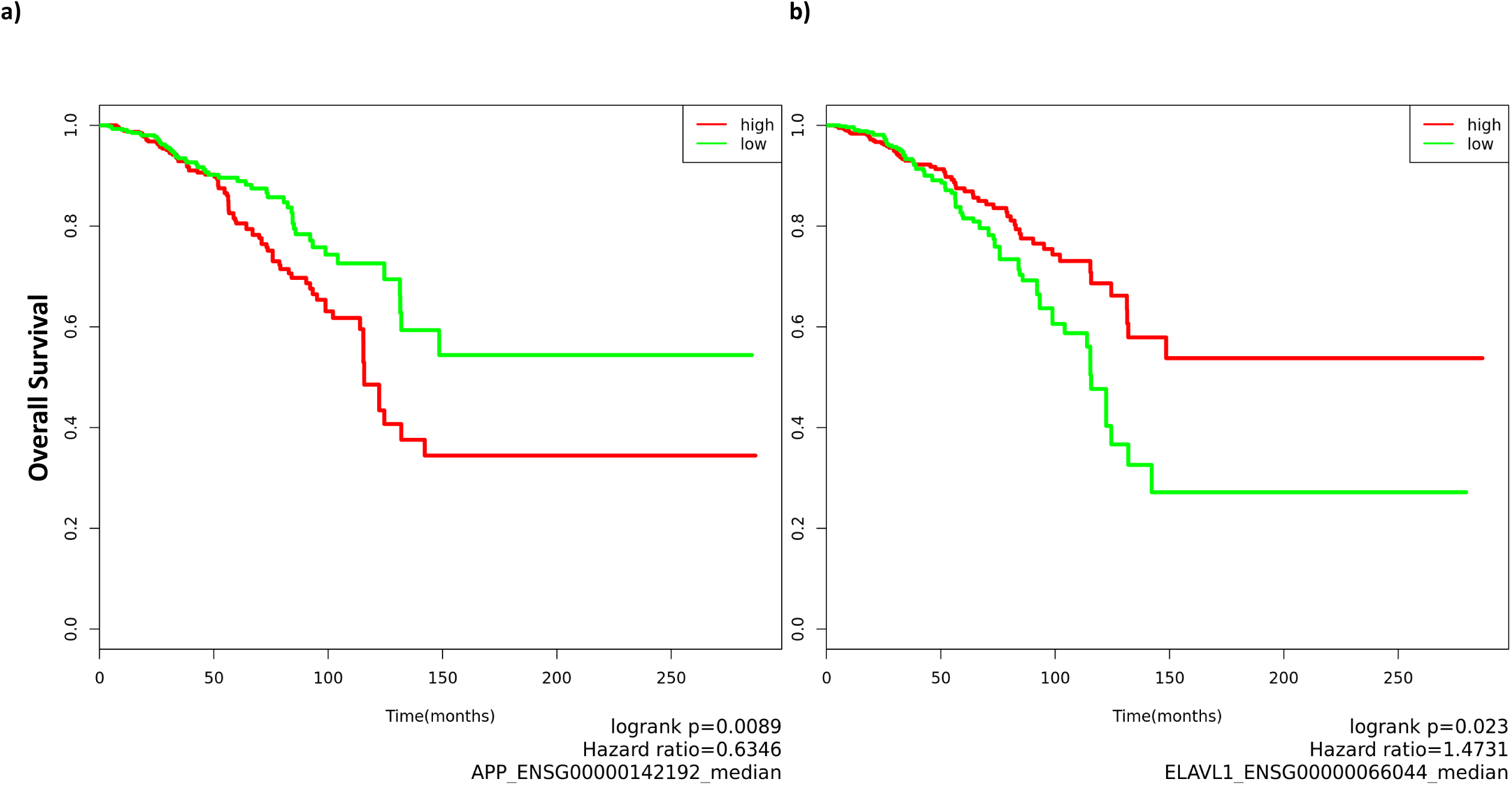
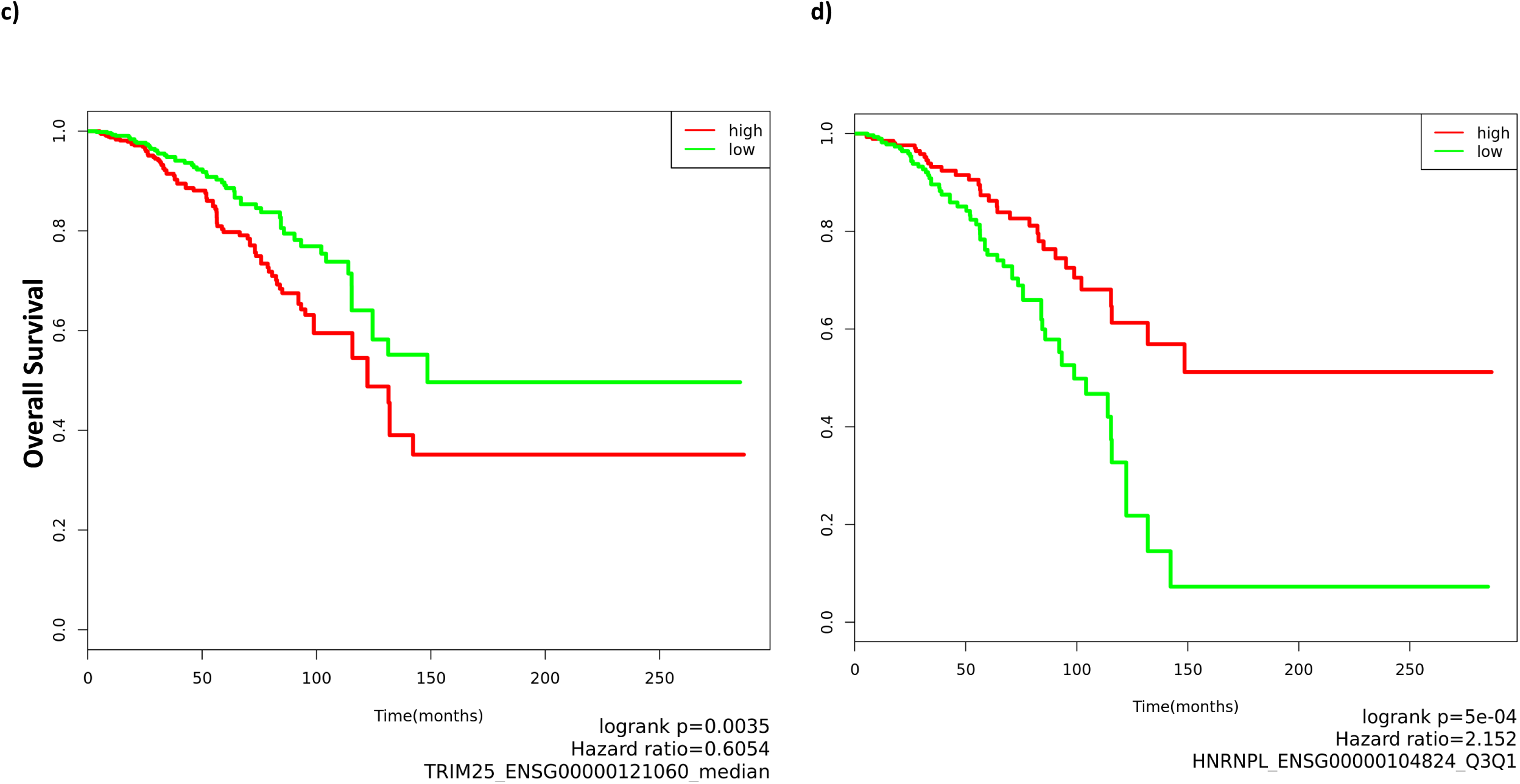
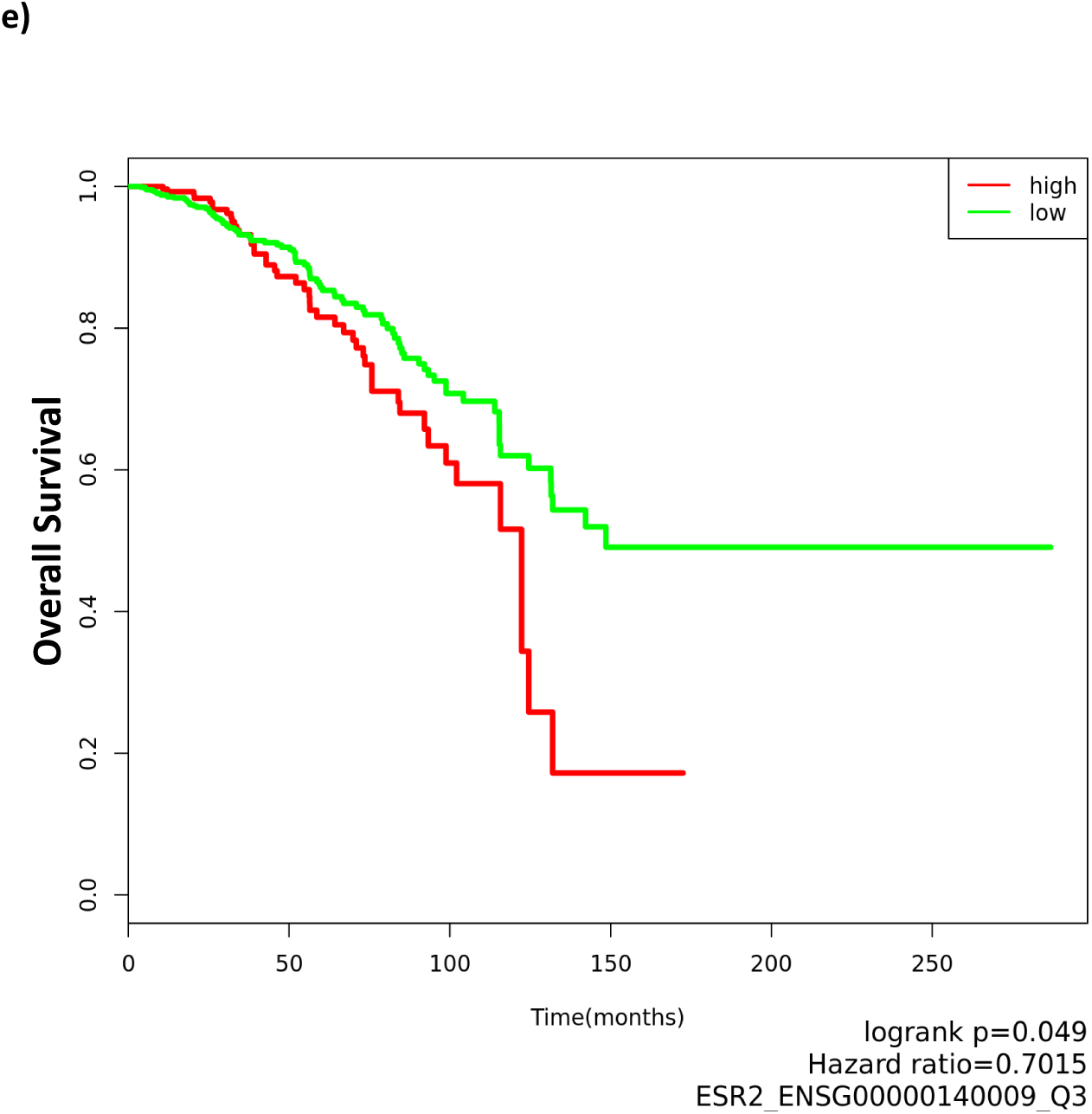
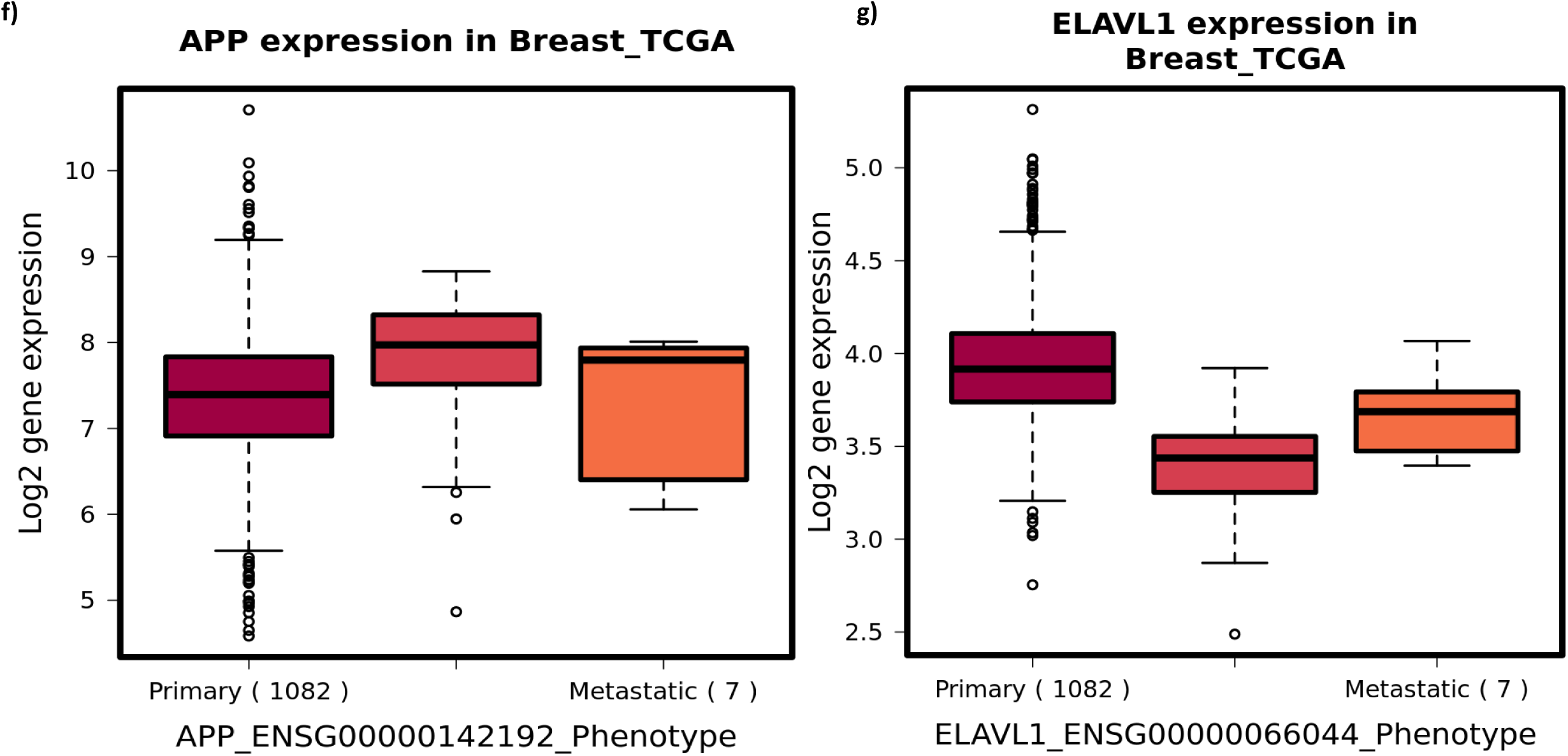
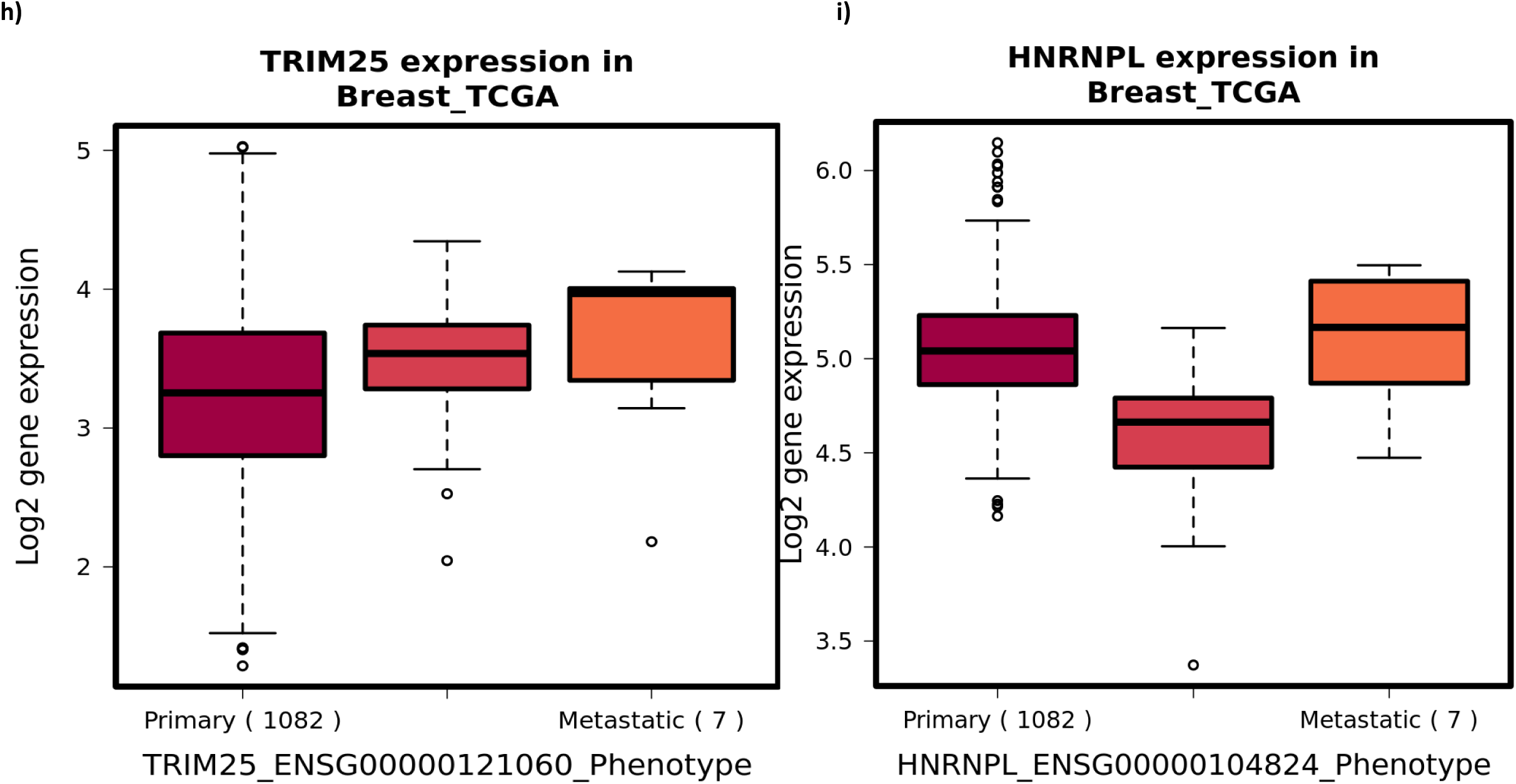
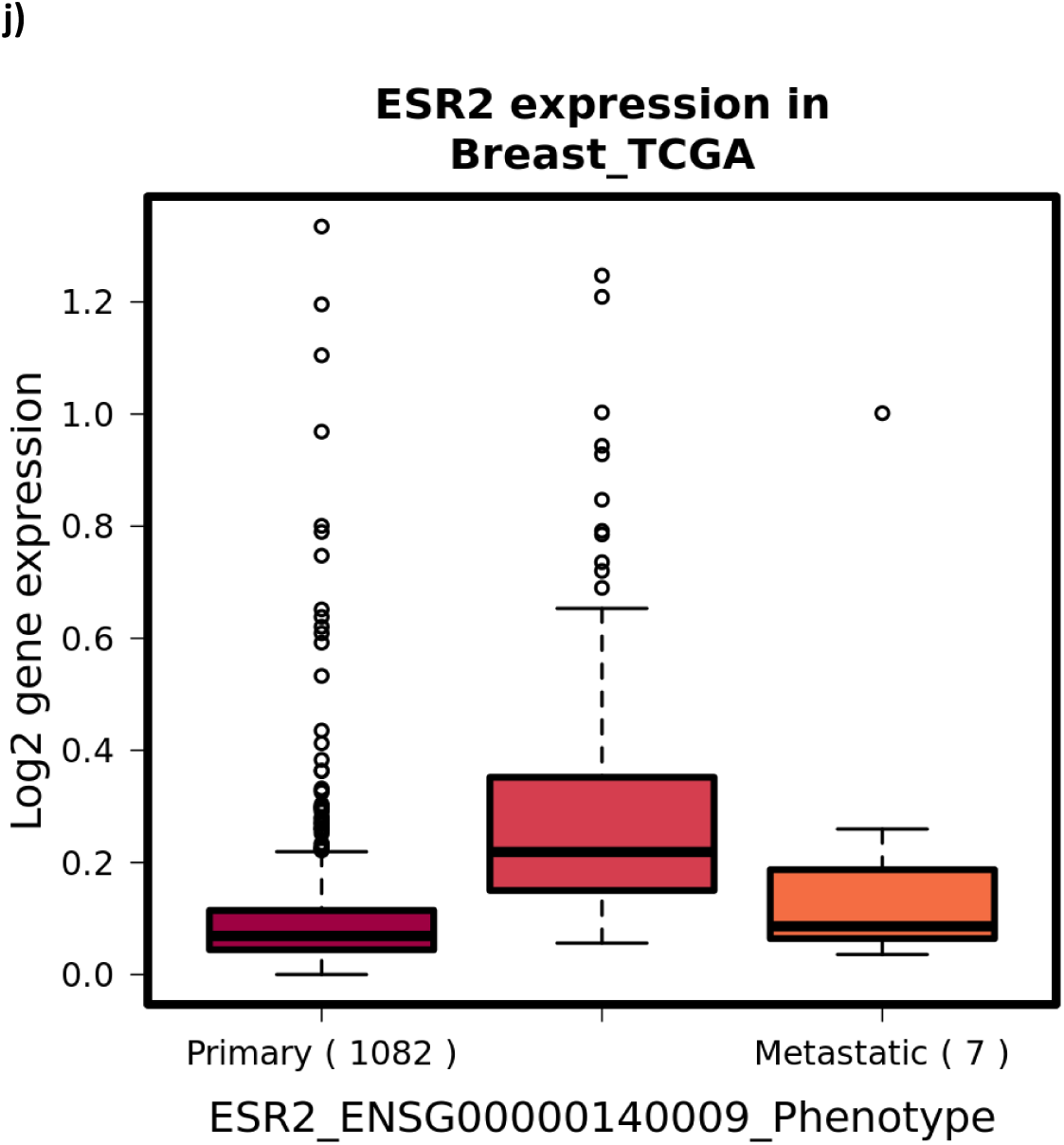
Pathway-pathway interaction networks under Actein (MDA-MB-453 cell line) and Withaferin A (MDA-MB-231 cell line) treatments. The network nodes represent individual pathways while the coloured clusters represent both pathway crosstalk and similarity. Pathway-pathway crosstalk (Jaccard Index) ≥0.25.

These findings indicate that these plant-derived compounds target gene subnetworks driven by well-established oncogenes. Importantly, the plant-based compounds exert their effects not directly through the central oncogenes but by perturbing a high number of their first neighbours to modify the underlying physiological conditions. This protein-disease-prognosis consistency is a validation of the efficiency of the applied method to capture biologically informative protein networks and shows the effectiveness of the compounds in cancer, permitting the constructed subnetworks as viable in hypothesis generation.

### Actein, indole-3-carbinol, CKI and Withaferin A target multiple oncogenic signaling pathways which coordinate to influence cellular processes

The current pharmacokinetics and pharmacodynamics studies are highly efficient in elucidating the mechanism of action of anti-microbial drugs. However, studies have consistently demonstrated that this simple framework is inefficient in addressing drug action in complex and multi-factorial disease systems. In such systems, limiting drug research to targeting single disease biomarkers is one of the main causes of drug failures in clinical trials ^1,2,42^. Drug induced reprogramming of cellular responses is directed through metabolic reactions, which are regulated by signaling pathways enormously enriched in protein-protein interactions. Thus, undeniably, studying drug effects on cellular pathways provides a holistic approach as to the molecular targets of drug candidates. Given the increased preference by tumors for only a handful number of such pathways, a sound anti-carcinogenic effect can thus be deduced by evaluating their activity upon treatment. A recent study evaluating oncogenesis related pathways based on gene profiling in various cancers^31^ provides a foundation for systemically evaluating the therapeutic relevance of drug-responsive pathways upon treatment in various tumors.

The pleiotropic nature of plant-derived drugs in cancer is well anchored in literature^7,43^. However, linking drug targeted networks from transcriptome data with oncogenesis processes to study the mechanism of action of natural products as a holistic approach has not been explored systematically. Thus, we reasoned that taking such an approach would present a novel method to studying the poly-pharmacologic compounds.

In this section, we aimed to comprehensively catalogue drug targeted oncogenic signaling pathways and their corresponding oncogenesis processes. In summary, the following procedure was followed: (i) pathway enrichment was applied to all the genes in a subnetwork, (ii) only oncogenic signaling pathways were retained, (iii) to identify and filter out redundant pathways coming from different databases, pathway-pathway correlation networks were constructed (iv) the final list of pathways were mapped on three major oncology related processes based on their semantic similarity to the 10 canonical oncogenic signalling pathways^31^ (see **Methods** section).

As described in the methods section, we performed pathway enrichment analysis using the genes in each identified subnetwork. An important factor in this systemic approach is the interconnectivity of the proteins used in pathway enrichment analysis. Thus, it is obvious that the enriched pathways are connected due to the shared targeted-network proteins. To illustrate this, first we eliminated all those pathways which were unrelated to cancer. **Supplementary Table 4 and Supplementary Table 5** report the enriched pathways from this analysis. Then we constructed unweighted pathway-pathway interaction networks based on common proteins shared between different pathways. We relied on a Jaccard similarity index of at least 25% to denote pathway crosstalk (through intersecting genes) and represented this by placing an edge between them in the network. **Figure 2a-b and Supplementary Figure 3a-g** shows the networks of various drug targeted pathways from the four drugs studied. This clustering allowed us to (i) prioritise meaningful signaling pathway terms for mapping on oncogenesis processes thus reducing redundancy (the pathways with J>0.60), and (ii) illustrate pathway-pathway crosstalk (interdependence) in a drug-targeted network. We reckon that this approach is much simpler and precise compared to Chen et al.^44^’s gene overlap index approach for pathway prioritisation.

We observed a characteristic clustering of related pathway terms across the various enrichment results. For instance, in the actein treated MDA-MB-453 dataset, we identified 10 pathway clusters out of 21 enriched pathways; only 5 of these (NRF2, Cell cycle, Apoptosis, Interferon signaling and TGF-beta) were identified as members of the defined oncogenic signaling pathways (see Methods). An examination of the various pathway clusters from all the datasets revealed two important features: (i) the clustered pathways were either semantically related or from the same database with similar functions, as is the case of ‘NRF2’ and ‘Nuclear receptor meta-pathway’ pathways in **Figure 2a** (J>0.60, pathway redundancy), and (ii) the interacting pathways are well-known to interact in literature acting as sub-pathways through the activation of the main pathway, as is the case of ‘apoptosis’, ‘TNF’ and ‘IL17’ in **Figure 2b** (pathway crosstalk), which is expected^45^. The pathway-pathway interaction networks from the other datasets are reported in **Supplementary Figure 3a-g**.

Next, to infer biological significance, we applied a two-tier approach. First, we relied on the predefined canonical oncogenic signaling pathways (see **Methods** section)^31^ for the concise terms. Additionally, though not captured in the TCGA^31^ analysis of the most frequently mutated canonical oncogenic signaling pathways since it is a response mechanism to foreign system, the role of the immune system signaling as a secondary response mechanism in cancer is significant and can be attributed to the inhibition/promotion of tumor initiation and metastasis in advanced cases. Thus, immune system related pathway terms were also included in the analysis results based on the known physiological roles of both the pathways and their enriched genes. Subsequently, we used pathway enrichment analysis results from the up-/down-regulated subnetworks (**Supplementary Table 5**) to assign these pathways as either up- or down-regulated. Eventually, with clear pathway clusters and only canonical-signaling-pathways relevant non-redundant terms, we mapped the resulting pathway terms on the three categories derived from major oncogenesis processes: (i) cell cycle, proliferation and apoptosis, (ii) cell metastasis and invasion, and (iii) angiogenesis. However, given the overlapping roles different pathways perform in biological systems, deciphering the affected processes is not straightforward. Therefore, to assign a pathway to either of the three groups, we looked up for the functional role(s) of the associated genes (both up- and down-regulated) in UniProtKB^46^ database. To deduce the targeted biological processes, we relied on those genes whose molecular functions match the biological roles of the pathways provided in literature. **Table 3** details the results of this grouping. To illustrate this approach, we provide a detailed description of the grouping as applied to the actein treated MDA-MB-453 cell line in **Supplementary Table 6** using enrichment results from Supplementary Table 5 and the pathway-pathway interaction networks (**Figure 2a, b** and **Supplementary Figure 3a-g**).

**Table 3:**
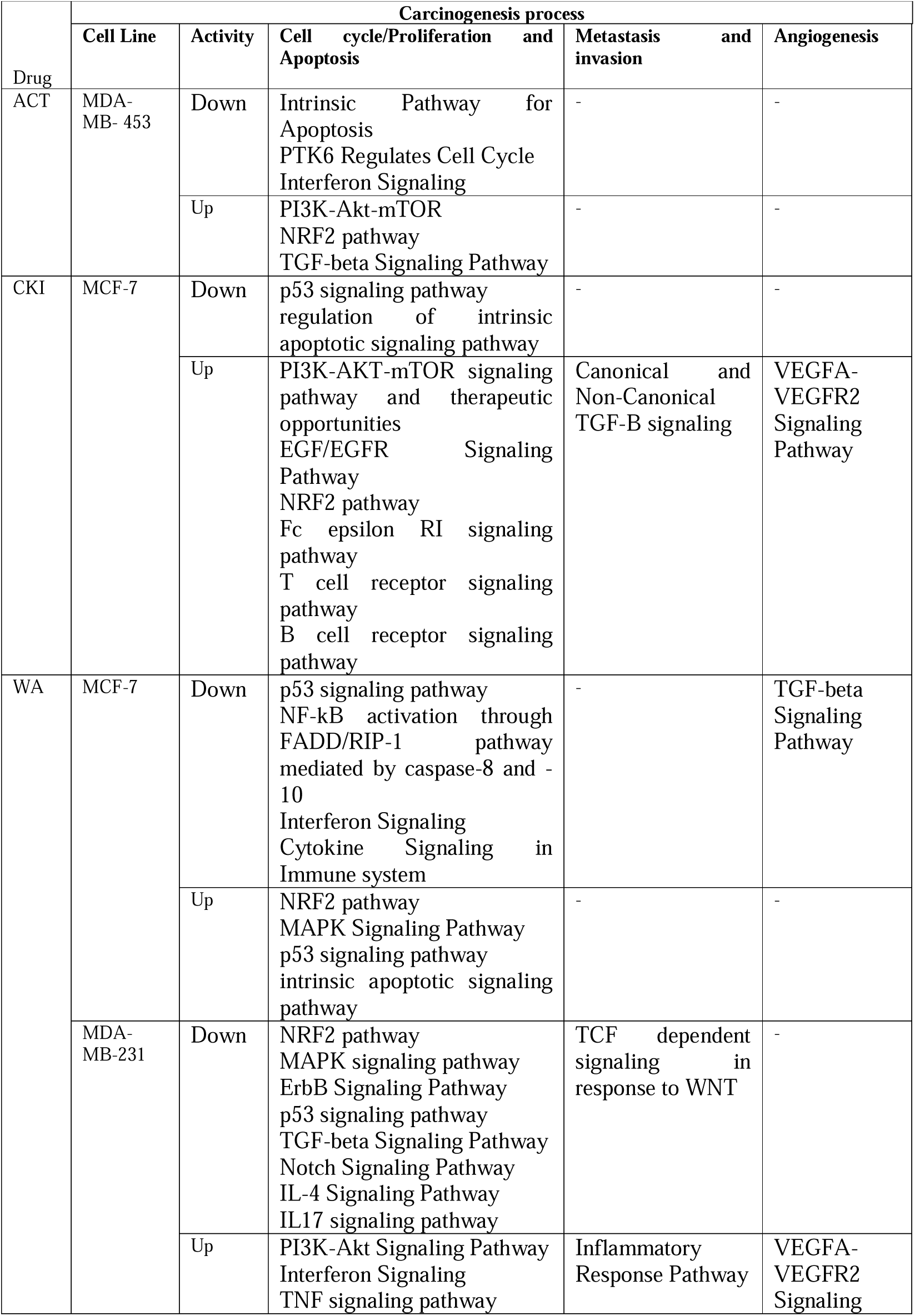
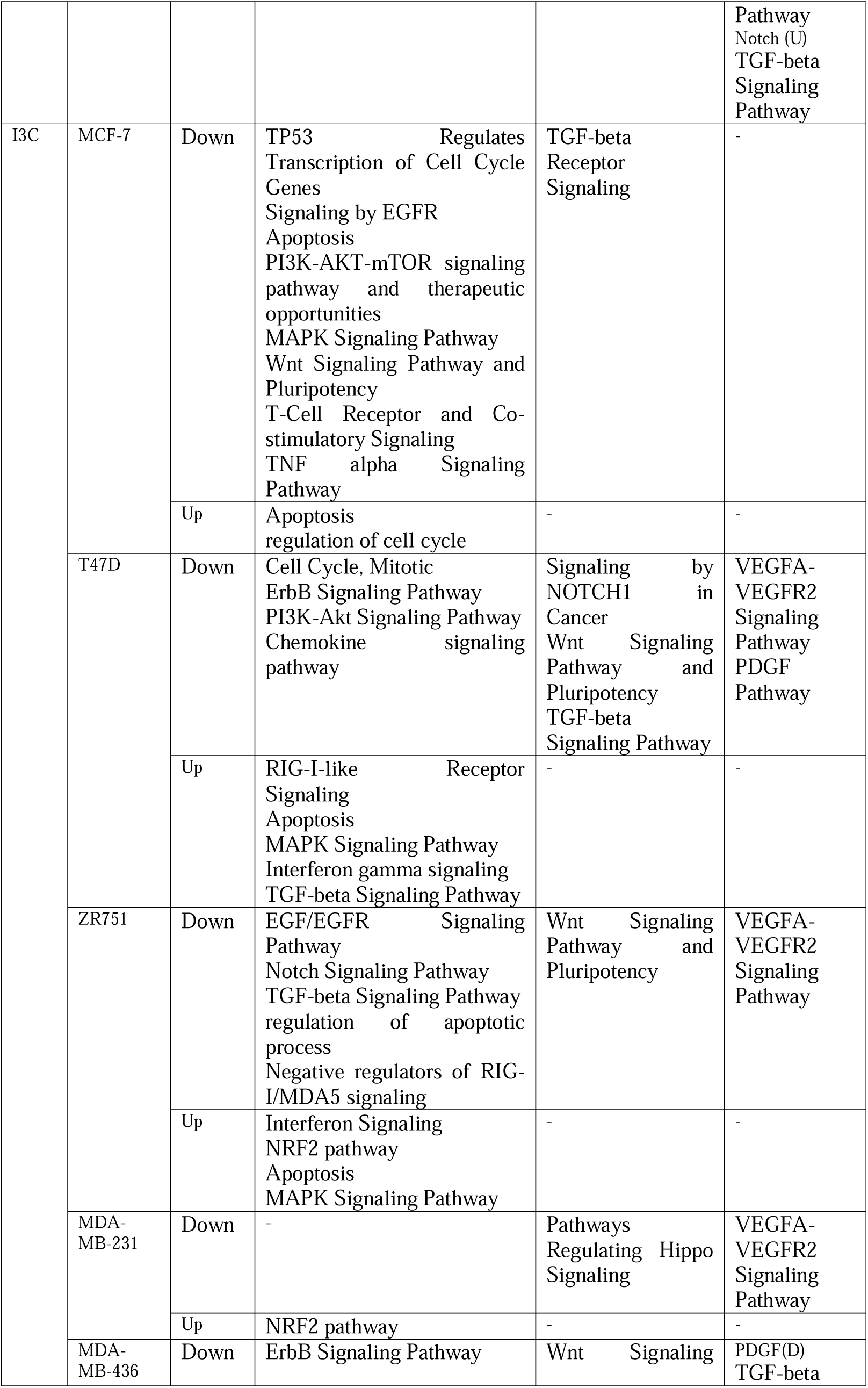
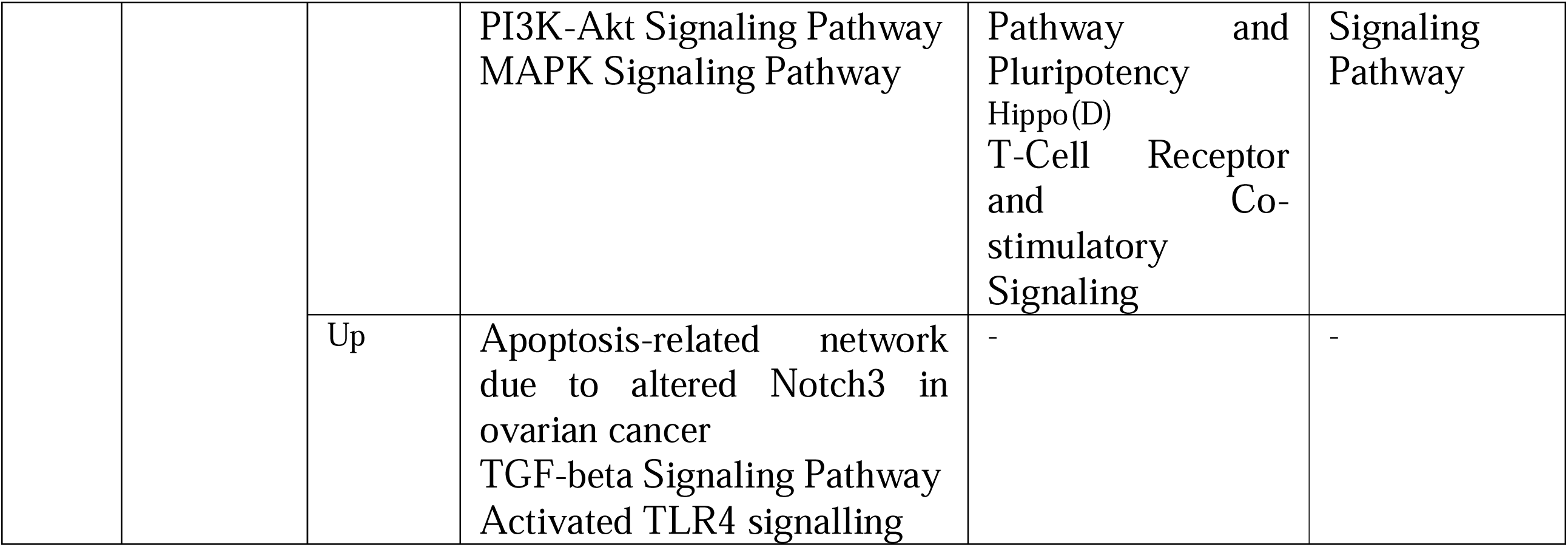
Grouping of targeted canonical oncogenic signaling pathways based on related cancer pathophysiologic processes. Three major oncological processes defining the diverse molecular processes associated with carcinogenesis were used to deduce biological roles of the various enriched oncological signaling pathways.

## Discussion

Systems pharmacology has evolved as a data-driven approach to bridge the gap between the increased amount of compound/drug perturbation data and drug discovery through systematic evaluations^36,47^. It gives new perspectives to drug/compound treated clinical and experimental publicly available omics data through well-grounded bioinformatics data analysis pipelines, speeding up the rate of understanding of the molecular mechanisms of action to identify targets of drug candidates^1,2,48^. In this study, we have mapped transcriptome data on protein interactome to construct networks targeted by plant derived compounds (Figure 1). Given the poly-pharmacologic properties of these compounds, the approach employed in this work projects the cellular behaviour in response to treatment on a physical interaction network; thereby, simplifying biological effect inference from omics data. Below, we provide correlates of our main findings with literature reports on the studied compounds.

In terms of drug specific molecular targets, actein, which is a widely studied natural triterpene glycoside that has been used by different civilizations for centuries to treat different ailments, has recently attracted attention in breast cancer due to its effects on various biological processes in cancer^18,49–52^. In this work, actein is shown to target multiple oncogenic signaling pathways in HER2+ (MDA-MB-453) subtype, of which four (Cell cycle, PI3K-Akt-mTOR and TGF-beta) are canonical oncogenic pathways^31^. Analysis of dysregulated subnetworks revealed that cell death and cell cycle arrest-related genes in TGF-beta, PI3K-Akt-mTOR and NRF2 pathways were up-regulated while cell cycle and proliferation genes in TGF-beta pathway were down-regulated. Additionally, the tumor microenvironment regulation through cytokine represented by interferon signaling pathway is down-regulated (**Table 3**). Available reports on breast and other cancers indicate that actein targets cell apoptosis^50,51^, cell adhesion^52^ and migration^50,52^. Notably, we observed actein to target oncogenic signal transduction pathways mainly regulating cell cycle/proliferation and apoptosis processes in this cell type.

CKI is an ancient formulation in the Chinese pharmacopoeia; derived from a mixture of *Radix sophorae flavescentis* and *Rhizoma smilactis glabrae* herbs. Mixed results were reported for its molecular effects on breast cancer ^53^. At the network scale, we found CKI to down-regulate P53 pathway, and up-regulate RTK-RAS-MAPK (EGFR, p38 and ErbB), PI3K-Akt-mTOR, NRF2 and TGF-beta pathways in MCF-7; which are defined canonical oncogenic signaling pathways^31^. Based on enriched genes, these pathways regulated cell proliferation and apoptosis (P53, RTK-RAS-MAPK, PI3K-Akt-mTOR and NRF2) and metastasis/invasion (TGF-beta). Moreover, CKI also targeted angiogenesis and tumor microenvironment regulating pathways through VEGFA/VEGFR2 and cytokine signaling (B cell receptor, T cell receptor and FC-epsilon signaling) respectively (**Supplementary Table 5**), which is consistent with a previous report^54^. The down-regulation of P53 pathway is in line with a previous observation of P53 independent apoptotic cell death^19^. Additional reports from other groups have shown that CKI directly regulates hepatocellular carcinoma (HCC) cell proliferation^55^, cell migration in HCC, colon and breast cancer^54,55^; and apoptosis in breast cancer^54^. The results in this work, therefore, provided further insights into the global effects of CKI on signal transduction pathways associated with Luminal A type carcinogenesis, and defined cell cycle/proliferation and apoptosis, metastasis/invasion, and angiogenesis as the targeted carcinogenesis processes (**Table 3**).

Indole 3-carbinol is a phytohormone derived from cruciferous vegetables and is a breakdown product of glucosinate 3-ylmethylglucosinate compound. It is a widely studied plant phytohormone and its therapeutic effectiveness is well defined in oestrogen receptor driven cancers^20,56–58^. We found a higher number of signal transduction terms in LA than TN subtypes. In LA cell types, these pathways were established to regulate cell proliferation and apoptosis (Wnt, cell cycle, Notch and TGF-beta) and invasion/metastasis (TGF-beta, Wnt and Notch). Taking into account the enriched genes in the different pathways, dramatic observations were made on TGF-beta, whose metastasis/invasion promoting genes were down-regulated in T47D and MCF-7 while cell death related genes were up-regulated in T47D and down-regulated in ZR751 (**Supplementary Table 5**). These pathways were spread across cell cycle/proliferation and apoptosis, metastasis/invasion, and angiogenesis as targeted carcinogenesis processes (**Table 3**).

The role of indole 3-carbinol on TN is less well elaborated and has been noted to be less effective in this cancer subtype^20^. Accordingly, in this study no oncogenic signaling pathway was enriched in the MDA-MB-157 subnetwork; illustrating an indole 3-carbinol -specific non-responsive subtype at the pathway level. This genomic tumor subtype has been previously found to be resistant to most chemotherapeutic interventions^59^, again exemplifying the unique response to treatment by cancers of the same subtype. Overall, more MDA-MB-436 signaling pathway terms were targeted by indole 3-carbinol than in MDA-MB-231 subtype (**Supplementary Table 5**). Further we found that these pathways control carcinogenesis by regulating cell cycle/proliferation and apoptosis, metastasis/invasion, and angiogenesis processes (**Table 3**).

Withaferin A is a steroidal lactone belonging to the withanolide group of compounds; a plant-derived natural compound from *Withania somnifera.* It is a vital component of the Indian Ayurvedic medicine. The characteristic anti-cancer effects of Withaferin A are well anchored in several scientific reports^21,60–70^. Multiple carcinogenesis processes have been proposed to be affected in breast cancer by WA^21,63–65,71^. Here, RTK-RAS-MAPK, TGF-beta, NRF2 and P53 oncogenic signaling pathways were found to be targeted in both TN and LA. Differential effects on Wnt, Notch, VEGFA-VEGFR2 and PI3K-Akt-mTOR in TN and cytokines in LA were observed (**Table 3**). Moreover, WA also targeted cytokine mediated signaling in both cells. The up-regulation of NRF2 pathway genes as observed is consistent with in vivo findings of induction of oxidative stress in the two cell lines^63,72^. These results illustrated multi-targeting of several carcinogenesis processes including cell proliferation and death, metastasis/invasion and angiogenesis (**Table 3**) in both TN and LA to induce the physiological phenotypes anchored in *in vitro* studies^21,63–65,71^.

Whereas this work attempts to associate the various targeted networks with cancer processes to explain the mechanism of action of poly-pharmacologic natural products, a major limitation arises on enumerating the benefits of such targets on the overall tumor prognosis. For instance, enrichment of a pathway in either up- or down-regulated subnetworks may not necessarily be directly translated as activation or inactivation of the related pathway-defined cellular process, as the same process may be indirectly co-/dys-regulated by other pathways which are either up- or down-regulated by the same drug. However, the *in vitro* reports on the activity of different drugs on cell lines^18–21^ provides an argumentative window to support current anti-cancer mechanism hypotheses. Hence, as more evidence trickles in, the results in this work will gain more proofs to support the current hypotheses. In line with common practice nowadays, we propose that integration of more omics data in drug related studies will provide a more precise picture on the exact mechanism of action of natural products^73^.

Another common challenge experienced in integrative biology approaches for targeted signal transduction pathway discovery from PPIN is its un-directionality. Thus, given the inherent directionality in such pathways, our future studies will rely on integrated directed networks from such bioresources as KEGG, REACTOME and WikiPathways, by leveraging on their comprehensiveness to define all-inclusive interaction networks for these studies.

Another interesting study that can be performed from these transcriptome datasets is simulation of the effect of drug combination to determine synergistic and antagonistic combinations especially in cases where the same cell type is treated with different natural products. As a foundation, Regan-Fendt *et al.*^74^ recently developed a computational drug combination approach using transcriptome data and disease specific root genes for malignant melanoma and successfully predicted vemurafenib and tretinoin as synergistic therapeutic combinations. Variants of this approach, for instance, modelling the active drug subnetworks using deep learning algorithms, could be applied to systematically screen and predict plant-derived compound combinations for precision medicine applications in complex diseases.

## Conclusion

This study generated two main outputs: (i) it proposed a data-driven workflow for the elucidation of the mechanism of action of pleiotropic natural product drug candidates from transcriptome data and protein interactome networks and (ii) by focussing on perturbed oncogenic signaling pathways existing in the constructed active protein interaction subnetworks, this approach demonstrated that plant-derived drugs (actein, indole-3-carbinol, withaferin A and CKI) are capable of simultaneously regulating multiple carcinogenesis processes. Thus, by sidestepping the abortive exquisite ‘target’ approach, this network-centric method can extract subtle systemic drug effects on cellular pathways that would otherwise be disregarded by the former. Aside from hypothesis generation in drug research, this approach would be applicable in precision medicine by predicting drug targeted networks and evaluating such targets against disease networks in a systems pharmacology approach. Importantly, by demonstrating the differential responses of cancers of the same/different subtype to the same compound, our work further challenges the prominent ‘one-size-fits-all’ paradigm in pharmacotherapy.

## Supplementary Figures

**Supplementary Figure 1:**
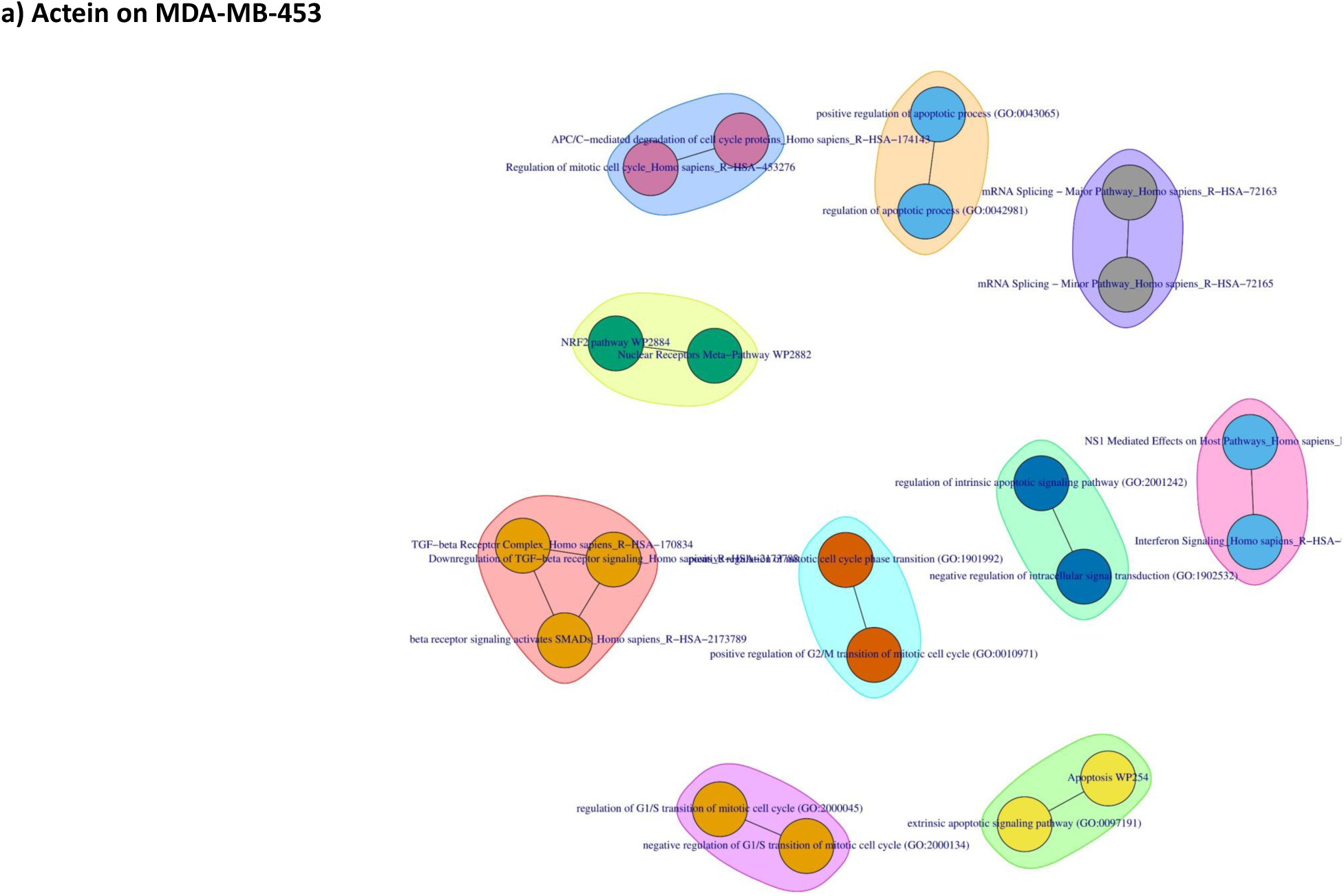
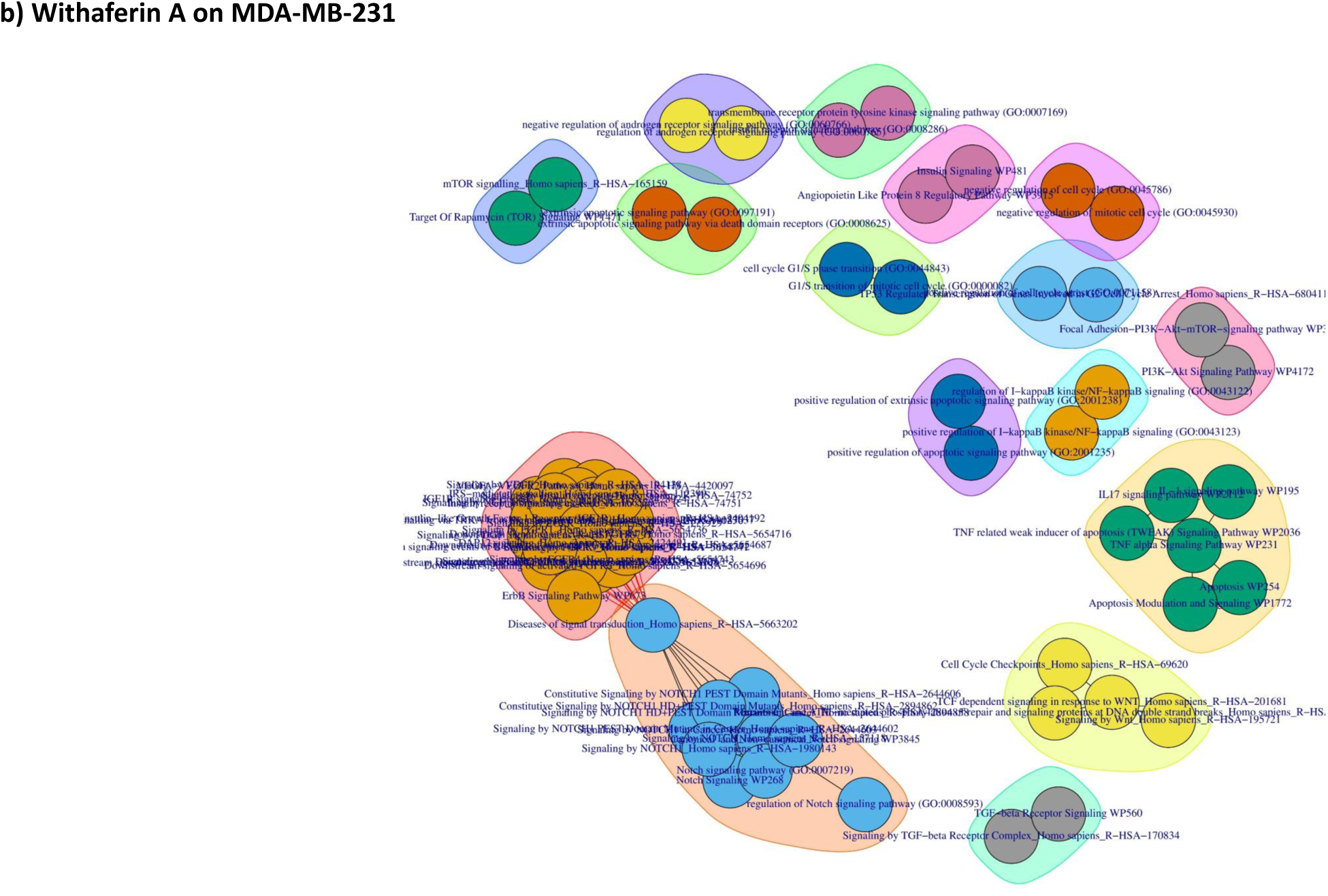
Principal component analysis (PCA) results of transcriptome samples for each dataset illustrating the distribution of variance in the first two components considered for sample separation. PC1: principal component 1, PC2: principal component 2. (a) actein on MDA-MB-453, (b) CKI on MCF-7, (c) Indole-3-Carbinol on MCF-7, (d) Indole-3-Carbinol on MDA-MB-231, (e) Indole-3-Carbinol on MDA-MB-436, (f) Indole-3-Carbinol on T47D, (g) Indole-3-Carbinol on ZR751, (h) Withaferin A on MCF-7 and (i) Withaferin A on MDA-MB-231.

**Supplementary Figure 2:**
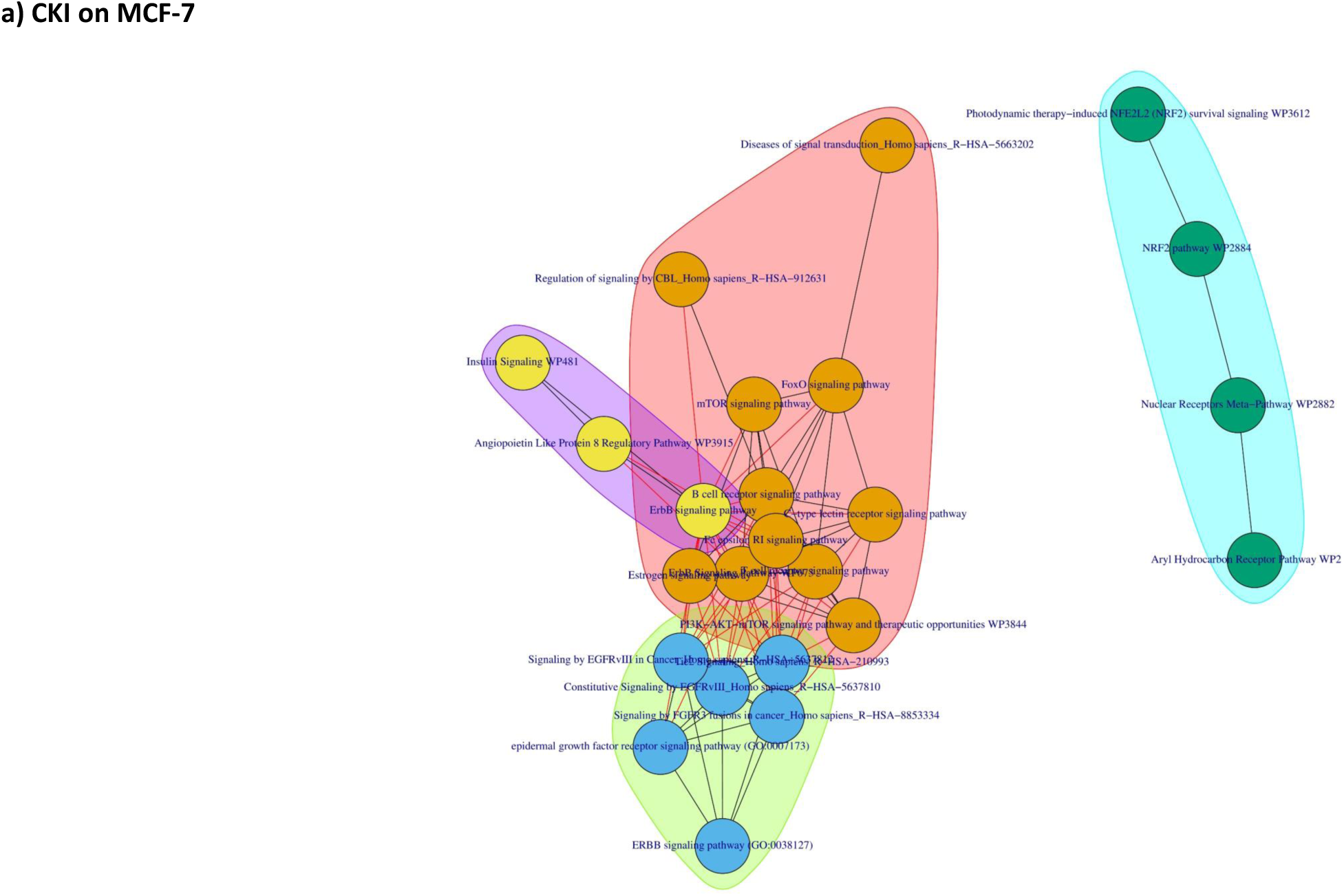
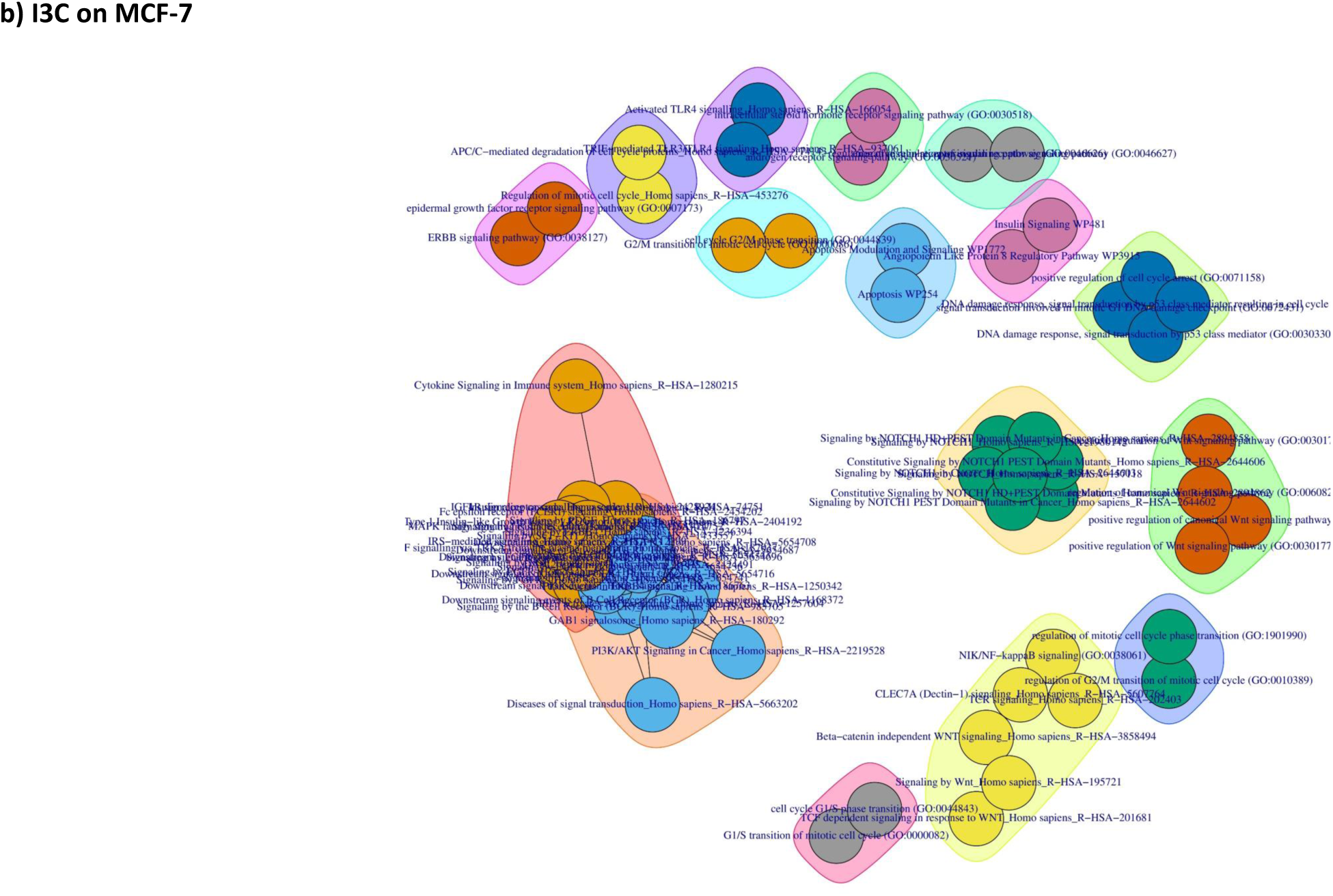
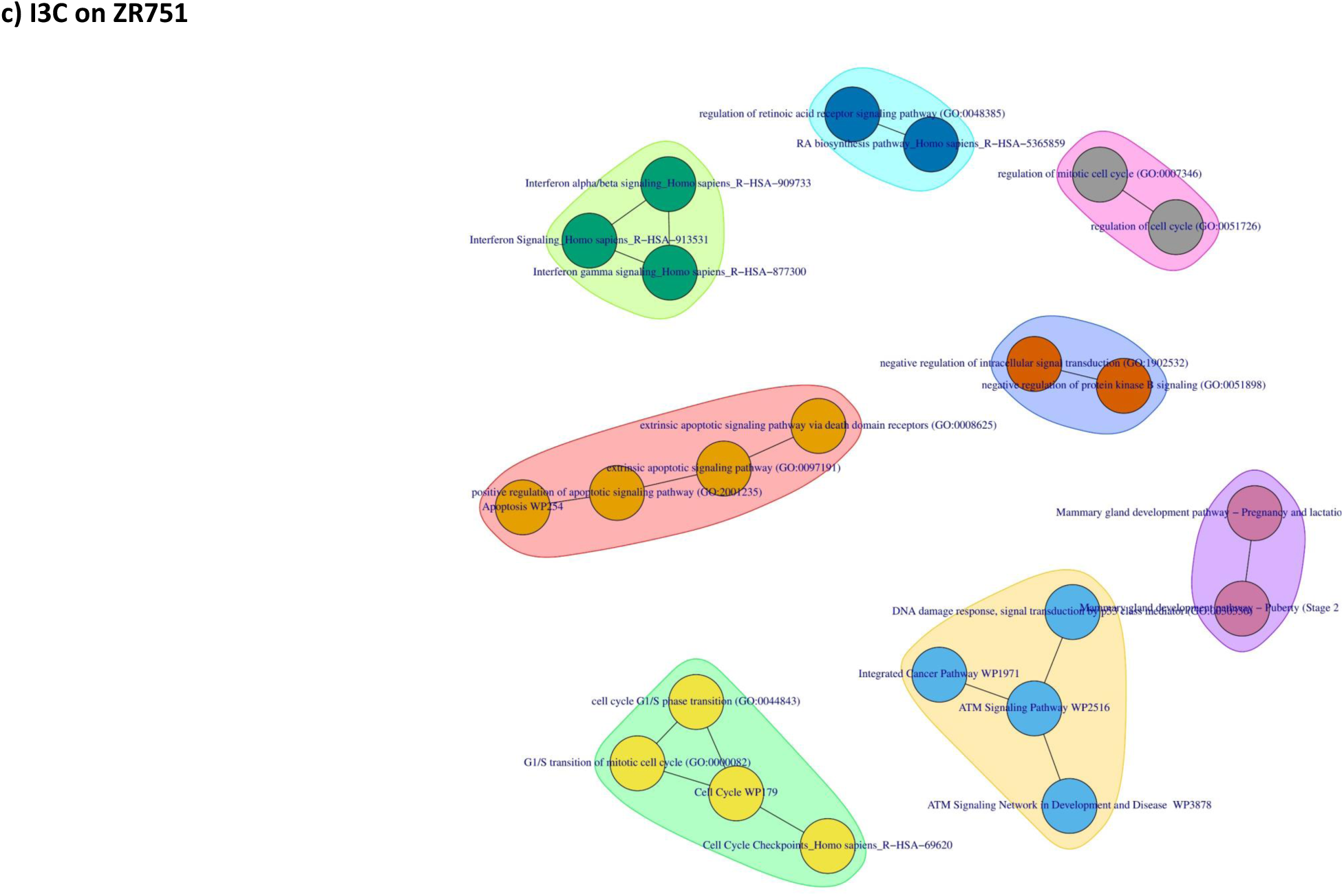
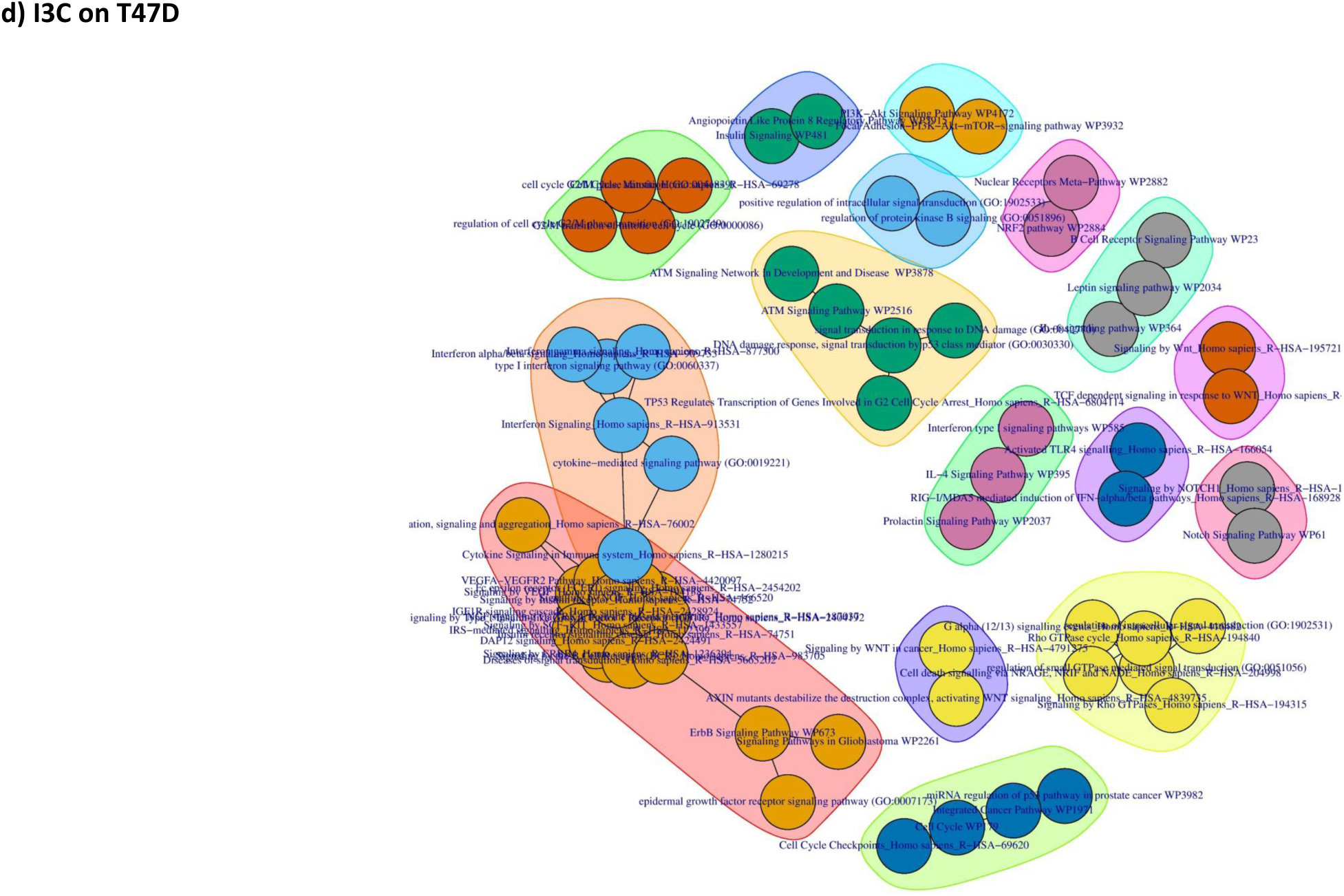
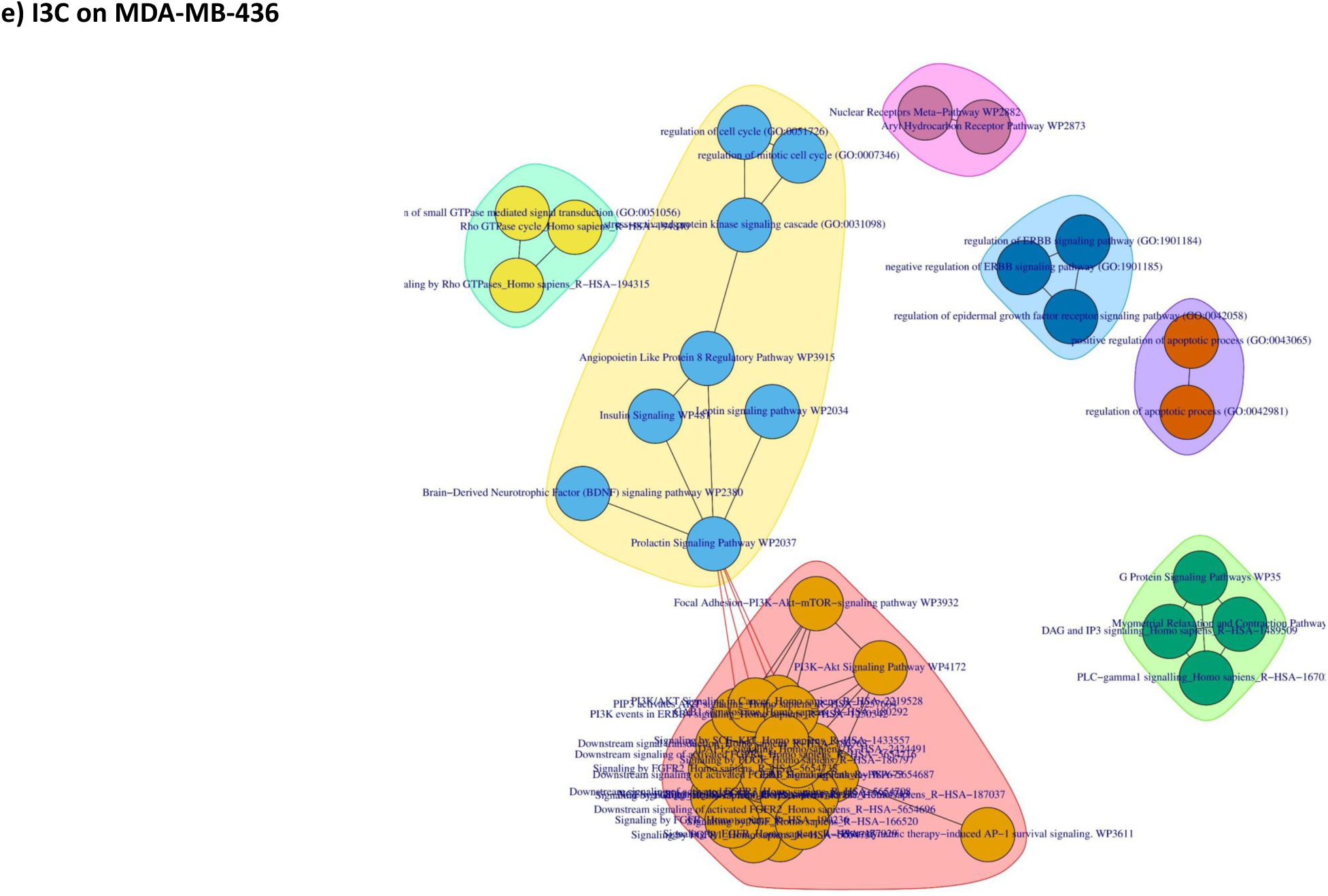
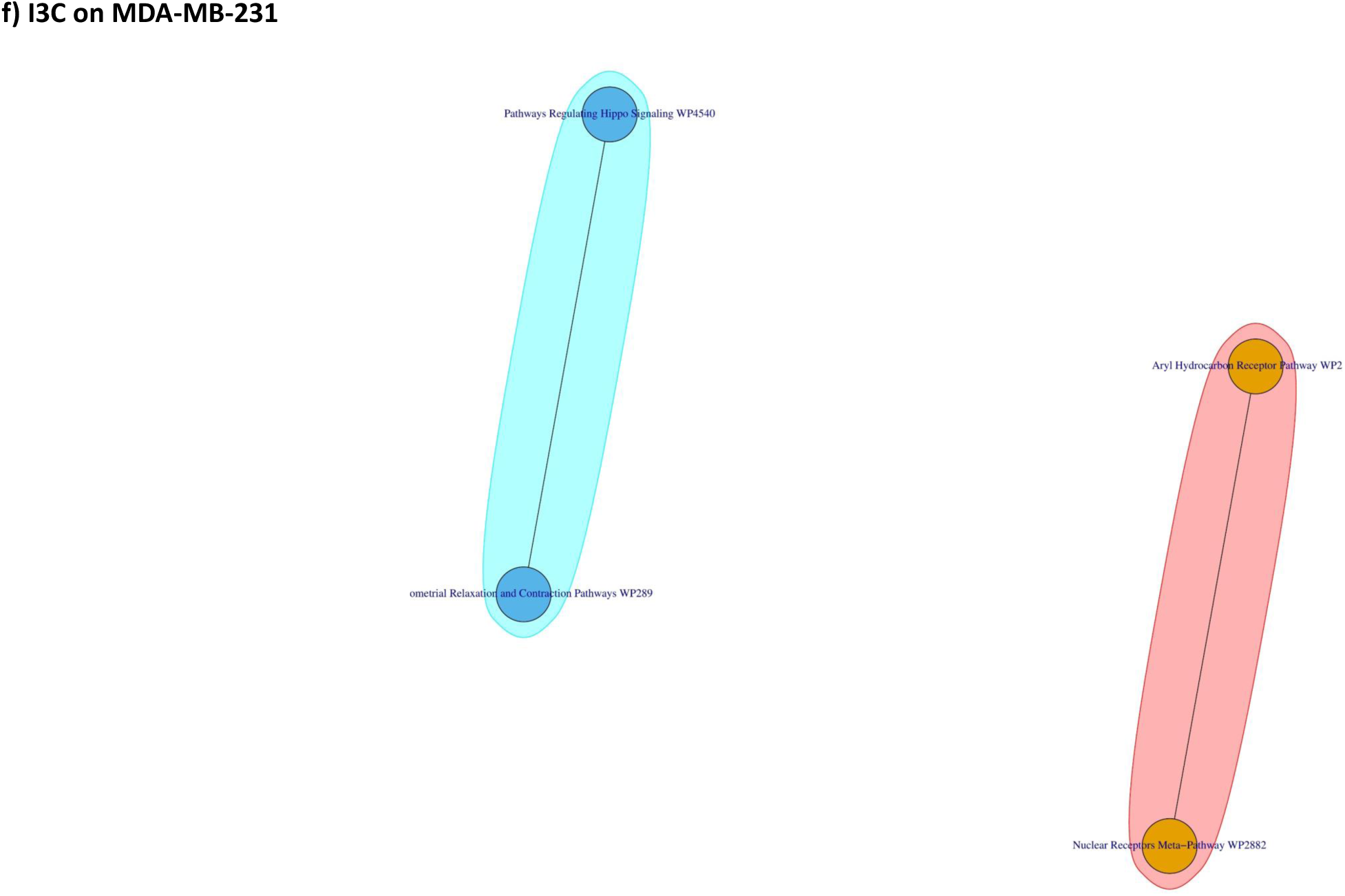
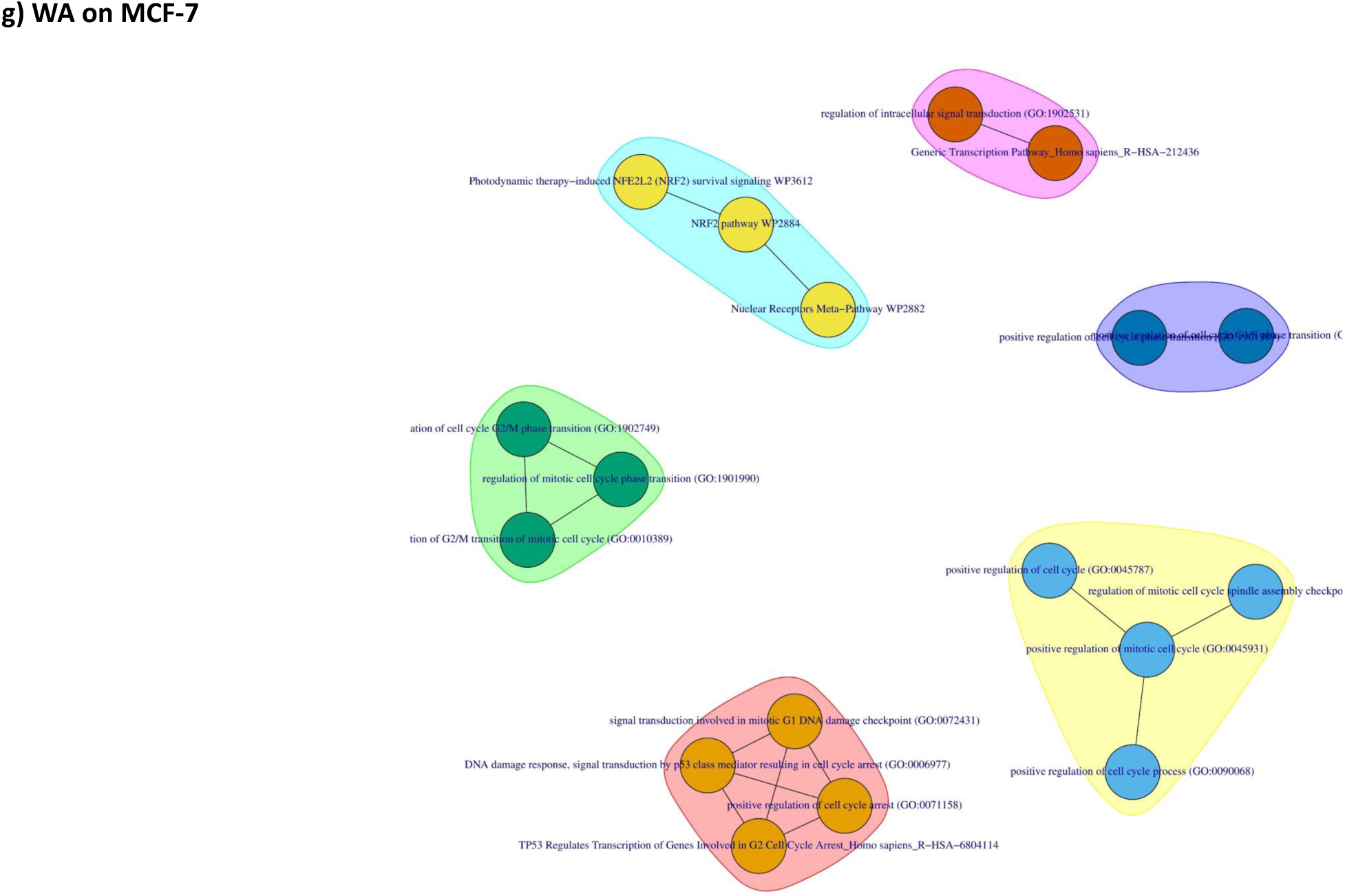
Prospective validation plots of most frequent central genes in the subnetworks. a-e) Overall survival plots showing bifurcate (APP, ELAVL1 and TRIM25), 75% vs 25% (HNRNPL) and 75% (ESR2) gene expression in relation to patient overall survival across TCGA breast cancer datasets. ‘High’ and ‘Low’ denotes patient cohorts with high median gene expression over the follow-up period. Logrank (p-value) < 0.05. f-j) Box-plots showing gene-phenotype (primary, normal and metastatic) association.

**Supplementary Figure 3:**
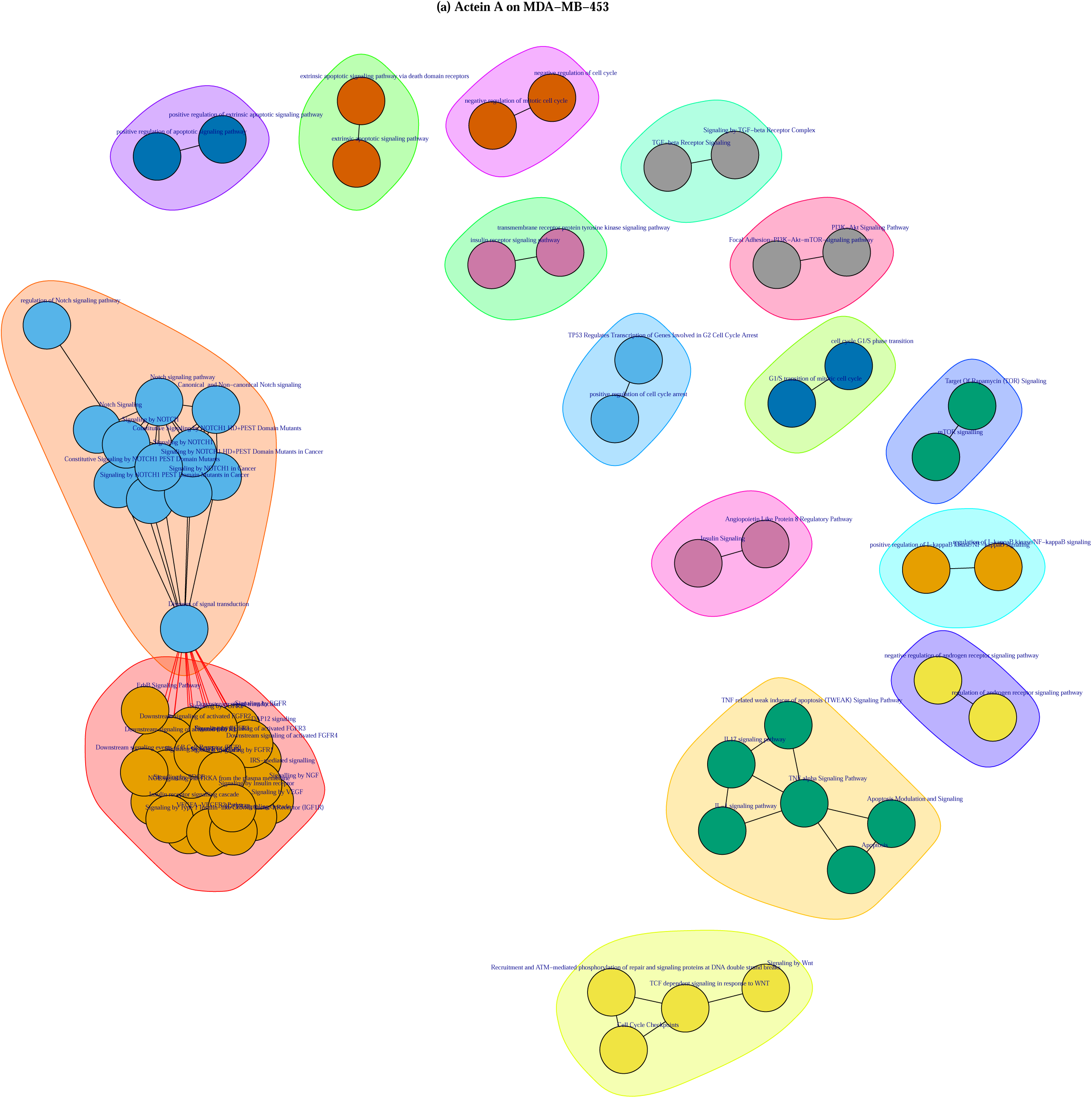
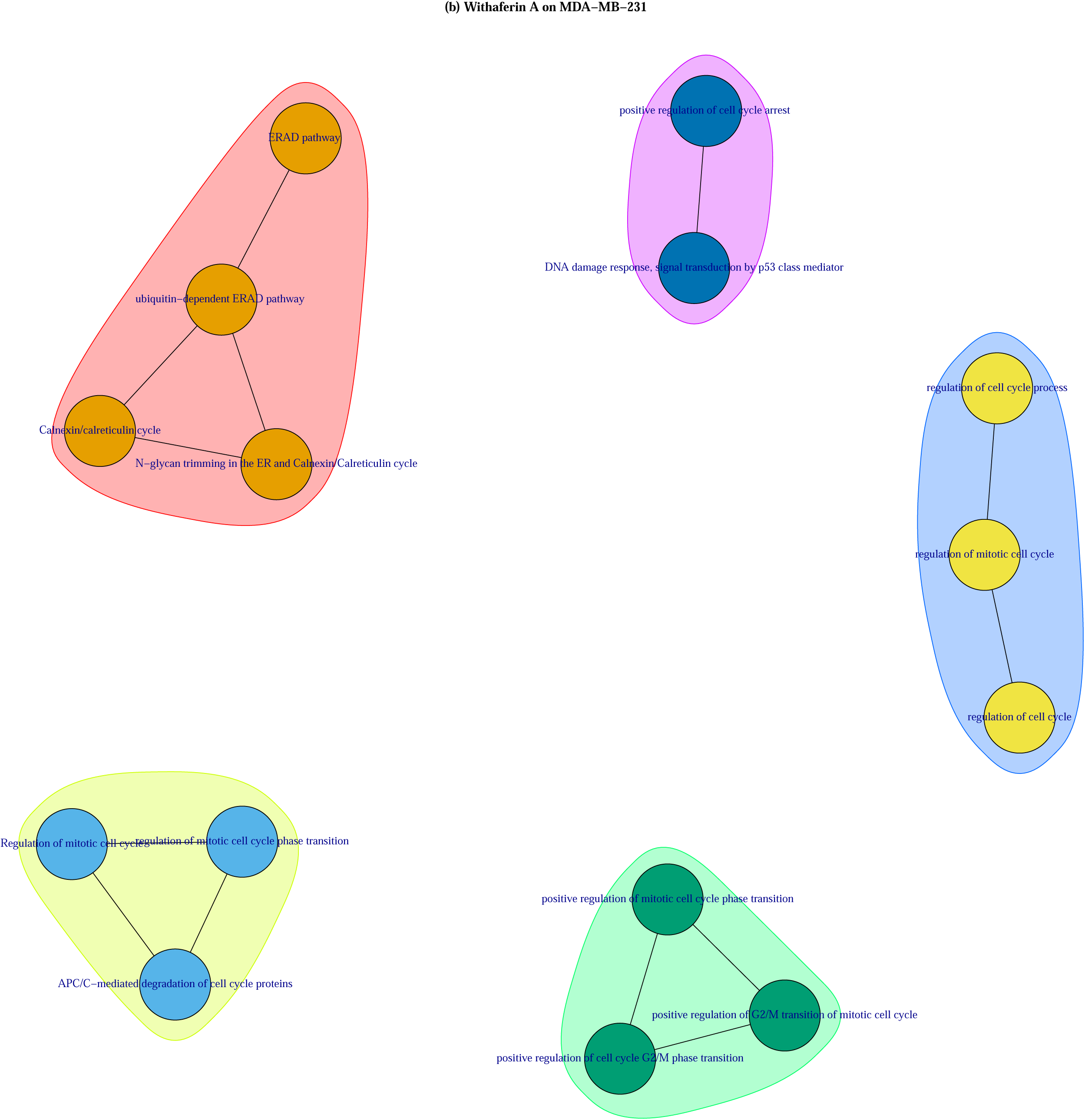
Pathway-pathway interaction networks based on shared enriched genes illustrating functional pathway cross-talk. The differently coloured clusters illustrate highly related pathways terms based on intersecting pathways. a-g: represents networks of pathways targeted by CKI on MCF-7, I3C on MCF-7, I3C on MDA-MB-436, I3C on T47D, I3C on ZR751 and WA on MCF-7.

## Supplementary Tables

**Supplementary Table 1**: Summary of the transcriptome datasets used and the molecular profiles of the cell lines. The columns Controls and Treatments list the number of samples in each case. (HER2+: human epidermal receptor 2 positive, LA: luminal A, TN: triple negative, AC: adenocarcinoma, IDC: invasive ductal carcinoma, MC: medullary carcinoma, Wt: wild type, Mut: Mutant, Del: deleted).

**Supplementary Table 2**: Summary of the differential expression analysis results. The number of differentially expressed genes under the respective plant-derived drugs/compounds are given in the table. DEG: Differentially expressed genes, FDR: False discovery rate, FC: Fold change.

**Supplementary Table 3**: Results of subnetwork betweenness- and degree centrality analysis.

**Supplementary Table 4**: Pathways enriched in whole subnetworks. FDR <0.05.

**Supplementary Table 5**: Enriched pathways in up- and down-regulated subnetworks.

**Supplementary Table 6**: An example of Actein targeted oncogenesis processes illustrating the approach used in grouping the oncogenic signaling pathways into different cancer pathophysiological processes based on the pathways’ enriched genes.

## Additional Information

### Ethics approval and consent to participate

This work did not require any ethical approval or consent as only publicly available data were used in this work.

### Consent for publication

All authors confirm the authenticity of the information provided and consent to the publication of this manuscript.

### Data availability

All relevant data are provided together with this manuscript and any additional data including the R scripts can be supplied upon request.

### Author’s contribution

R.O, A.D.Z and T.C conceived the study. R.O performed the simulations. R.O, A.D.Z and T.C contributed to the scientific discussion and data interpretation. R.O and T.C wrote the manuscript. All authors reviewed the manuscript.

## Acknowledgements

The authors would like to thank the Computational Systems Biology Laboratory and the team at the Department of Bioengineering of Gebze Technical University for offering useful insights and the computational infrastructure used in this work.

## References

1. Hopkins, A. L. Network pharmacology: the next paradigm in drug discovery. Nat. Chem. Biol. 4, 682–690 (2008).

2. Boran, A. D. W. & Iyengar, R. Systems pharmacology. Mt. Sinai J. Med. 77, 333–44 (2010).

3. Erler, J. T. & Linding, R. Network medicine strikes a blow against breast cancer. Cell 149, 731–3 (2012).

4. Barabási, A.-L., Gulbahce, N. & Loscalzo, J. Network Medicine: A Network-based Approach to Human Disease. Nat. Rev. Genet. 12, 56 (2011).

5. Rosen, L. S., Ashurst, H. L. & Chap, L. Targeting signal transduction pathways in metastatic breast cancer: a comprehensive review. Oncologist 15, 216–35 (2010).

6. Waks, A. G. & Winer, E. P. Breast Cancer Treatment. JAMA 321, 288 (2019).

7. Greenwell, M. & Rahman, P. K. S. M. Medicinal Plants: Their Use in Anticancer Treatment. Int. J. Pharm. Sci. Res. 6, 4103–4112 (2015).

8. Barabási, A.-L. & Oltvai, Z. N. Network biology: understanding the cell’s functional organization. Nat. Rev. Genet. 5, 101–113 (2004).

9. Mason, O. & Verwoerd, M. Graph Theory and Networks in Biology. (2006).

10. Mitra, K., Carvunis, A.-R., Ramesh, S. K. & Ideker, T. Integrative approaches for finding modular structure in biological networks. Nat. Rev. Genet. 14, 719–732 (2013).

11. Batra, R. et al. On the performance of de novo pathway enrichment. npj Syst. Biol. Appl. 3, 6 (2017).

12. Cursons, J. et al. Stimulus-dependent differences in signalling regulate epithelial-mesenchymal plasticity and change the effects of drugs in breast cancer cell lines. Cell Commun. Signal. 13, 26 (2015).

13. Engelmann, J. C. et al. Causal Modeling of Cancer-Stromal Communication Identifies PAPPA as a Novel Stroma-Secreted Factor Activating NFκB Signaling in Hepatocellular Carcinoma. PLOS Comput. Biol. 11, e1004293 (2015).

14. Yan, H. et al. Aberrant expression of cell cycle and material metabolism related genes contributes to hepatocellular carcinoma occurrence. Pathol. - Res. Pract. 213, 316–321 (2017).

15. AbdulHameed, M. D. M. et al. Systems Level Analysis and Identification of Pathways and Networks Associated with Liver Fibrosis. PLoS One 9, e112193 (2014).

16. Cantone, M. et al. A gene regulatory architecture that controls region□independent dynamics of oligodendrocyte differentiation. Glia 67, 825–843 (2019).

17. Frisch, T. et al. MS Atlas – A molecular map of brain lesion stages in progressive multiple sclerosis. bioRxiv 584920 (2019). doi:10.1101/584920

18. Einbond, L. S. et al. The growth inhibitory effect of actein on human breast cancer cells is associated with activation of stress response pathways. Int. J. Cancer 121, 2073–2083 (2007).

19. Qu, Z. et al. Identification of candidate anti-cancer molecular mechanisms of Compound Kushen Injection using functional genomics. Oncotarget 7, 66003–66019 (2016).

20. Caruso, J. A. et al. Indole-3-carbinol and its N-alkoxy derivatives preferentially target ER α -positive breast cancer cells. Cell Cycle 13, 2587–2599 (2014).

21. Szarc vel Szic, K. et al. Pharmacological Levels of Withaferin A (Withania somnifera) Trigger Clinically Relevant Anticancer Effects Specific to Triple Negative Breast Cancer Cells. PLoS One 9, e87850 (2014).

22. Ritchie, M. E. et al. limma powers differential expression analyses for RNA-sequencing and microarray studies. Nucleic Acids Res. 43, e47–e47 (2015).

23. Alcaraz, N. et al. Efficient key pathway mining: combining networks and OMICS data. Integr. Biol. 4, 756–764 (2012).

24. Oughtred, R. et al. The BioGRID interaction database: 2019 update. Nucleic Acids Res. 47, D529–D541

25. Tang, Y., Li, M., Wang, J., Pan, Y. & Wu, F.-X. CytoNCA: A cytoscape plugin for centrality analysis and evaluation of protein interaction networks. Biosystems 127, 67–72 (2015).

26. Chen, X., Miao, Z., Divate, M., Zhao, Z. & Cheung, E. KM-express: an integrated online patient survival and gene expression analysis tool for the identification and functional characterization of prognostic markers in breast and prostate cancers. Database (Oxford). 2018, (2018).

27. Kuleshov, M. V et al. Enrichr: a comprehensive gene set enrichment analysis web server 2016 update. Nucleic Acids Res. 44, W90–7 (2016).

28. Liu, K.-Q., Liu, Z.-P., Hao, J.-K., Chen, L. & Zhao, X.-M. Identifying dysregulated pathways in cancers from pathway interaction networks. BMC Bioinformatics 13, 126 (2012).

29. Wang, Q., Shi, C.-J. & Lv, S.-H. Benchmarking pathway interaction network for colorectal cancer to identify dysregulated pathways. Brazilian J. Med. Biol. Res. = Rev. Bras. Pesqui. medicas e Biol. 50, e5981 (2017).

30. Ju, W., Li, J., Yu, W. & Zhang, R. iGraph: an incremental data processing system for dynamic graph. Front. Comput. Sci. 10, 462–476 (2016).

31. Sanchez-Vega, F. et al. Oncogenic Signaling Pathways in The Cancer Genome Atlas. Cell 173, 321-337.e10 (2018).

32. Desmedt, C. et al. Biological Processes Associated with Breast Cancer Clinical Outcome Depend on the Molecular Subtypes. Clin. Cancer Res. 14, 5158–5165 (2008).

33. Dai, X., Cheng, H., Bai, Z. & Li, J. Breast Cancer Cell Line Classification and Its Relevance with Breast Tumor Subtyping. J. Cancer 8, 3131–3141 (2017).

34. Alcaraz, N., Kücük, H., Weile, J., Wipat, A. & Baumbach, J. KeyPathwayMiner: Detecting Case-Specific Biological Pathways Using Expression Data. Internet Math. 7, 299–313 (2011).

35. Beisser, D., Klau, G. W., Dandekar, T., Muller, T. & Dittrich, M. T. BioNet: an R-Package for the functional analysis of biological networks. Bioinformatics 26, 1129–1130 (2010).

36. van der Greef, J. & McBurney, R. N. Rescuing drug discovery: in vivo systems pathology and systems pharmacology. Nat. Rev. Drug Discov. 4, 961–967 (2005).

37. Wozniak, J. & Ludwig, A. Novel role of APP cleavage by ADAM10 for breast cancer metastasis. EBioMedicine 38, 5–6 (2018).

38. Walsh, L. A. et al. An Integrated Systems Biology Approach Identifies TRIM25 as a Key Determinant of Breast Cancer Metastasis. Cell Rep. 20, 1623–1640 (2017).

39. Wang, J. et al. Multiple functions of the RNA-binding protein HuR in cancer progression, treatment responses and prognosis. Int. J. Mol. Sci. 14, 10015–41 (2013).

40. Xiu, B., Chi, Y., Ji, W., Zhang, Q. & Wu, J. Abstract P6-05-08: LINC02273 interacts with hnRNPL and promotes metastasis through directly activating AGR2 in breast cancer. in P6-05-08-P6-05–08 (American Association for Cancer Research (AACR), 2019). doi:10.1158/1538-7445.sabcs18-p6-05-08

41. Maguire, P. et al. Estrogen receptor beta (ESR2) polymorphisms in familial and sporadic breast cancer. Breast Cancer Res. Treat. 94, 145–152 (2005).

42. Costa, J. Systems medicine in oncology. Nat. Clin. Pract. Oncol. 5, 117–117 (2008).

43. Levitsky, D. O. & Dembitsky, V. M. Anti-breast Cancer Agents Derived from Plants. Nat. Products Bioprospect. 5, 1 (2014).

44. Chen, Y.-A. et al. Integrated Pathway Clusters with Coherent Biological Themes for Target Prioritisation. PLoS One 9, e99030 (2014).

45. Wajant, H. The Role of TNF in Cancer. in Results and problems in cell differentiation 49, 1–15 (2009).

46. Boutet, E. et al. UniProtKB/Swiss-Prot, the Manually Annotated Section of the UniProt KnowledgeBase: How to Use the Entry View. in 23–54 (Humana Press, New York, NY, 2016). doi:10.1007/978-1-4939-3167-5_2

47. 49 van Hasselt, J. G. C. & Iyengar, R. Systems Pharmacology: Defining the Interactions of Drug Combinations. Annu. Rev. Pharmacol. Toxicol. 59, 21–40 (2019).

48. Zhang, G.-B., Li, Q.-Y., Chen, Q.-L. & Su, S.-B. Network pharmacology: a new approach for chinese herbal medicine research. Evid. Based. Complement. Alternat. Med. 2013, 621423 (2013).

49. Yue, G. G.-L. et al. New potential beneficial effects of actein, a triterpene glycoside isolated from Cimicifuga species, in breast cancer treatment. Sci. Rep. 6, 35263 (2016).

50. Zhang, Y., Lian, J. & Wang, X. Actein inhibits cell proliferation and migration and promotes cell apoptosis in human non-small cell lung cancer cells. Oncol. Lett. 15, 3155–3160 (2018).

51. Ji, L. et al. Actein induces autophagy and apoptosis in human bladder cancer by potentiating ROS/JNK and inhibiting AKT pathways. Oncotarget 8, 112498–112515 (2017).

52. Wu, X.-X. et al. Actein Inhibits the Proliferation and Adhesion of Human Breast Cancer Cells and Suppresses Migration in vivo. Front. Pharmacol. 9, 1466 (2018).

53. Yu, L., Zhou, Y., Yang, Y., Lu, F. & Fan, Y. Efficacy and Safety of Compound Kushen Injection on Patients with Advanced Colon Cancer: A Meta-Analysis of Randomized Controlled Trials. Evid. Based. Complement. Alternat. Med. 2017, 7102514 (2017).

54. Nourmohammadi, S. et al. Effect of Compound Kushen Injection, a Natural Compound Mixture, and Its Identified Chemical Components on Migration and Invasion of Colon, Brain, and Breast Cancer Cell Lines. Front. Oncol. 9, 314 (2019).

55. Gao, L. et al. Uncovering the anticancer mechanism of Compound Kushen Injection against HCC by integrating quantitative analysis, network analysis and experimental validation. Sci. Rep. 8, 624 (2018).

56. Weng, J.-R., Tsai, C.-H., Kulp, S. K. & Chen, C.-S. Indole-3-carbinol as a chemopreventive and anti-cancer agent. Cancer Lett. 262, 153–63 (2008).

57. Hyo, A. Indole-3-Carbinol Mediated Anti-Proliferative Regulation of Breast Cancer Stem Cells and Malignant Melanoma-Initiating Cells. (2016).

58. Katz, E., Nisani, S. & Chamovitz, D. A. Indole-3-carbinol: a plant hormone combatting cancer. F1000Research 7, (2018).

59. Chavez, K. J., Garimella, S. V & Lipkowitz, S. Triple negative breast cancer cell lines: one tool in the search for better treatment of triple negative breast cancer. Breast Dis. 32, 35–48 (2010).

60. Choi, M. J., Park, E. J., Min, K. J., Park, J.-W. & Kwon, T. K. Endoplasmic reticulum stress mediates withaferin A-induced apoptosis in human renal carcinoma cells. Toxicol. Vitr. 25, 692–698 (2011).

61. vel Szic, K. S. et al. Epigenetic silencing of triple negative breast cancer hallmarks by Withaferin A. Oncotarget 8, 40434–40453 (2017).

62. Sehrawat, A. et al. Withaferin A-mediated apoptosis in breast cancer cells is associated with alterations in mitochondrial dynamics. Mitochondrion (2019). doi:10.1016/J.MITO.2019.01.003

63. Ghosh, K., De, S., Das, S., Mukherjee, S. & Sengupta Bandyopadhyay, S. Withaferin A Induces ROS-Mediated Paraptosis in Human Breast Cancer Cell-Lines MCF-7 and MDA-MB-231. PLoS One 11, e0168488 (2016).

64. Hahm, E.-R. & Singh, S. V. Withaferin A-induced apoptosis in human breast cancer cells is associated with suppression of inhibitor of apoptosis family protein expression. Cancer Lett. 334, 101–8 (2013).

65. Ghosh, K. et al. Withaferin A induced impaired autophagy and unfolded protein response in human breast cancer cell-lines MCF-7 and MDA-MB-231. Toxicol. Vitr. 44, 330–338 (2017).

66. Hassannia, B. et al. Nano-targeted induction of dual ferroptotic mechanisms eradicates high-risk neuroblastoma. J. Clin. Invest. 128, 3341–3355 (2018).

67. Yu, Y. et al. Withaferin-A kills cancer cells with and without telomerase: chemical, computational and experimental evidences. Cell Death Dis. 8, e2755–e2755 (2017).

68. Bargagna-Mohan, P. et al. The Tumor Inhibitor and Antiangiogenic Agent Withaferin A Targets the Intermediate Filament Protein Vimentin. Chem. Biol. 14, 623–634 (2007).

69. Stan, S. D., Hahm, E.-R., Warin, R. & Singh, S. V. Withaferin A Causes FOXO3a- and Bim-Dependent Apoptosis and Inhibits Growth of Human Breast Cancer Cells In vivo. Cancer Res. 68, 7661–7669 (2008).

70. Manoharan, S., Panjamurthy, K., Balakrishnan, S., Vasudevan, K. & Vellaichamy, L. Circadian time-dependent chemopreventive potential of withaferin-A in 7,12-dimethyl-benz[a]anthracene-induced oral carcinogenesis. (2009).

71. Wang, H.-C. et al. Different effects of 4β-hydroxywithanolide E and withaferin A, two withanolides from Solanaceae plants, on the Akt signaling pathway in human breast cancer cells. Phytomedicine 53, 213–222 (2019).

72. Hahm, E.-R. et al. Withaferin A-Induced Apoptosis in Human Breast Cancer Cells Is Mediated by Reactive Oxygen Species. PLoS One 6, e23354 (2011).

73. Chaudhary, K., Poirion, O. B., Lu, L. & Garmire, L. X. Deep learning–based multiomics integration robustly predicts survival in liver cancer. Clin. Cancer Res. 24, (2018).

74. Regan-Fendt, K. E. et al. Synergy from gene expression and network mining (SynGeNet) method predicts synergistic drug combinations for diverse melanoma genomic subtypes. npj Syst. Biol. Appl. 5, 6 (2019).

