## Supplementary Table 2 for "A systems pharmacology approach to determine the mechanisms of action of pleiotropic natural products in breast cancer from transcriptome data"

**Supplementary Table 2: Summary of the differential expression analysis results.** The number of differentially expressed genes under the respective plant-derived drugs/compounds are given in the table. DEG: Differentially expressed genes, FDR: False discovery rate, FC: Fold change

| **Drug/Compound** | **Dosage** | **Cell Line** | **DEGs** | **FDR cut-off** | **FC cut-off** |
| --- | --- | --- | --- | --- | --- |
| Actein | 40µg/ml | MDA-MB-453 | 520 | 0.01 | 2 |
| CKI | 2mg | MCF-7 | 1661 | 0.01 | 2 |
| I3C | 200µM | MCF-7 | 3115 | 0.005 | 2 |
|  |  | T47D | 2462 | 0.005 | 2 |
|  |  | ZR751 | 2125 | 0.005 | 2 |
|  |  | MDA-MB-231 | 202 | 0.005 | 2 |
|  |  | MDA-MB-157 | 430 | 0.005 | 2 |
|  |  | MDA-MB-436 | 869 | 0.005 | 2 |
| WA | 700nM | MDA-MB-231 | 1614 | 0.005 | 2 |
|  |  | MCF-7 | 482 | 0.005 | 2 |
