## Supplementary Table 6 for "A systems pharmacology approach to determine the mechanisms of action of pleiotropic natural products in breast cancer from transcriptome data"

**Supplementary Table 6: An example of the approach used in grouping the oncogenic signalling pathways into different cancer pathophysiological processes based on each pathway's enriched genes for MDA-MB-453 cell line under Actein treatment.** The pathways highlighted here were derived from the enrichment analysis results of the whole subnetworks (**Supp Table 4**). Gene functions were derived from UniProt database. The reasons for considering each gene in arriving at a conclusion are given.

| **Oncogenic Signalling Pathway** | **Known biological role(s)** | **Gene** | **Gene Functions (uniprot)** | **Status of gene in decision** | **Targeted Oncogenesis Process** |
| --- | --- | --- | --- | --- | --- |
| Intrinsic Pathway for Apoptosis_Homo sapiens_R-HSA-109606 | (i) Cell death | TFDP1 | (i) Binds DNA cooperatively with E2F family members whose products are involved in cell cycle regulation or in DNA replication (pubmed:8405995, pubmed:7739537). (ii) The E2F1:DP complex appears to mediate both cell proliferation and apoptosis. | **Considered because of relation to cell apoptosis** | **Cell cycle/Proliferation and Apoptosis** |
|  |  | E2F1 | (i) Transcription activator that binds DNA cooperatively with DP proteins whose products are involved in cell cycle regulation or in DNA replication. (ii) The DRTF1/E2F complex functions in the control of cell-cycle progression from G1 to S phase. (iii) It can mediate both cell proliferation and TP53/p53-dependent apoptosis. | **Considered because of relation to cell apoptosis** |  |
|  |  | BID | (i) The major proteolytic product p15 BID allows the release of cytochrome c. Isoform 1, isoform 2 and isoform 4 induce ICE-like proteases and apoptosis. Isoform 3 does not induce apoptosis. Counters the protective effect of Bcl-2 | **Considered because of relation to cell apoptotic genes** |  |
|  |  | BCL2L1 | (i) Potent inhibitor of cell death. (ii) Acts as a regulator of G2 checkpoint and progression to cytokinesis during mitosis. (iii) May attenuate inflammation impairing NLRP1-inflammasome activation, hence CASP1 activation and IL1B release (pubmed:17418785). (iv) Isoform Bcl-X(S) promotes apoptosis. | **Considered because of relation to cell apoptosis** |  |
| Interferon Signaling_Homo sapiens_R-HSA-913531 | (i) immune response to viral infection (ii) antitumor immunity (iii) cell growth inhibition | NUP93 | (i) Regulates podocyte migration and proliferation through SMAD4 signalling (pubmed:26878725). | **Considered because of relation to cell proliferation** | **Cell cycle/Proliferation and Apoptosis** |
|  |  | NDC1 | (i) Component of the nuclear pore complex (NPC), which plays a key role in de novo assembly and insertion of NPC in the nuclear envelope. | **Considered because of relation to cell proliferation** |  |
|  |  | NUP205 | (i) Plays a role in the nuclear pore complex (NPC) assembly and/or maintenance (pubmed:9348540). | **Not considered** |  |
|  |  | IFITM2 | (i) IFN-induced antiviral protein which inhibits the entry of viruses to the host cell cytoplasm. (ii) Induces cell cycle arrest and mediates apoptosis by caspase activation and in p53-independent manner. | **Considered because of relation to immune response** |  |
|  |  | NUP107 | (i) Plays a role in the nuclear pore complex (NPC) assembly and/or maintenance (pubmed:12552102). | **Considered** |  |
|  |  | NUP188 | (i) May function as a component of the nuclear pore complex (NPC). | **Not considered** |  |
|  |  | NUP155 | (i) Essential for embryogenesis. (ii) Involved both in binding and translocating proteins during nucleocytoplasmic transport. | **Not considered** |  |
|  |  | NUP85 | (i) Essential component of the nuclear pore complex (NPC) that seems to be required for NPC assembly and maintenance (pubmed:12718872). (ii) Involved in CCR2-mediated chemotaxis of monocytes and may link activated CCR2 to the phosphatidyl-inositol 3-kinase-Rac-lammellipodium protrusion cascade (pubmed:15995708). | **Considered because of relation to immune response** |  |
|  |  | OAS3 | (i) Plays a critical role in cellular innate antiviral response (ii) May also play a role in other cellular processes such as apoptosis, cell growth, differentiation and gene regulation. | **Considered because of relation to apoptosis** |  |
|  |  | TRIM14 | (i) Plays a role in the innate immune defense against viruses (ii) Facilitates the type I IFN response (pubmed:24379373) (iii) Positively regulates the CGAS-induced type I interferon signalling pathway (pubmed:27666593) | **Considered because of relation to immune response** |  |
|  |  | NUP160 | (i) Involved in poly(A)+ RNA transport. | **Not considered** |  |
|  |  | PIAS1 | (i) Functions as an E3-type small ubiquitin-like modifier (SUMO) ligase, stabilizing the interaction between UBE2I and the substrate, and as a SUMO-tethering factor (ii) Plays a crucial role as a transcriptional coregulation in the STAT pathway, the p53 pathway and the steroid hormone signalling pathway. | **Considered because of relation to cell proliferation and death pathways** |  |
| PI3K-AKT-mTOR signaling pathway and therapeutic opportunities WP3844 | (i) apoptosis (ii) autophagy (iii) metastasis (iv) cell growth | RB1CC1 | (i) Involved in autophagy (pubmed:21775823). (ii) Involved in repair of DNA damage caused by ionizing radiation, which subsequently improves cell survival by decreasing apoptosis (By similarity). (iii) Plays a role as a modulator of TGF-beta-signalling. | **Considered because of relation to autophagic cell death** | **Cell cycle/Proliferation and Apoptosis** |
|  |  | EIF4EBP1 | (i) Mediates the regulation of protein translation by hormones, growth factors and other stimuli that signal through the MAP kinase and mtorc1 pathways | **Considered because of relation to cell growth and proliferation** |  |
|  |  | GRB10 | (i) Binds to, and suppress signals from, activated receptors tyrosine kinases, including the insulin (INSR) and insulin-like growth factor (IGF1R) receptors. (ii) Negatively regulates Wnt signalling. (iii) Positive regulator of the KDR/VEGFR-2 signalling pathway | **Considered because of relation to cell growth and proliferation** |  |
|  |  | PTEN | (i) Tumor suppressor. (ii) Antagonizes the PI3K-AKT/PKB signalling pathway thereby modulating cell cycle progression and cell survival. (iii) The nuclear monoubiquitinated form possesses greater apoptotic potential, whereas the cytoplasmic nonubiquitinated form induces less tumor suppressive ability. | **Considered because of relation to cell proliferation and apoptosis** |  |
|  |  | ULK1 | (i) [Involved in autophagy in response to starvation. (pubmed:25040165).](https://www.uniprot.org/citations/25040165) | **Considered because of relation to autophagic cell death** |  |
| NRF2 pathway WP2884 | (i) detoxification and metabolism of xenobiotics (ii) cell differentiation and apoptosis (iii) cell proliferation | ABCC3 | (i) Act as an inducible transporter in the biliary and intestinal excretion of organic anions. | **Considered because of relation to cell proliferation** | **Cell cycle/Proliferation and Apoptosis** |
|  |  | GCLC | (i) This protein is involved in step **1** of the subpathway that synthesizes glutathione from L-cysteine and L-glutamate | **Considered because of relation to cell proliferation** |  |
|  |  | SLC2A10 | (i) Facilitative glucose transporter required for the development of the cardiovascular system | **Not considered** |  |
|  |  | TXNRD1 | (i) Isoform 5 also mediates cell death induced by a combination of interferon-beta and retinoic acid | **Considered because of relation to immune mediated cell death** |  |
|  |  | MAFG | (i) Serve as transcriptional activators by dimerizing with other (usually larger) basic-zipper proteins, such as NFE2, NFE2L1 and NFE2L2, and recruiting them to specific DNA-binding sites (pubmed:8932385, pubmed:9421508, pubmed:11154691). (ii) Small Maf proteins heterodimerize with Fos and may act as competitive repressors of the NFE2L2 transcription factor (pubmed:11154691). (iii) May be involved in signal transduction of extracellular H+ (By similarity) | **Not considered** |  |
|  |  | FTH1 | (i) Stores iron in a soluble, non-toxic, readily available form. Important for iron homeostasis. (ii) Also plays a role in delivery of iron to cells. | **Not considered** |  |
|  |  | HMOX1 | (i) Exhibits cytoprotective effects since excess of free heme sensitizes cells to undergo apoptosis. | **Considered because of relation to apoptosis** |  |
|  |  | GCLM | (i) This protein is involved in step **1** of the subpathway that synthesizes glutathione from L-cysteine and L-glutamate. | **Not considered** |  |
|  |  | FGF13 | (i) Is involved in both polymerization and stabilization of microtubules. | **Not considered** |  |
|  |  | SQSTM1 | (i) Autophagy receptor required for selective macroautophagy (aggrephagy) (pubmed:16286508, pubmed:20168092, pubmed:24128730, pubmed:28404643, pubmed:22622177). (ii) May be involved in cell differentiation, apoptosis, immune response and regulation of K+ channels. (iii) Acts as an activator of the NFE2L2/NRF2 pathway (pubmed:20452972, pubmed:28380357). | **Considered because of relation to apoptosis, immune response and autophagy** |  |
|  |  | SLC39A14 | (i) [Broad-scope metal ion transporter with a preference for zinc uptake (pubmed:29621230).](https://www.uniprot.org/citations/29621230) | **Not considered** |  |
|  |  | FTL | (i) Stores iron in a soluble, non-toxic, readily available form. (ii) Plays a role in delivery of iron to cells. | **Not considered** |  |
| TGF-beta Signaling Pathway WP366 | (i) cancer progression (ii) invassion (iii) metastasis (iv) cell growth and survival (v) cell cycle arrest (vi) apoptosis (vii) angiogenesis (viii) inflammation | TGIF1 | (i) Active transcriptional corepressor of SMAD2. Links the nodal signalling pathway to the bifurcation of the forebrain and the establishment of ventral midline structures. | **Considered because of relation to cell proliferation** | **Cell cycle/Proliferation and Apoptosis** |
|  |  | APP | (i) Functions on the surface of neurons relevant to neurite growth, neuronal adhesion and axonogenesis. (ii) Couples to apoptosis-inducing pathways such as those mediated by g(o) and jip. | **Considered because of relation to apoptosis** |  |
|  |  | CDKN1A | (i) Involved in p53/TP53 mediated inhibition of cellular proliferation in response to DNA damage. (ii) Binds to and inhibits cyclin-dependent kinase activity and blocking cell cycle progression. (pubmed:11595739) | **Considered because of relation to cell proliferation** |  |
|  |  | ITCH | (i) Acts as an E3 ubiquitin-protein ligase (pubmed:14602072,). (ii) Mediates the antiapoptotic activity of epidermal growth factor through the ubiquitination and proteasomal degradation of p15 BID (pubmed:20392206). | **Considered because of relation to apoptosis** |  |
|  |  | KLF6 | (i) Could play a role in B-cell growth and development | **Considered because of relation to immune response** |  |
|  |  | NEDD4L | (i) E3 ubiquitin-protein ligase (ii) Inhibits TGF-beta signalling by triggering SMAD2 and TGFBR1 ubiquitination and proteasome-dependent degradation. (iii) Involved in the regulation of TOR signalling (pubmed:27694961). | **Considered because of relation to cell proliferation** |  |
|  |  | ATF3 | (i) This protein binds the camp response element (CRE) (consensus: 5'-GTGACGT[AC][AG]-3'), a sequence present in many viral and cellular promoters. Represses transcription from promoters with ATF sites. | **Considered because of relation to apoptosis** |  |
