## Supplementary Table 1 for "A systems pharmacology approach to determine the mechanisms of action of pleiotropic natural products in breast cancer from transcriptome data"

**Supplementary Table 1:** **Summary of the transcriptome datasets used and the molecular profiles of the cell lines.** The columns Controls and Treatments list the number of samples in each case. (HER2+: human epidermal receptor 2 positive, LA: luminal A, TN: triple negative, AC: adenocarcinoma, IDC: invasive ductal carcinoma, MC: medullary carcinoma, Wt: wild type, Mut: Mutant, Del: deleted).

| Drug | Platform | Cell line | Subtype | Pathology | BRCA1 | P53 | Controls | Treatments |
| --- | --- | --- | --- | --- | --- | --- | --- | --- |
| Actein | Affymetrix Human Array | MDA-MB-453 | HER2+ | AC | Wt | Del | 4 | 3 |
| CKI | Illumina HiSeq 2500 | MCF-7 | LA | IDC | Wt | Wt | 3 | 3 |
| I3C | Illumina beadchip | MCF-7 | LA | IDC | Wt | Wt | 3 | 3 |
|  |  | T47D | LA | IDC | Wt | Mut | 3 | 3 |
|  |  | ZR751 | LA | IDC | Wt | Wt | 3 | 3 |
|  |  | MDA-MB-231 | TN | MC | Wt | Mut | 3 | 3 |
|  |  | MDA-MB-157 | TN | AC | Wt | Mut | 3 | 3 |
|  |  | MDA-MB-436 | TN | AC | Mut | Mut | 3 | 3 |
| WA | Illumina beadchip | MDA-MB-231 | TN | MC | Wt | Mut | 3 | 3 |
|  |  | MCF-7 | LA | IDC | Wt | Wt | 3 | 3 |
