## Supplementary Figure 2 for "A systems pharmacology approach to determine the mechanisms of action of pleiotropic natural products in breast cancer from transcriptome data"

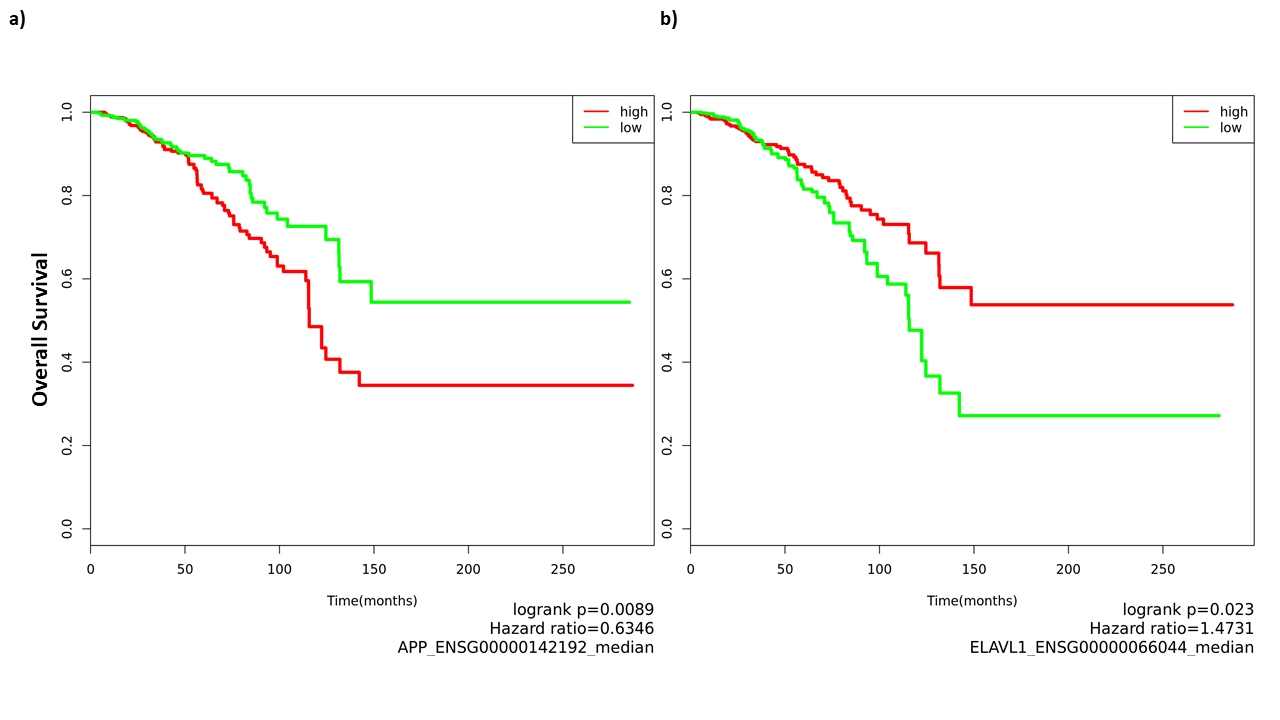

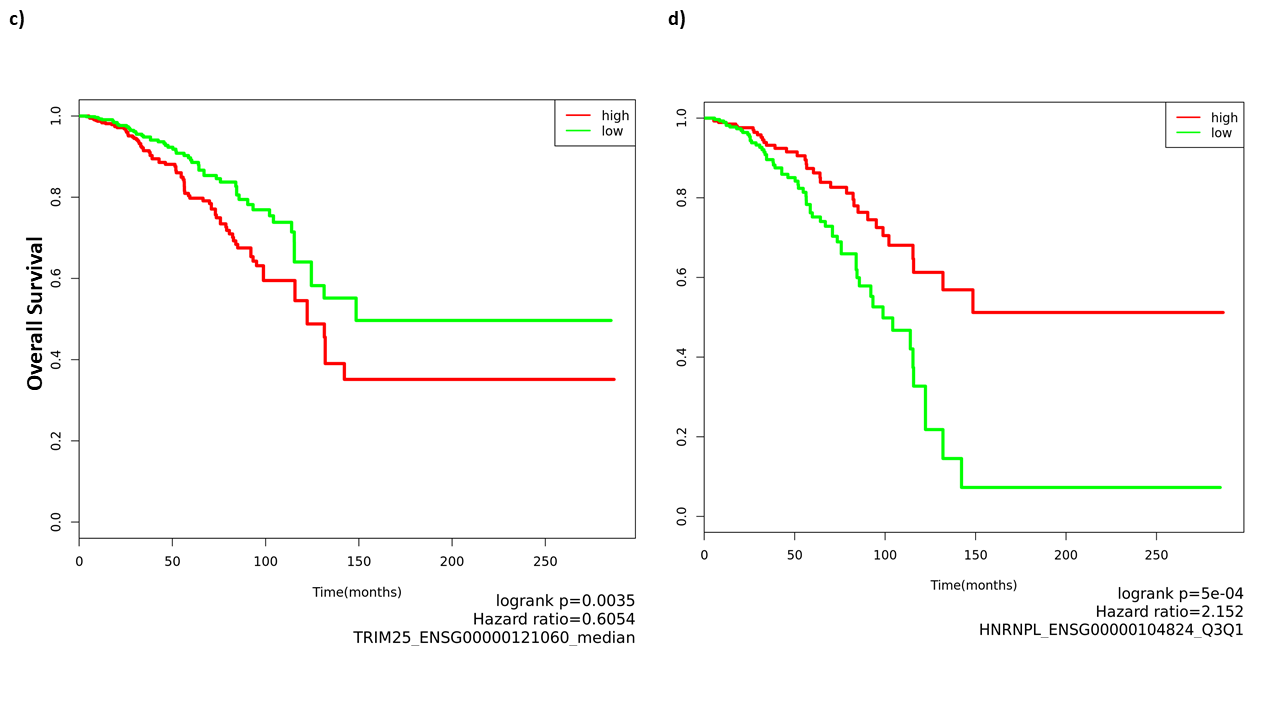

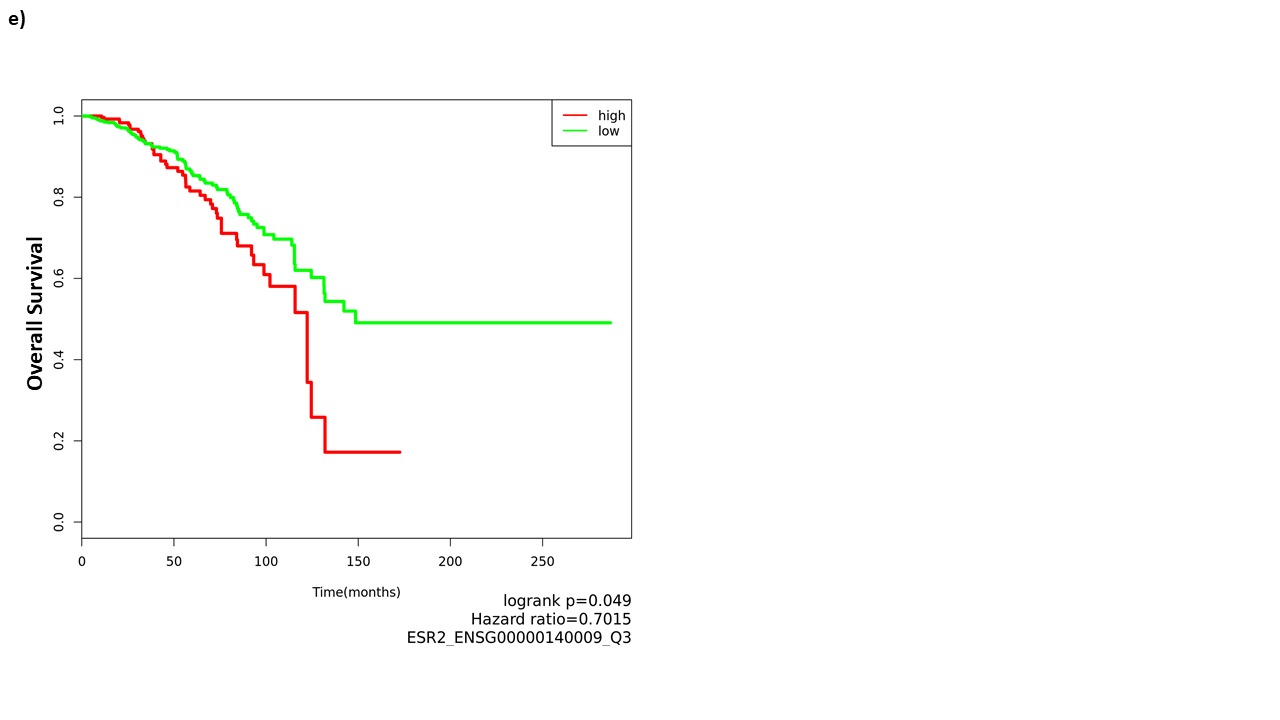

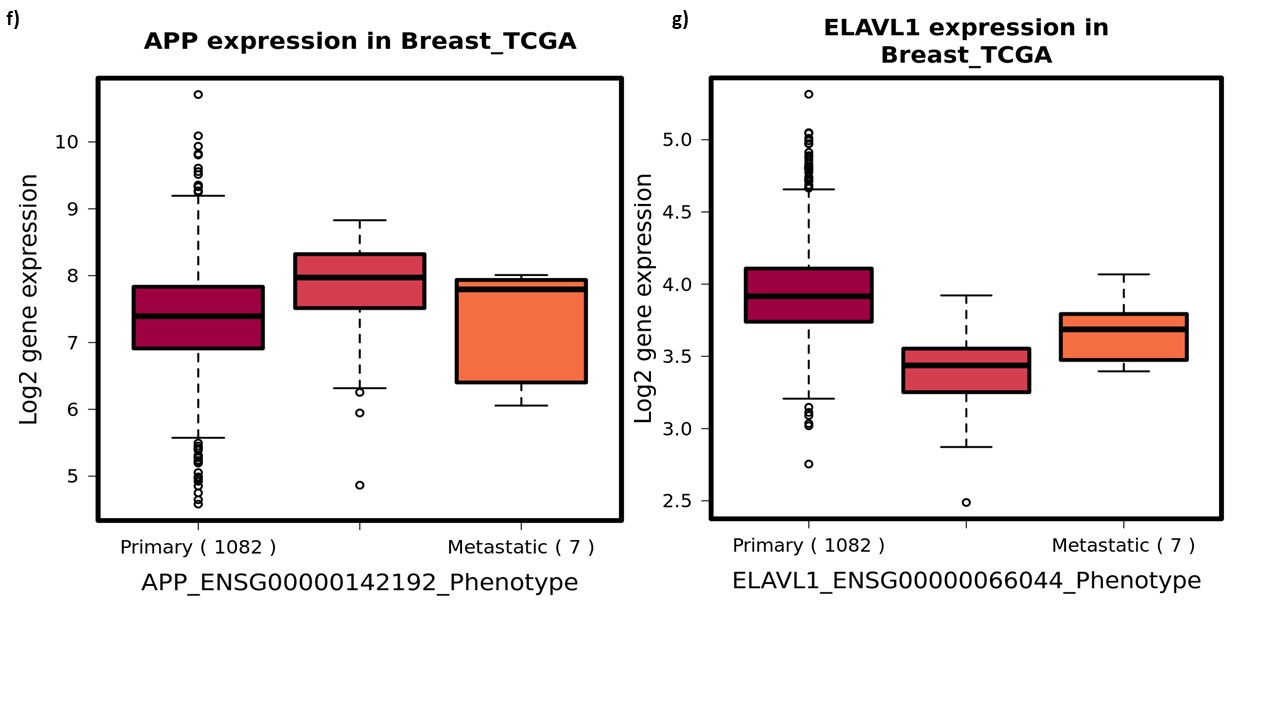

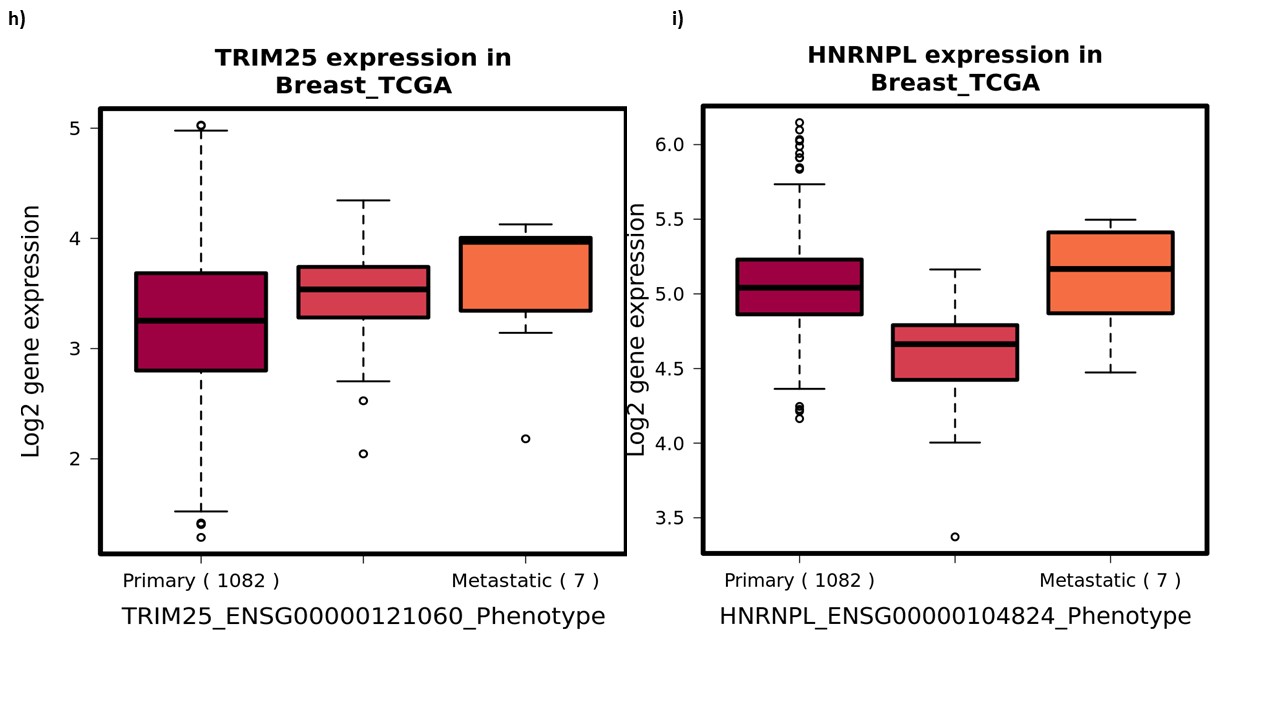

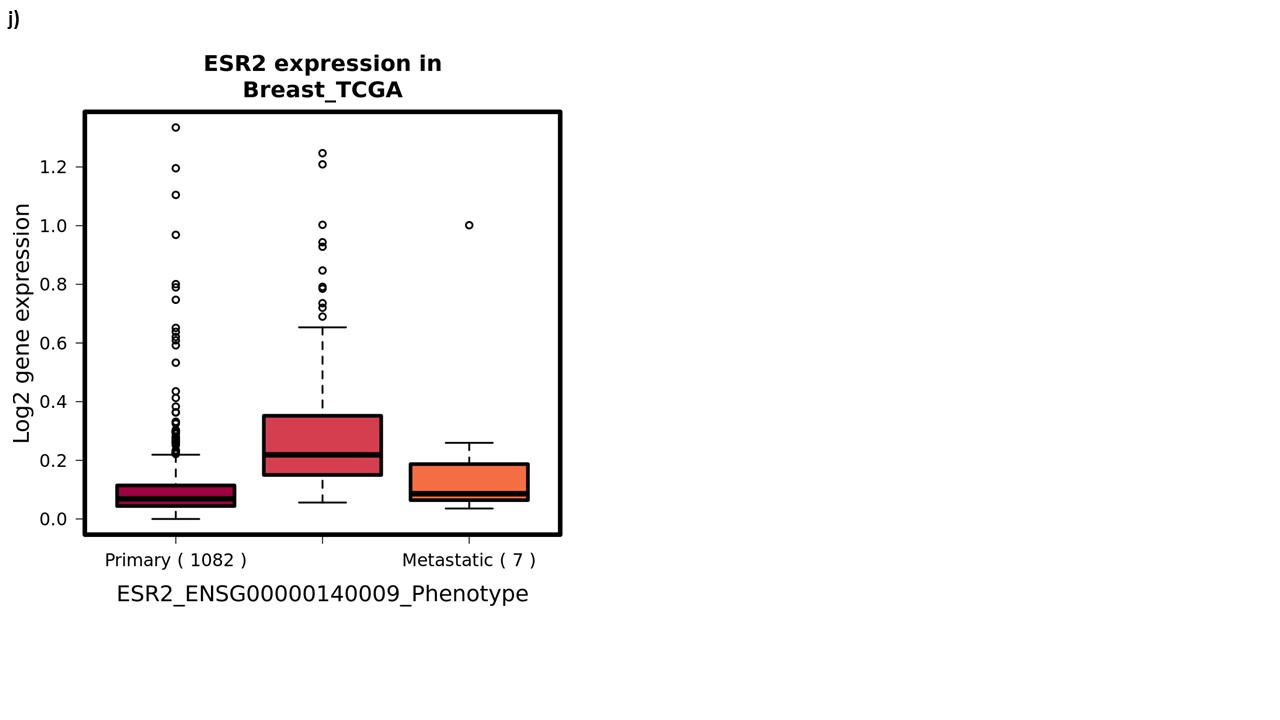

**Supplementary Figure 2: Prospective validation plots of most frequent central genes in the subnetworks**. a-e) Overall survival plots showing bifurcate (APP, ELAVL1 and TRIM25), 75% vs 25% (HNRNPL) and 75% (ESR2) gene expression in relation to patient overall survival across TCGA breast cancer datasets. ‘High’ and ‘Low’ denotes patient cohorts with high median gene expression over the follow-up period. Logrank (p-value) $<0.05$. f-j) Box-plots showing gene-phenotype (primary, normal and metastatic) association.
