## Supplementary Figure 3 for "A systems pharmacology approach to determine the mechanisms of action of pleiotropic natural products in breast cancer from transcriptome data"

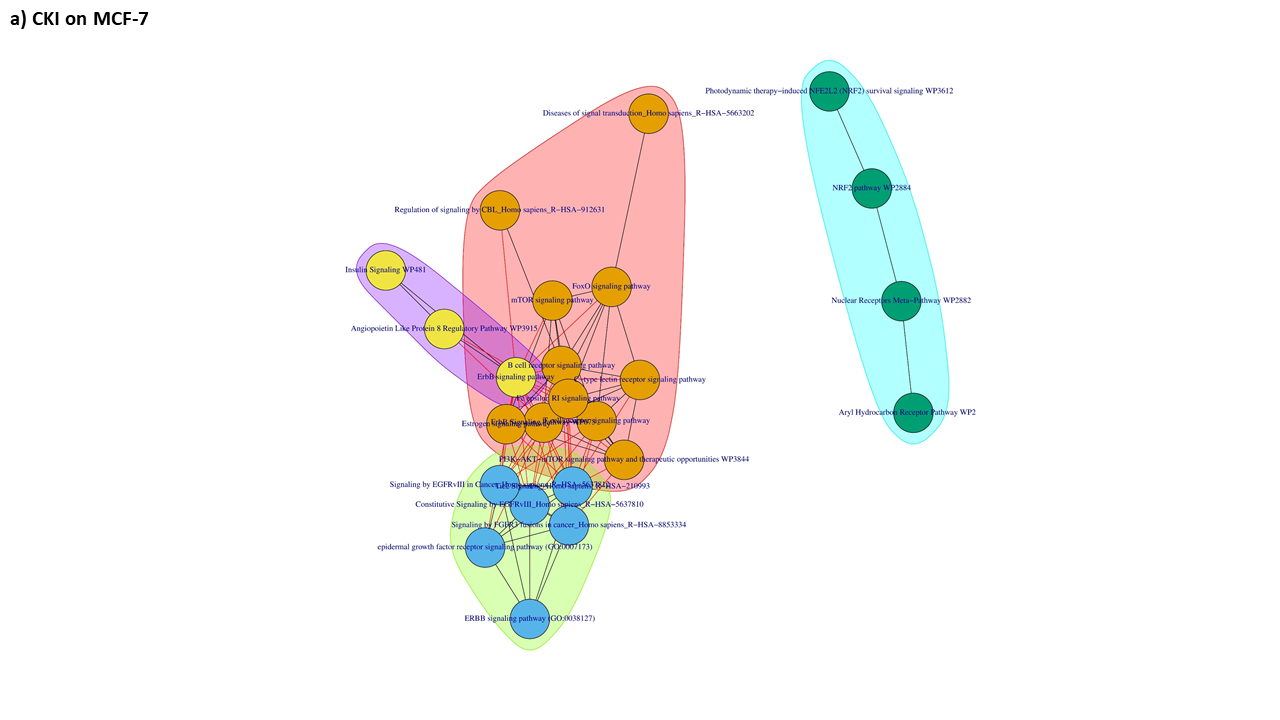

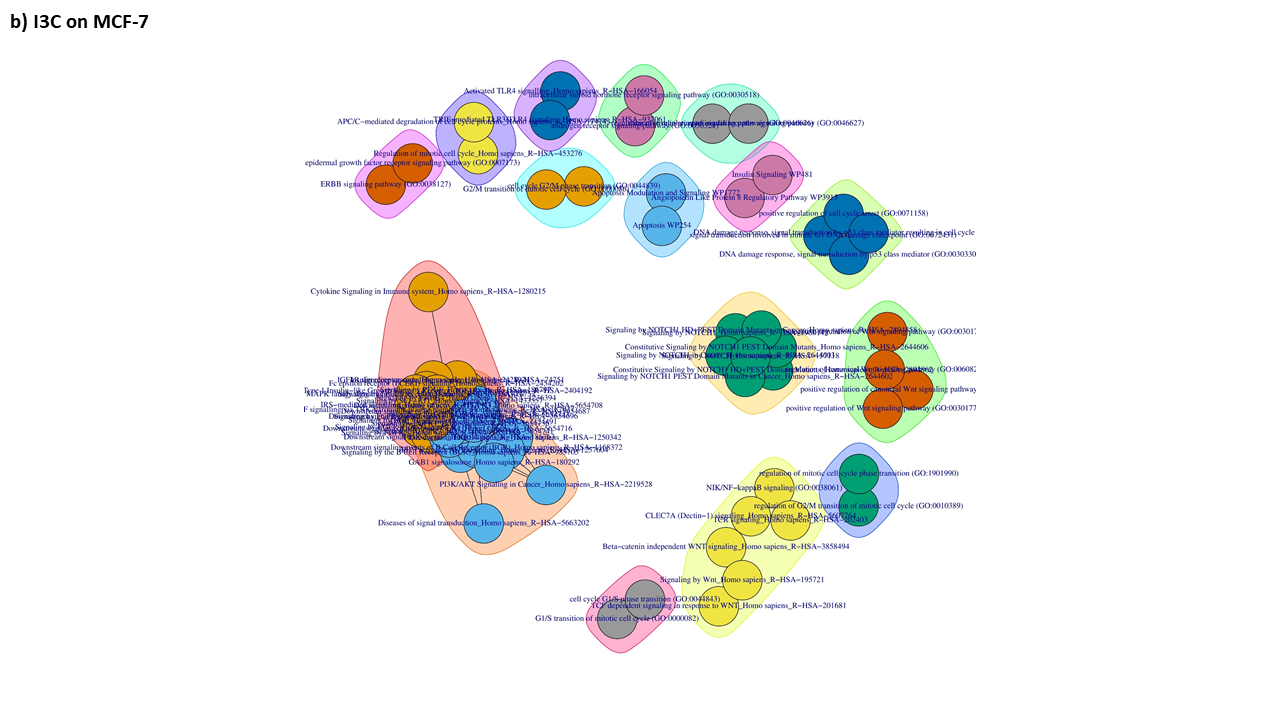

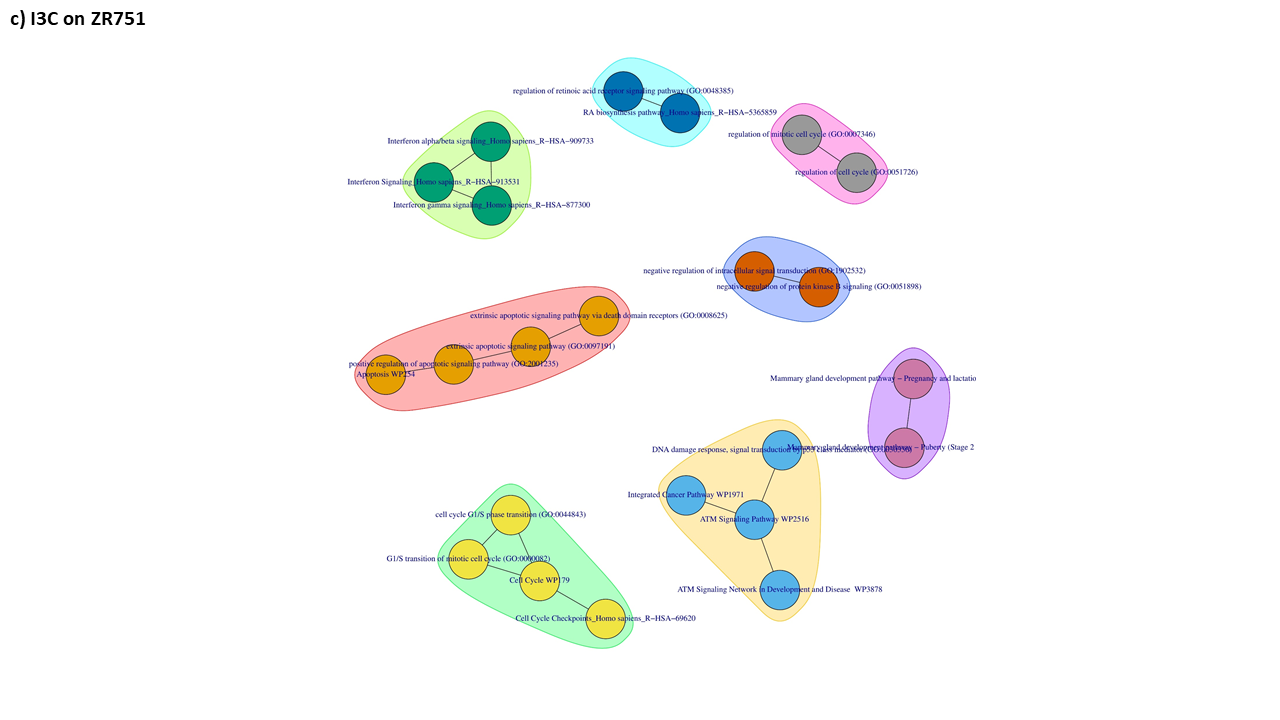

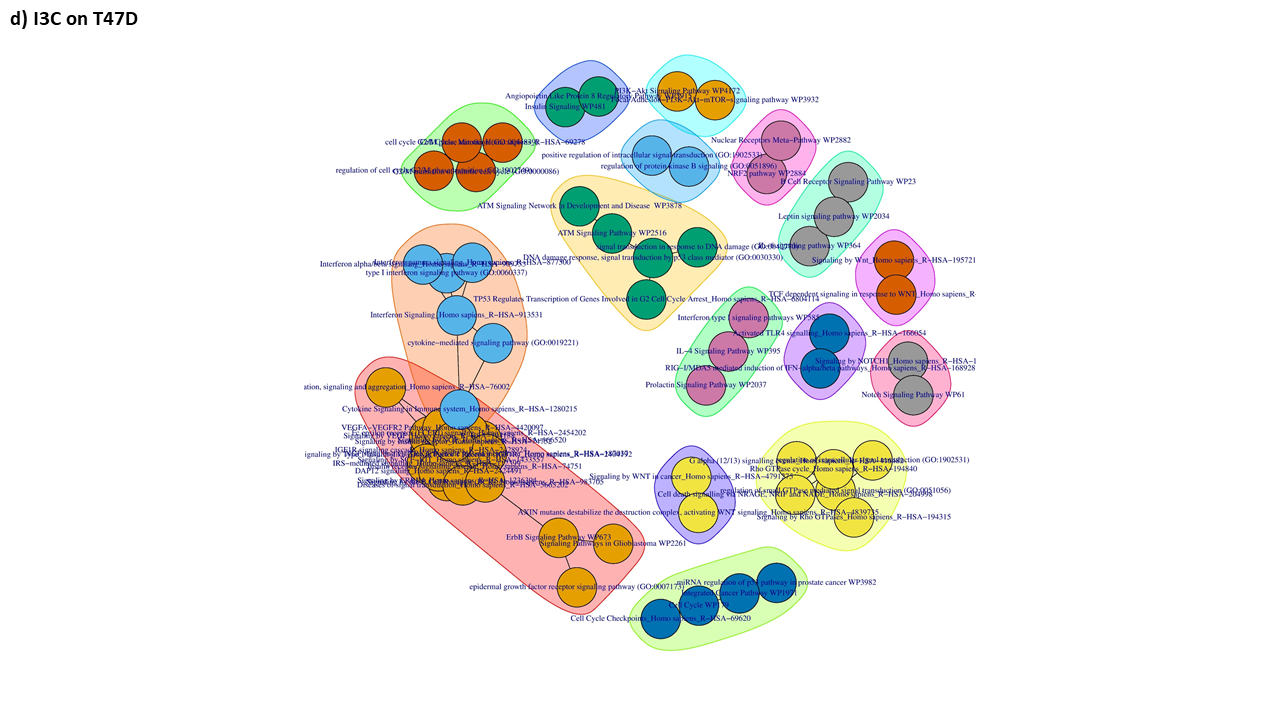

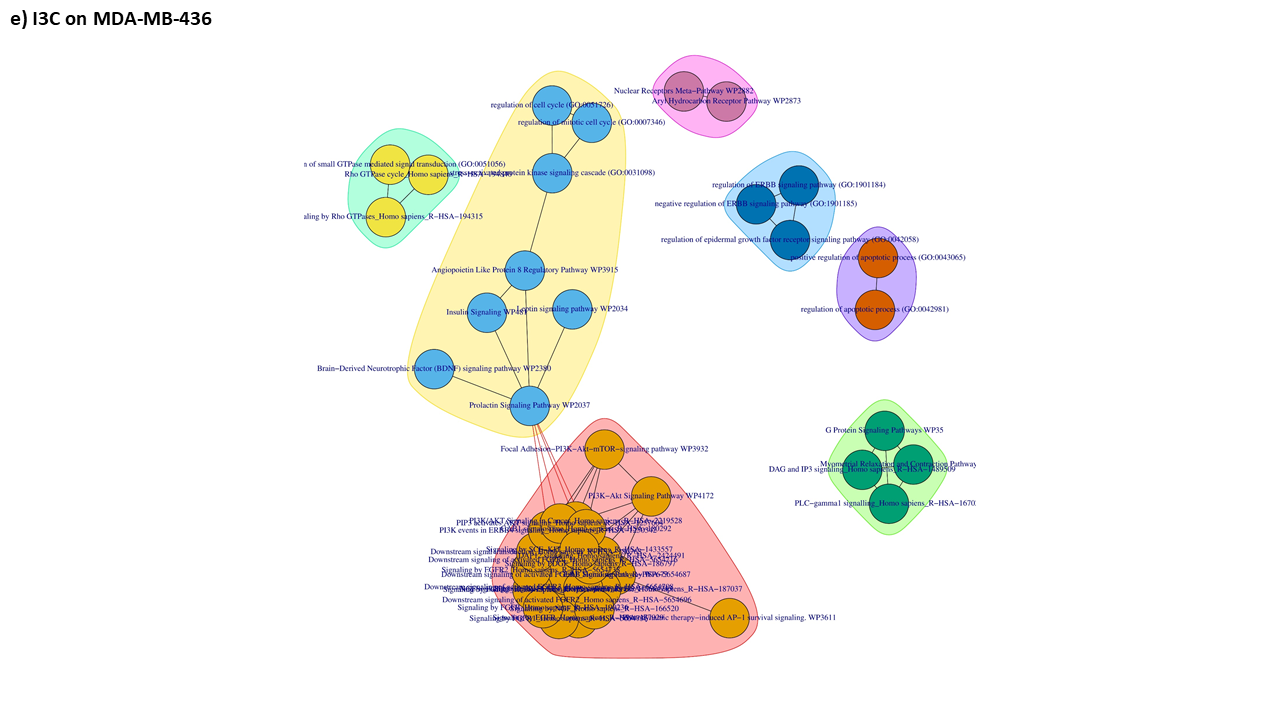

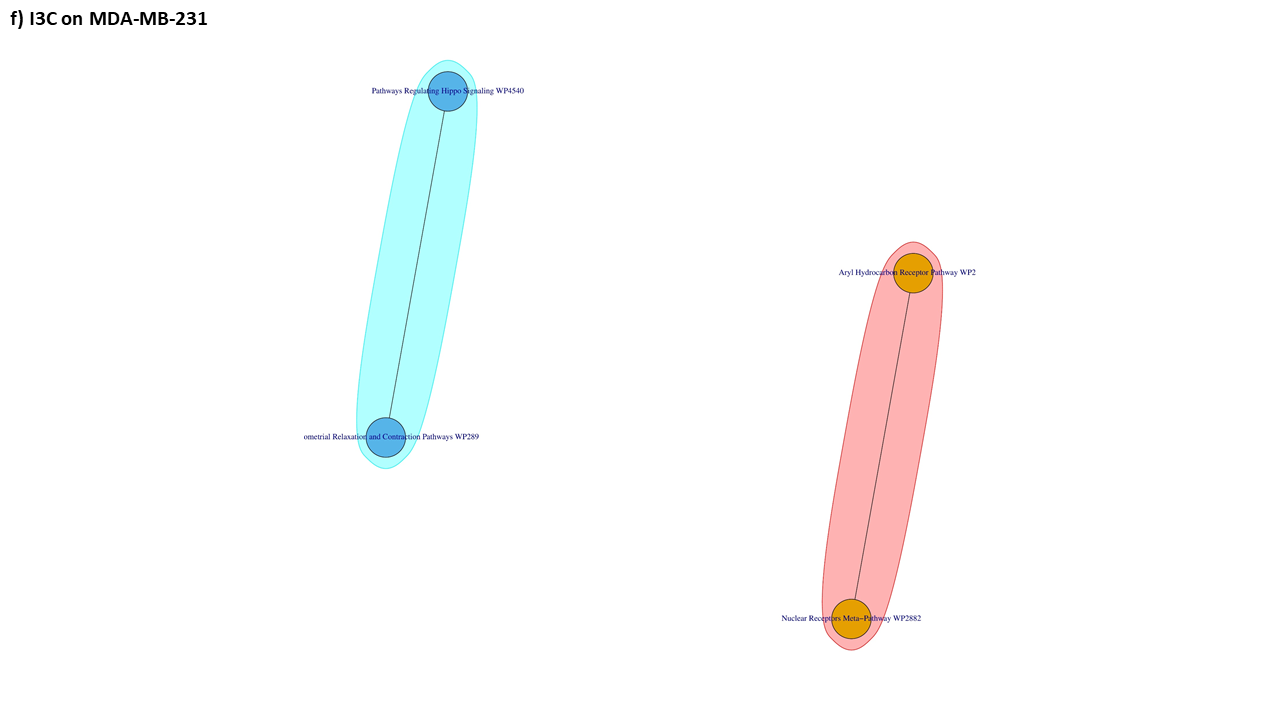

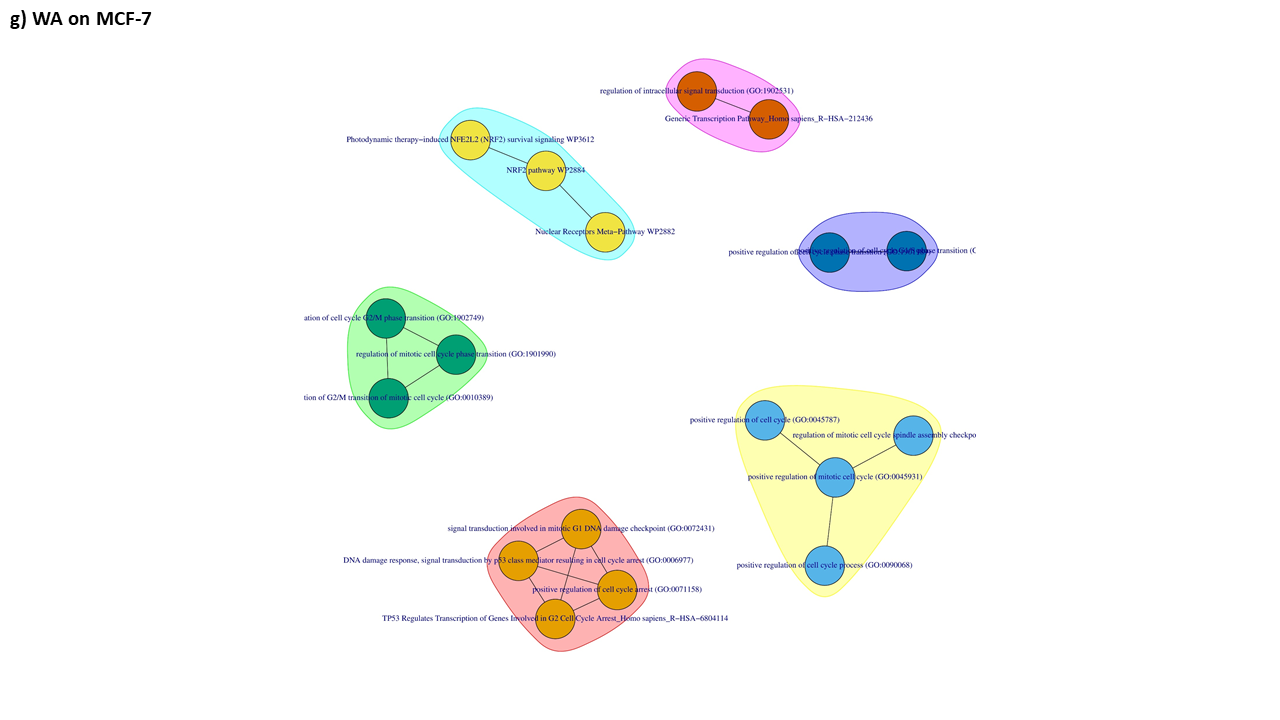

**Supplementary Figure 3: Pathway-pathway interaction networks based on shared enriched genes illustrating functional pathway cross-talk.** The differently coloured clusters illustrate highly related pathways terms based on intersecting pathways. a-g: represents networks of pathways targeted by CKI on MCF-7, I3C on MCF-7, I3C on MDA-MB-436, I3C on T47D, I3C on ZR751 and WA on MCF-7.
