## Supplementary Figure 1 for "A systems pharmacology approach to determine the mechanisms of action of pleiotropic natural products in breast cancer from transcriptome data"

Principal component analysis (PCA) results of transcriptome samples for each dataset illustrating the distribution of variance in the first two components considered for sample separation. PC1: principal component 1, PC2: principal component 2. (a) actein on MDA-MB-453, (b) CKI on MCF-7, (c) Indole-3-Carbinol on MCF-7, (d) Indole-3-Carbinol on MDA-MB-231, (e) Indole-3-Carbinol on MDA-MB-436, (f) Indole-3-Carbinol on T47D, (g) Indole-3-Carbinol on ZR751, (h) Withaferin A on MCF-7 and (i) Withaferin A on MDA-MB-231.

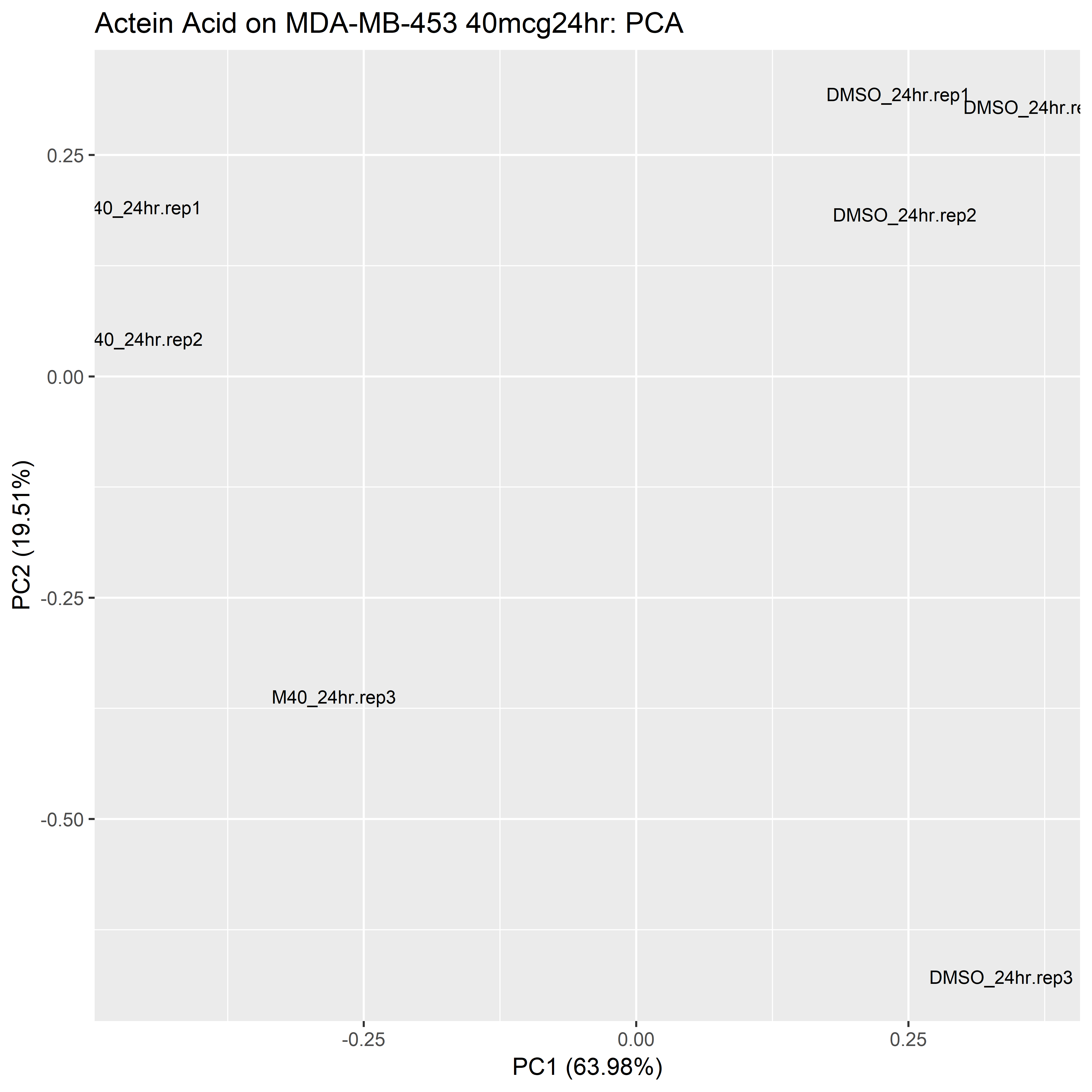

**(a)** actein on MDA-MB-453.

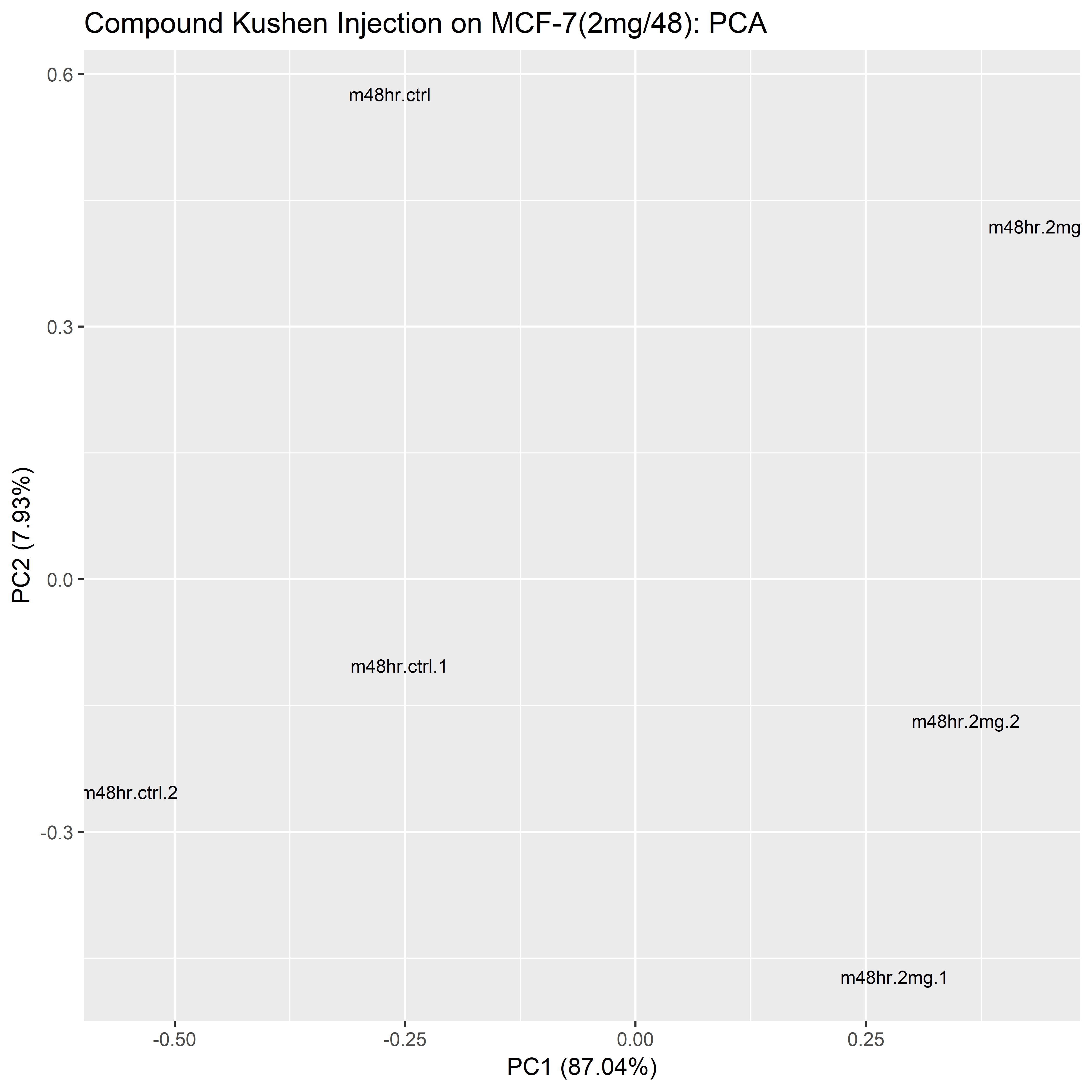

**(b)**  CKI on MCF-7.

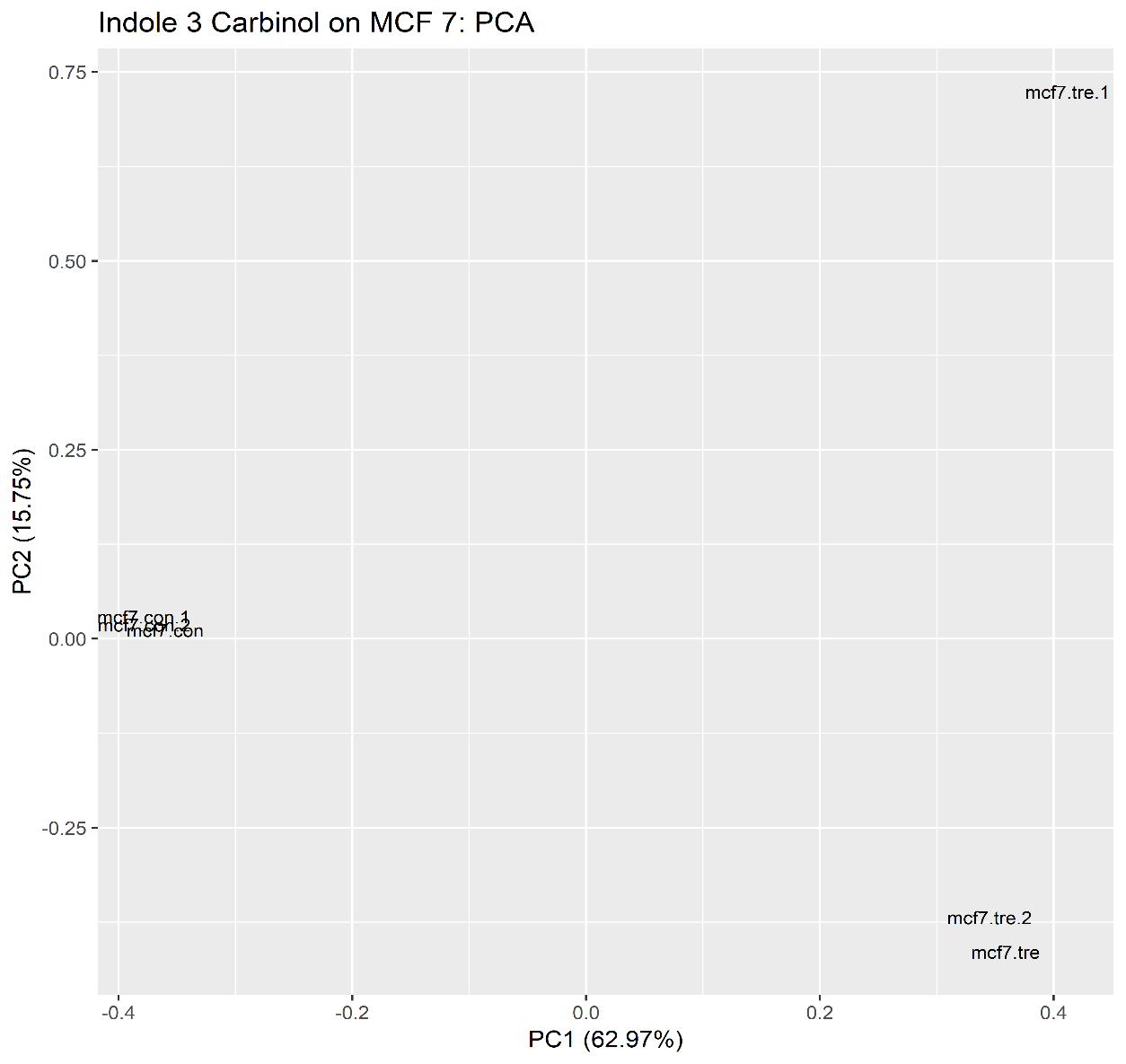

**(c)** Indole-3-Carbinol on MCF-7.

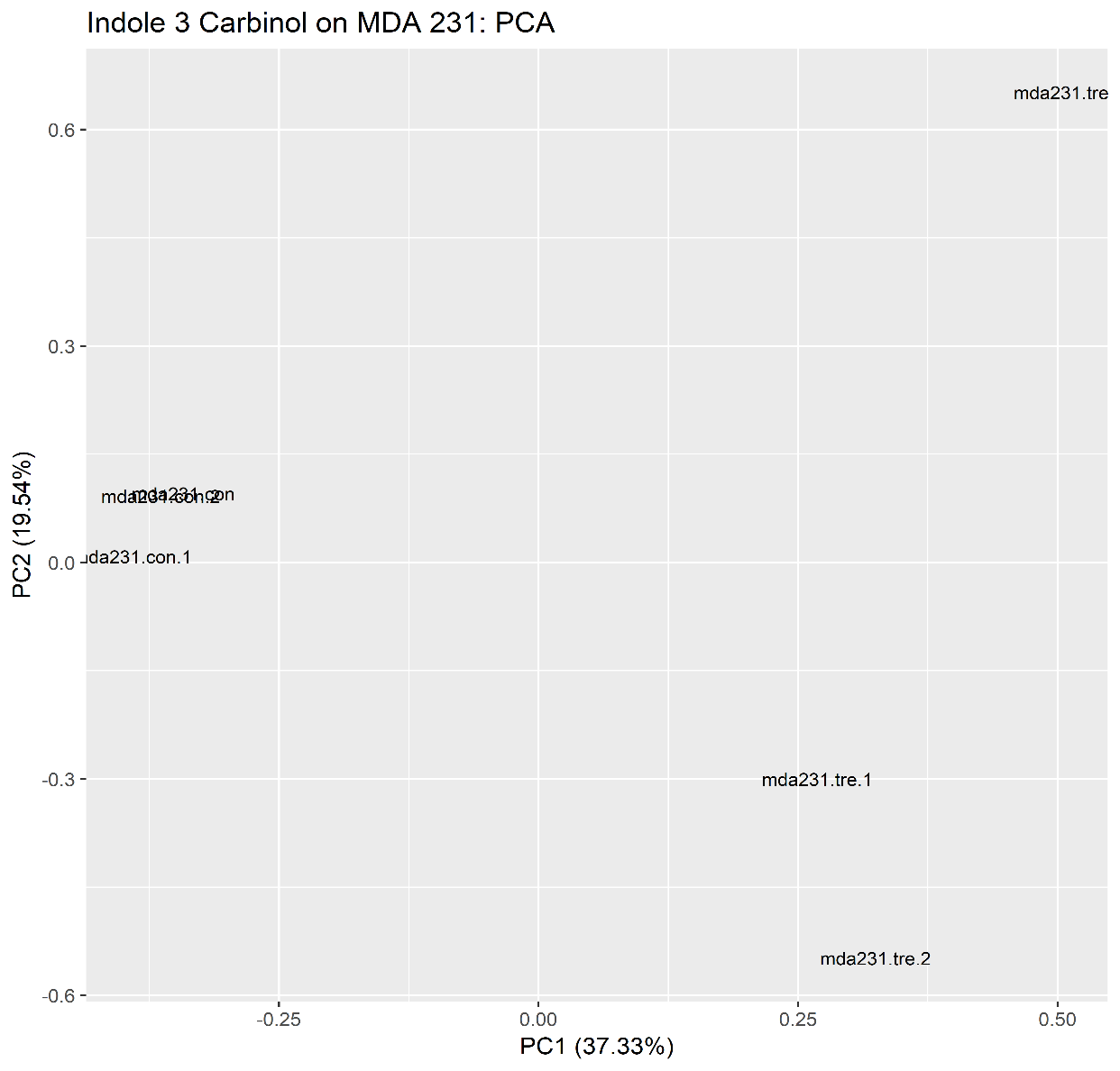

**(d)** Indole-3-Carbinol on MDA-MB-231.

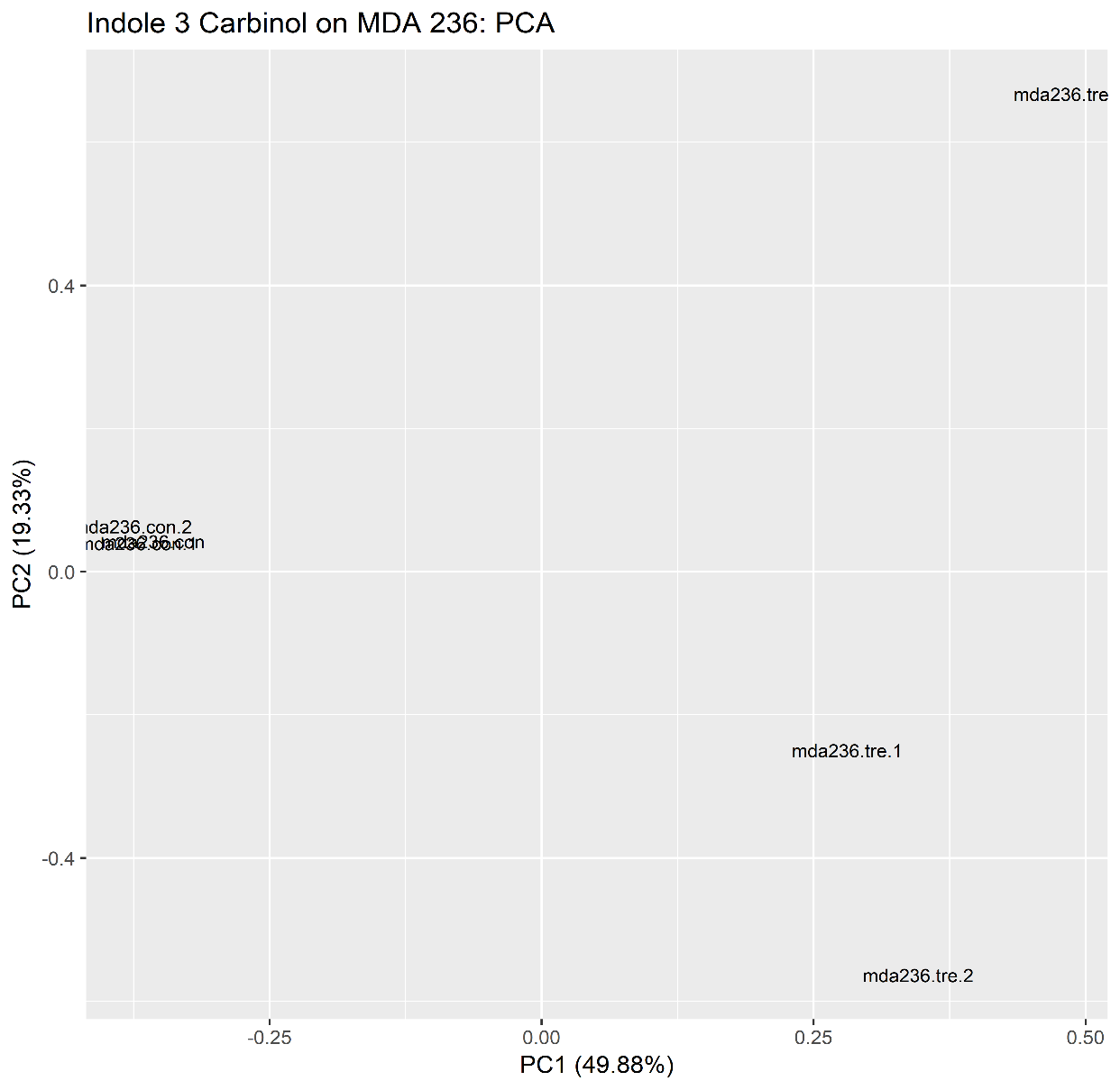

**(e)** Indole-3-Carbinol on MDA-MB-436.

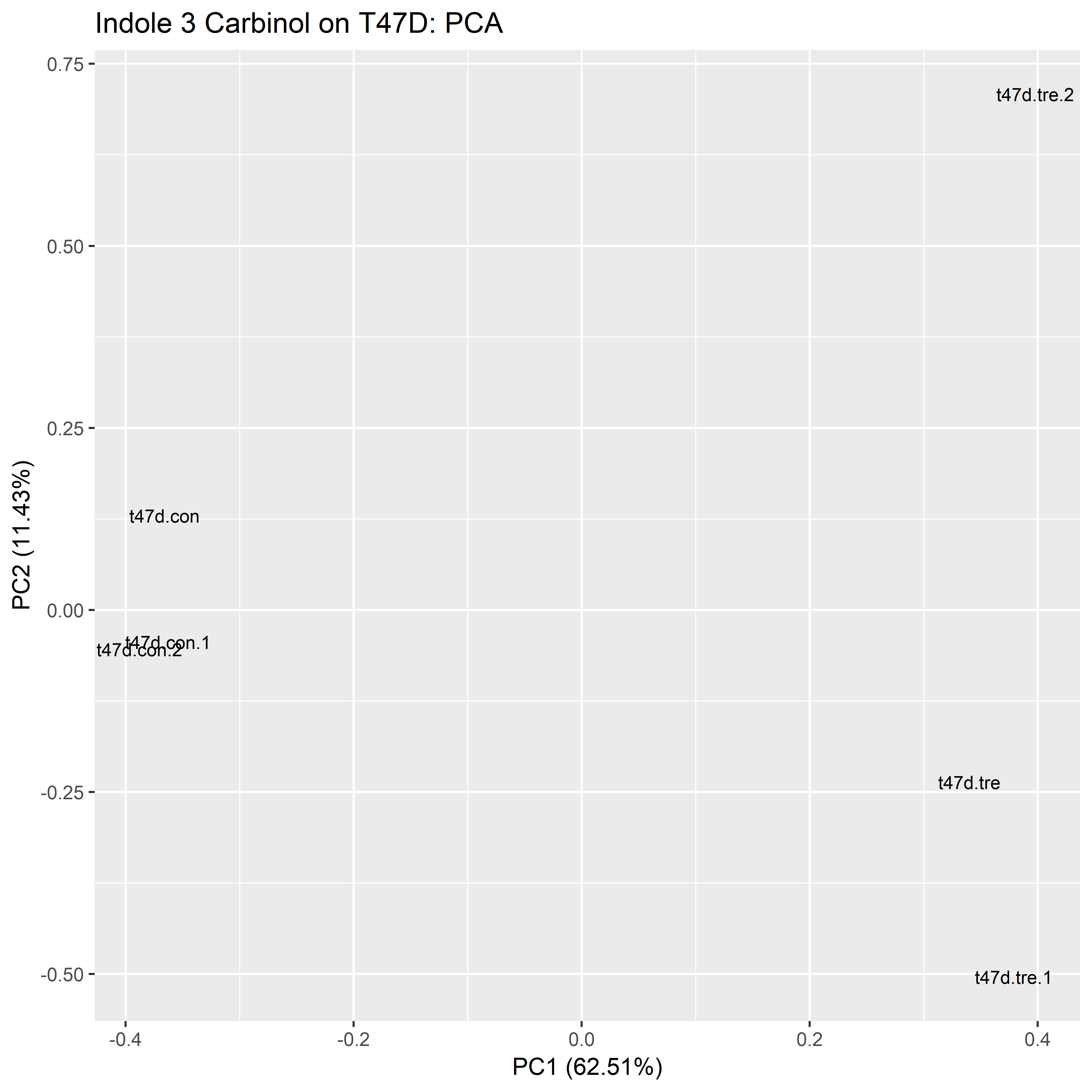

**(f)** Indole-3-Carbinol on T47D.

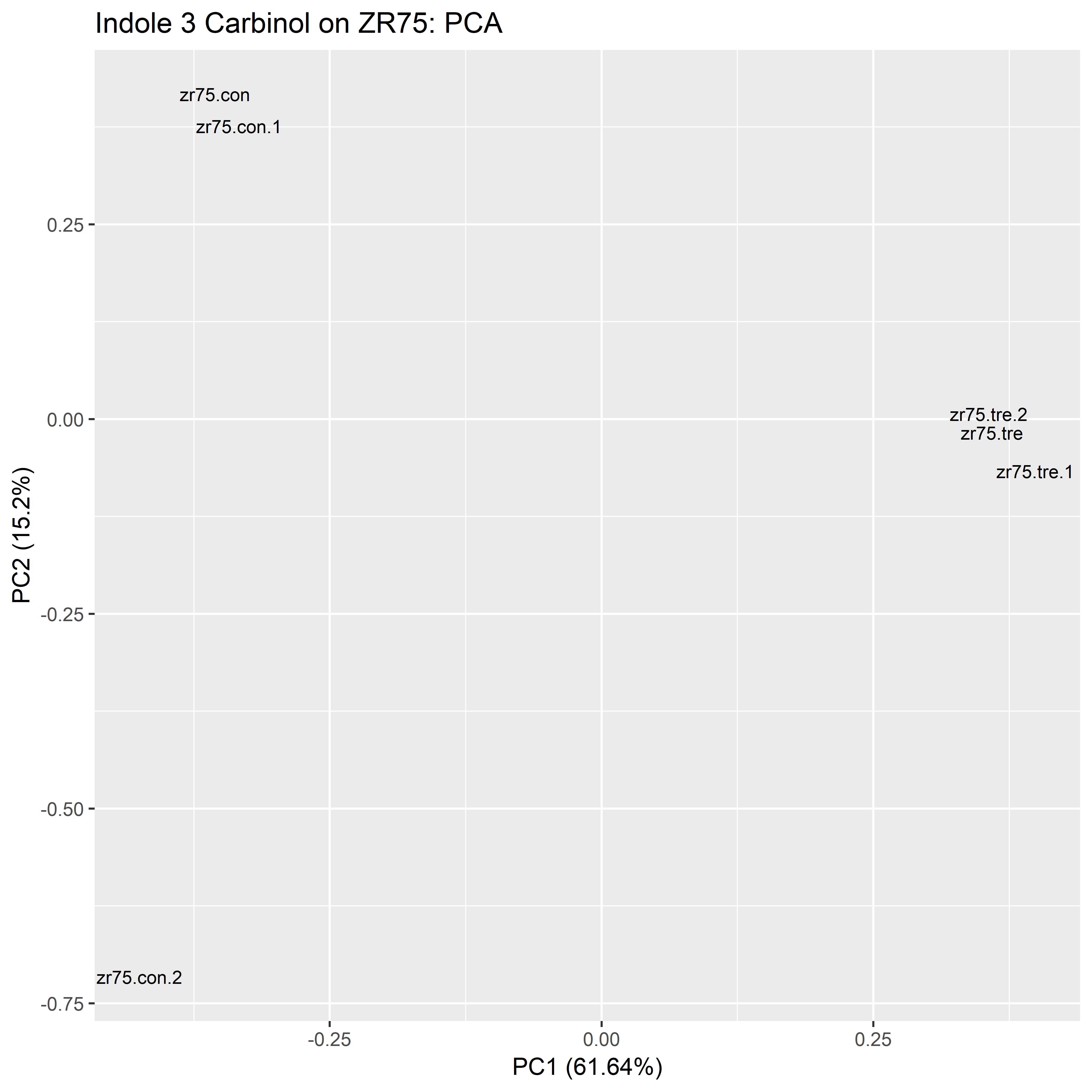

**(g)** Indole-3-Carbinol on ZR751.

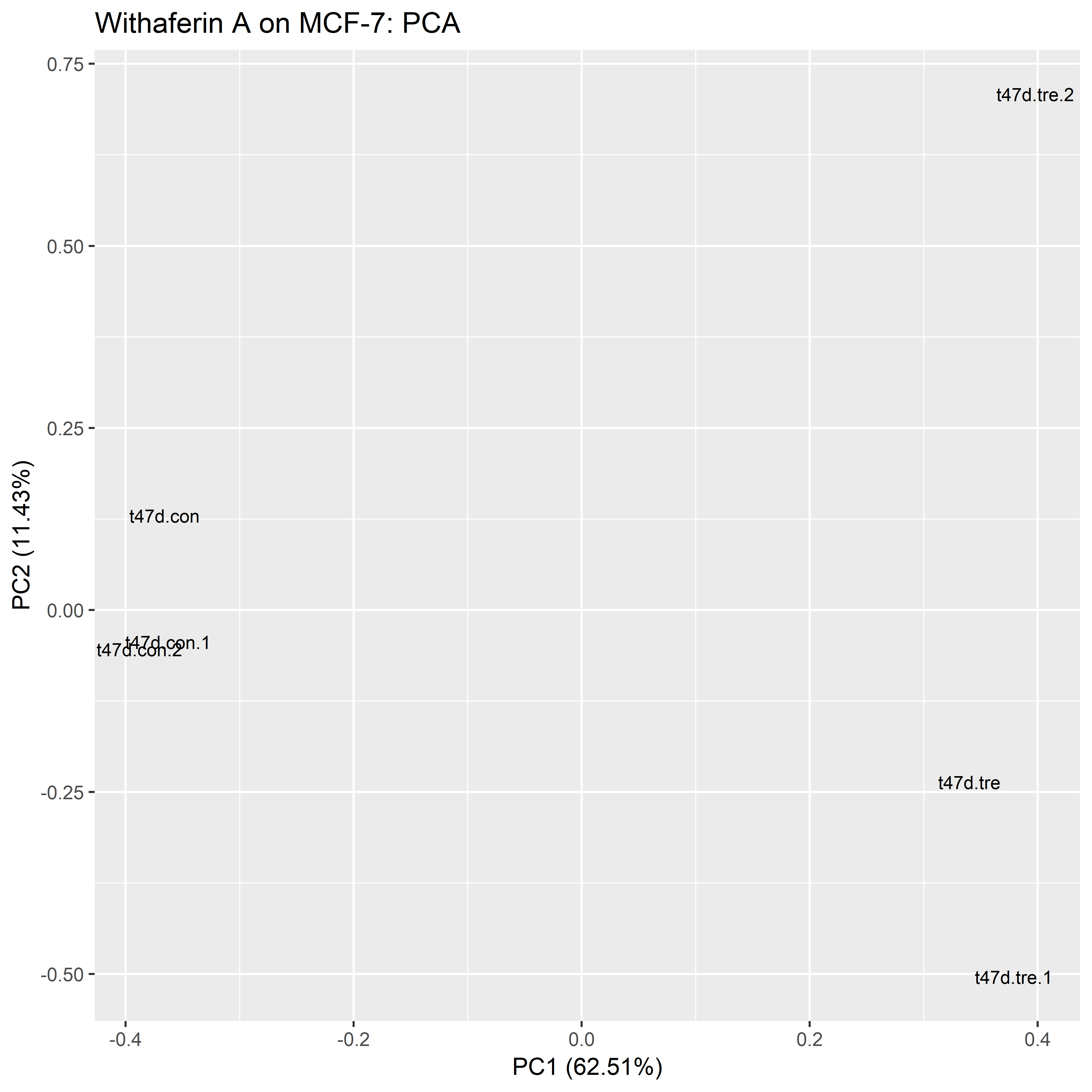

**(h)** Withaferin A on MCF-7.

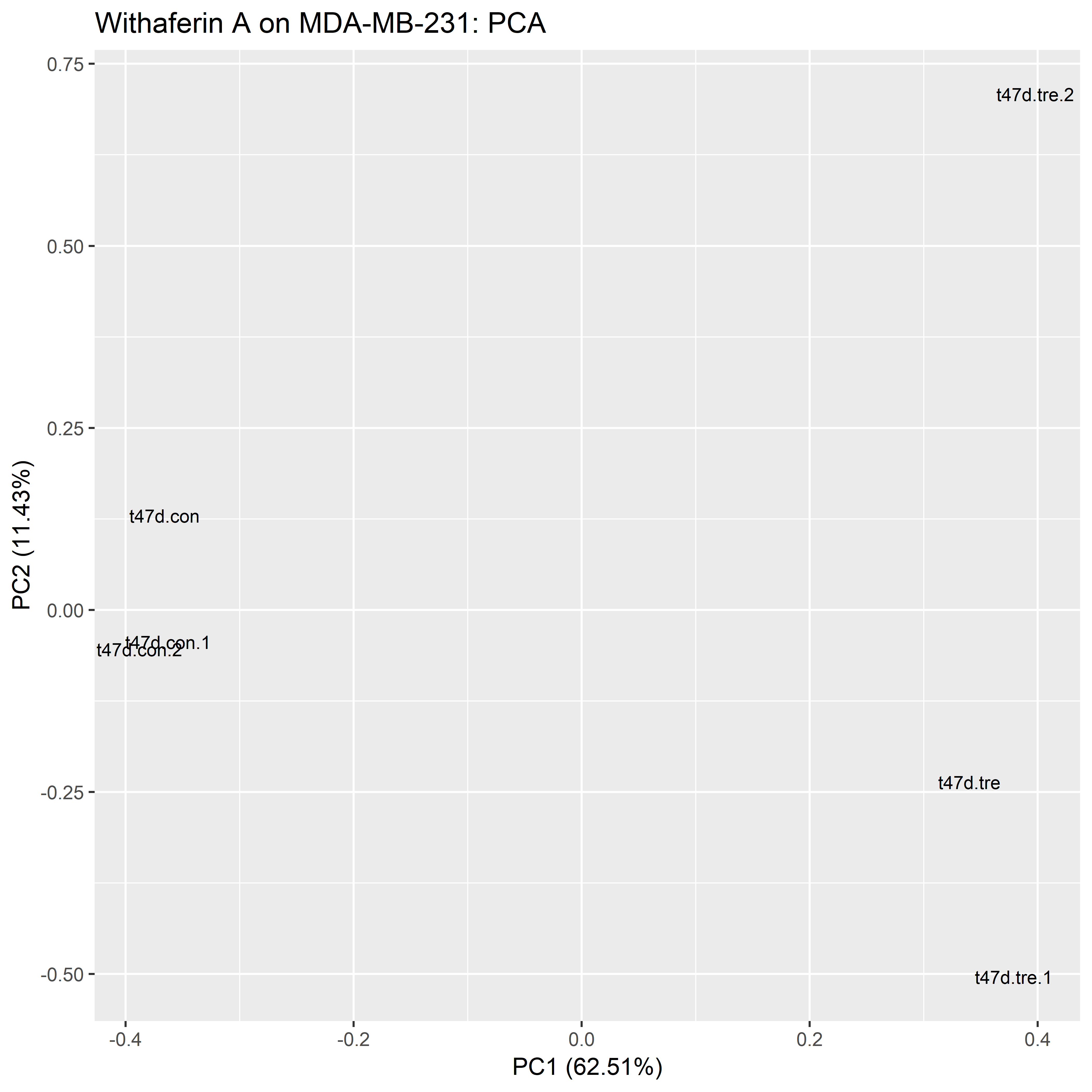

**(i)** Withaferin A on MDA-MB-231.
